# IDTrack: Time- and Namespace-Aware Identifier Harmonization for Reproducible Workflows

**DOI:** 10.64898/2026.02.05.703984

**Authors:** Kemal Inecik, Erkmen Erken, Fabian J. Theis

## Abstract

**Motivation:** Reproducibility in computational biology fails silently when gene identifiers drift beneath unchanged analysis code: the same frozen pipeline, rerun months later, yields different results not because biology evolved but because identifier semantics shifted with upstream annotation releases—a failure mode invisible to version control and containerization because the mapping layer itself constitutes an undeclared coordinate system whose time axis advances independently of downstream workflows. Gene identifiers occupy positions in a joint space of namespace, annotation release, genome assembly, and entity layer that evolves through retirements, merges, splits, and nomenclature reassignments, so atlas integration, retrospective reanalysis, and perturbation screens inherit temporal dependencies that existing utilities cannot surface: current mappers answer what an identifier resolves to now rather than under what declared contract the feature space was constructed.

**Results:** IDTrack reconceptualizes identifier harmonization as a time-indexed coordinate transformation by materializing annotation release history into a snapshot-bounded identifier graph and solving conversions through a time-traveling, contract-constrained pathfinder that pins release boundaries, assembly contexts, and ambiguity policies as explicit parameters rather than implicit endpoint state. This architecture surfaces reachability and ambiguity as interpretable outcome classes—unmapped, uniquely resolved, or ambiguously multi-target—enables atlas-scale harmonization with explicit collision handling, and records every mapping decision in a provenance ledger that transforms invisible preprocessing into citable methodological infrastructure whose coordinate choices can be inspected, compared, and reproduced rather than lost as ephemeral preprocessing.

**Availability and Implementation:** Code: https://github.com/theislab/idtrack; package: [pip install idtrack](https://pypi.org/project/idtrack/).

**Supplementary Information:** Supplementary material elaborates on architectural decisions and implementation details.

## Introduction

Gene identifiers appear stable—ENSG00000141510 looks like a permanent address, BRCA1 seems unambiguous—yet this apparent stability conceals a coordinate system that evolves with every Ensembl release, nomenclature revision, and cross-reference update, so that the same string can denote different biological entities depending on when the mapping was performed. Atlas integration (Figure 1) exposes this instability concretely: studies quantified against different Ensembl releases and identifier conventions scatter the same gene across non-matching labels, fragmenting equivalent features into redundant columns or losing them entirely to string-mismatch exclusion when pooled without explicit harmonization (1; 2; 3). The instability is a structural property of identifier semantics rather than a bug: each string occupies a position in a four-dimensional space of namespace, Ensembl release, genome assembly, and entity layer—coordinates that evolve through branching histories as biannual releases retire, merge, or split stable identifiers, as nomenclature authorities reassign symbols, and as cross-references expand through independent curation cycles (4; 5; 6; 7). This branching coordinate system induces structured ambiguity whose outcome classes must be surfaced explicitly: a single input identifier can have no reachable target (1→0), a unique target (1→1), or multiple plausible targets (1→n) under a fixed release boundary, while at dataset scale the same operator produces complementary outcomes—feature absence (0→1) and feature collisions (n→1)—whose handling determines what the integrated feature space denotes (Figure 2). Identifier harmonization is therefore not clerical preprocessing but the join operation that defines what a feature matrix means: every statistical test, integration model, and reported marker inherits the mapping layer’s coordinate assumptions whether or not they were declared, turning coordinate drift into an unreported experimental condition that rewrites the feature space between pipeline reruns under unchanged analysis code (8).

**Fig. 1.**
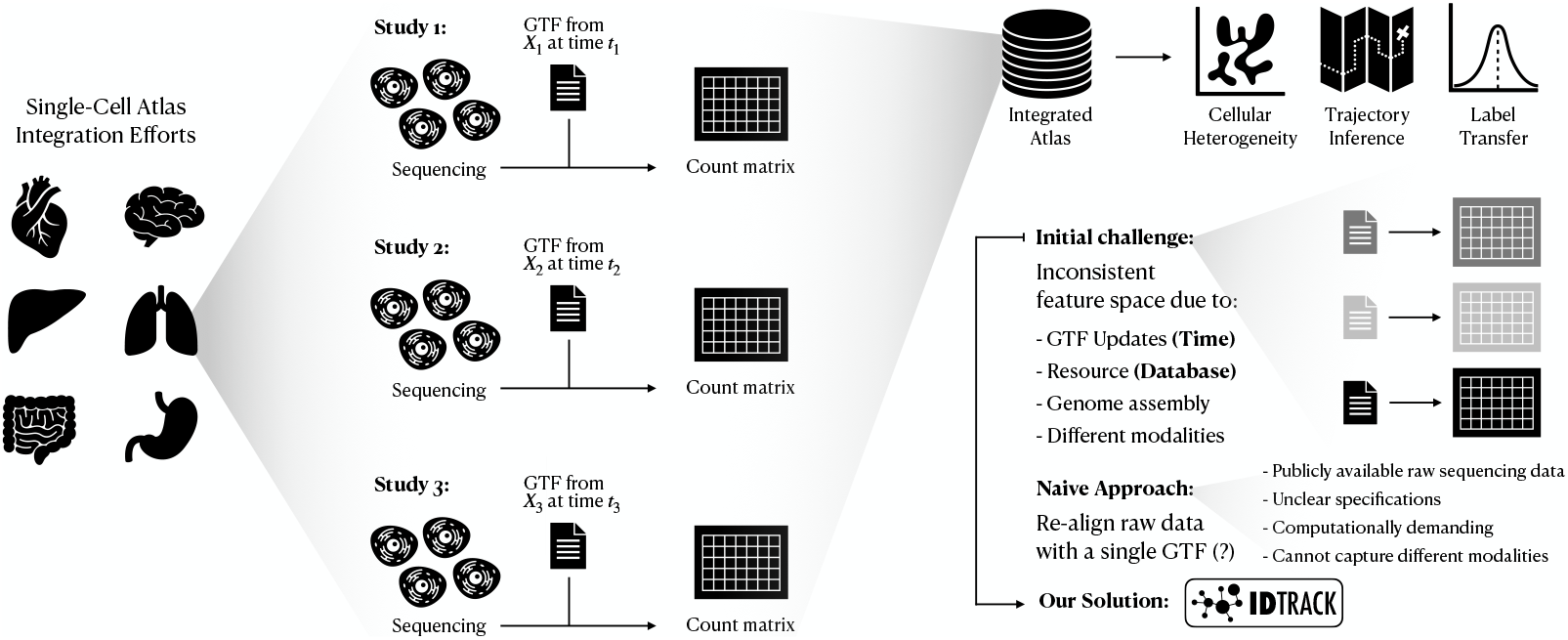
Datasets quantified against different annotation coordinates encode incompatible feature spaces whose reconciliation is the identifier harmonization problem. Concretely, studies processed at different times against different annotation snapshots, genome assemblies, and identifier namespaces produce count matrices whose column labels encode positions in a joint coordinate space of namespace, release, assembly, and entity layer—pooling such matrices without explicit coordinate reconciliation makes downstream integration, label transfer, and reporting depend on an implicit mapping layer that can drift under unchanged analysis code. This failure mode extends beyond atlas construction to affect variant annotation, perturbation screens, proteomics aggregation, knowledge-graph construction, and clinical traceability—any domain where identifier reuse mediates scientific claims and where the mapping layer’s coordinate assumptions propagate into downstream results. For settings where re-aligning raw reads to a single reference is infeasible, IDTrack restores a declared feature vocabulary by pinning a mapping contract that specifies snapshot boundary, assembly context, external evidence policy, and ambiguity strategy, producing an auditable harmonization ledger that transforms invisible preprocessing into a publishable methodological object.

**Fig. 2.**
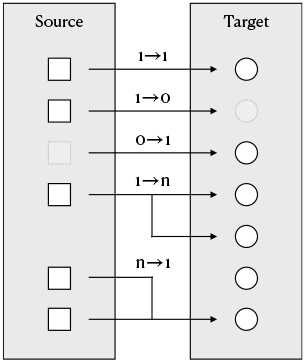
Drift-aware harmonization requires mapping outcome classes to be surfaced as first-class outputs rather than absorbed by downstream join logic. Because Ensembl histories branch through merges and splits and because cross-references across external namespaces overlap through synonym reuse and granularity mismatches, a single input identifier can have no reachable target (1→0), a unique target (1→1), or multiple plausible targets (1→n) under a fixed contract—outcomes that reflect coordinate-system properties rather than implementation accidents. Applying the same operator at dataset scale induces complementary join outcomes: feature absence (0→1) when target identifiers lack corresponding sources, and feature collisions (n→1) when multiple sources collapse onto one target—both requiring explicit policy decisions whose handling must be reported as part of the mapping contract.

This integration failure is not an isolated case but the visible symptom of a pervasive condition: coordinate drift propagates silently through every analysis that reuses identifiers across temporal or database boundaries, producing results that complete without errors yet encode assumptions that cannot be audited after the fact. In expression analysis, a reported differential marker can appear because two genes merged between annotation releases rather than because of biology— signal that fragments or vanishes when the same analysis runs against updated coordinates, potentially invalidating published findings without any error to flag the discrepancy (4). In proteomics, peptide-to-gene aggregation silently reroutes when isoform definitions change: the same mass-spectrometry signal maps to different genes depending on which annotation snapshot was used, propagating inconsistencies into multi-omic integration where transcript and protein layers must align (9; 10). In functional genomics, perturbation screens lose traceability when gene symbols are reassigned after library design—the reported hit no longer points to the gene that was actually perturbed, yet the mismatch surfaces only if someone manually audits the historical mapping (11). Across all these domains, a pipeline rerun under identical code can yield different results because the “latest” API endpoint moved—no error, no warning, just a silently rewritten feature vocabulary that invalidates comparisons to earlier runs and makes long-horizon reanalysis unreliable. The scientific gap is therefore not that any single domain is technically difficult but that identifier harmonization mediates reuse across all of them, yet the ecosystem treats mapping as undocumented preprocessing whose coordinate choices are rarely surfaced as publishable, auditable contract objects (see Suppl. Sections 3 and 4).

The bioinformatics ecosystem offers no shortage of mapping utilities—*BioMart* for schema-level queries, *MyGene.info* and *gget* for high-level aggregation, *BridgeDb* for backend abstraction, *Bioconductor ‘s AnnotationDbi* for static annotation packages—each indispensable for point-in-time lookup and exploratory annotation, yet each answering a question orthogonal to the drift problem: “what does this identifier resolve to now” rather than “under what declared contract was this feature space constructed” (12; 13; 14; 15; 16; 17). The consequence is that harmonization remains an unprincipled, user-dependent preprocessing step whose coordinate choices are encoded in ad hoc scripts, undocumented endpoint calls, and implicit join logic that cannot be inspected, reproduced, or published as a methodological object—each analyst reinventing resolution heuristics whose assumptions are neither declared nor portable across compute environments. What the ecosystem lacks is not another mapping endpoint but a contract-first framework that elevates harmonization from invisible preprocessing into principled methodology: one in which release boundaries, assembly contexts, namespace policies, and ambiguity strategies constitute an explicit coordinate system whose declarations can be archived alongside data and code, whose traversal semantics are mechanically auditable, and whose outputs encode outcome classes as interpretable properties of the declared contract rather than as implementation accidents absorbed by downstream analysis (see Suppl. Section 4 for an extended capability taxonomy).

IDTrack realizes this contract-first framework by treating identifier harmonization as a coordinate transformation whose every parameter is declared, frozen, and publishable: the tool embeds Ensembl release history as a navigable time axis, materializes a snapshot-bounded and assembly-aware identifier graph as a portable artifact, and resolves conversions through a pathfinder whose admissibility constraints, traversal semantics, and outcome classes derive entirely from the declared contract rather than from implicit endpoint state or ad hoc join logic. The scientific payoff is that harmonization becomes a citable methodological object: the contract specification travels with data and code, reviewers and collaborators can inspect the coordinate choices that produced a feature space, reanalysis under identical declarations yields identical outputs regardless of when or where computation occurs, and outcome classes surface as interpretable properties of the declared coordinate system rather than as silent artifacts of undocumented preprocessing. This architecture targets the practical reality that read-level reprocessing is often infeasible—missing raw files, unknown aligner configurations, prohibitive compute costs, data governance restrictions—yet drift-sensitive reuse spans the full application surface from atlas harmonization and long-horizon reanalysis to legacy gene-list rescue, multisite integration, and cross-database interoperability (see Suppl. Section 3 for an extended application taxonomy). We demonstrate that branching *Ensembl* histories and bounded external bridges induce interpretable outcome profiles whose structure reflects coordinate-system properties, and show atlas-scale harmonization that preserves a declared feature vocabulary across heterogeneous release contexts under a single published contract whose audit trail remains mechanically reproducible.

## Methods

We formalize IDTrack as a build-once, query-many mapping operator on a snapshot-bounded identifier multigraph indexed by a declared contract, so identifier conversion becomes a reproducible coordinate transform whose semantics are stable under reruns and whose failure/ambiguity modes are returned as first-class outputs rather than absorbed as implicit preprocessing. We (i) define the mapping contract 𝒞 and its materialization 𝒢_𝒞_, (ii) formalize contract-admissible path witnesses and a lexicographic selector that surfaces 1→0/1→1/1→n outcomes with auditable evidence, and (iii) lift the same operator to dataset-scale feature harmonization with explicit collision handling and a provenance ledger (implementation-level defaults and pseudo-specs: Suppl. Sections 6.1–6.14 and 7.1–7.2).

This workflow is parameterized by a mapping contract 𝒞 = (org, ℛ_𝒞_, a^⋆^, Λ_𝒞_, Θ_𝒞_) that fixes an inclusive release window ℛ_𝒞_ = [r_min_, r_max_], a primary assembly a^⋆^ with an organism-scoped assembly-priority order, an opt-in external allowlist Λ_𝒞_, bounded bridging parameters Θ_𝒞_ (synonym depth limits, external jump budgets, and hyperconnective-hub filtering), and a deterministic ambiguity policy (`strategy=“best”` vs. `strategy=“all”`). Each conversion claim then instantiates 𝒞 by choosing `to_release`∈ ℛ_𝒞_ and optionally `from release` and `final database`, which together fix admissibility and the reported codomain of Map_𝒞_and therefore must be disclosed alongside 𝒞. Interpreting each identifier string as a partial coordinate in namespace × release × assembly × layer makes conversion a coordinate completion followed by projection, and it makes Figure 1 a statement that feature matrices are comparable only when their coordinate systems are declared rather than assumed. The contract makes “current” illegal by construction, because admissibility and scoring are evaluated only on releases inside ℛ_𝒞_ rather than on a moving endpoint.

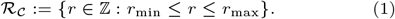

To make the partial-coordinate view executable, raw query strings are resolved by a deterministic canonicalization map *c(·)* implemented as a bounded cascade (exact match; case-insensitive lookup via an injective lowercase dictionary guaranteed by build-time merging of case-colliding external nodes; regex stripping of terminal version-like suffixes; dash/underscore permutations), and the same cascade is retried under an explicit synonym-prefix namespace so normalization remains auditable rather than analysis-specific. We write *q := c(q_raw_)* for the canonical node label when resolution succeeds (otherwise c(q_raw_) = ∅). Given 𝒞, IDTrack materializes these coordinates as a frozen identifier graph 𝒢_𝒞_ serialized together with the allowlist and cached source tables, so reruns differ only when the declared contract or archived artifacts differ. All downstream tasks reduce to bounded operations on this frozen substrate (Figure S1), including source inference for unlabeled lists, contract-constrained time travel into `to_release` with optional cross-namespace projection, and dataset-scale harmonization with an explicit mapping ledger rather than implicit join logic. For a canonical query node *q*, IDTrack returns a set of targets *Map*_𝒞_(*q*) whose cardinality induces the outcome class (1→0, 1→1, 1→n), and when audit evidence is requested each returned target is paired with an admissible path witness that certifies admissibility under 𝒞.

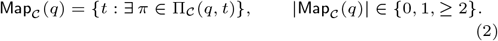

Here Π_𝒞_(q, t) is the set of contract-admissible path witnesses defined below, so reachability can be certified by an explicit witness rather than assumed from endpoint trust. When a single representative is requested (`strategy=“best”`), IDTrack returns a deterministic selector BestMap_𝒞_(q) ∈ Map_𝒞_(q) defined by a lexicographic objective over path costs and final-conversion confidence, whereas `strategy=“all”` returns all scored targets so ambiguity remains explicit rather than being coerced away.

To make 𝒞 operational, graph construction and downstream queries are snapshot-first rather than endpoint-first, because once 𝒢_𝒞_ is built under the declared contract, conversions are solved by bounded traversal and deterministic scoring on a frozen substrate rather than by repeated remote lookup with implicit defaults. Because all admissibility predicates and scoring terms are defined on edges of 𝒢_𝒞_, snapshot construction fixes the feasible search space and is therefore a methodological choice rather than an implementation detail.

### Snapshot-bounded, assembly-aware graph construction

Figure 3 fixes the object that the contract materializes: a snapshot-bounded, assembly-aware identifier multigraph in which Ensembl stable-ID history provides the time axis and allowlisted cross-references provide bounded lateral evidence, so time travel and cross-namespace conversion are solved by a single admissibility predicate on one substrate and reachability/ambiguity become functions of declared bounds (ℛ_𝒞_, a^⋆^, Λ_𝒞_) rather than accidental properties of a queried endpoint.

**Fig. 3.**
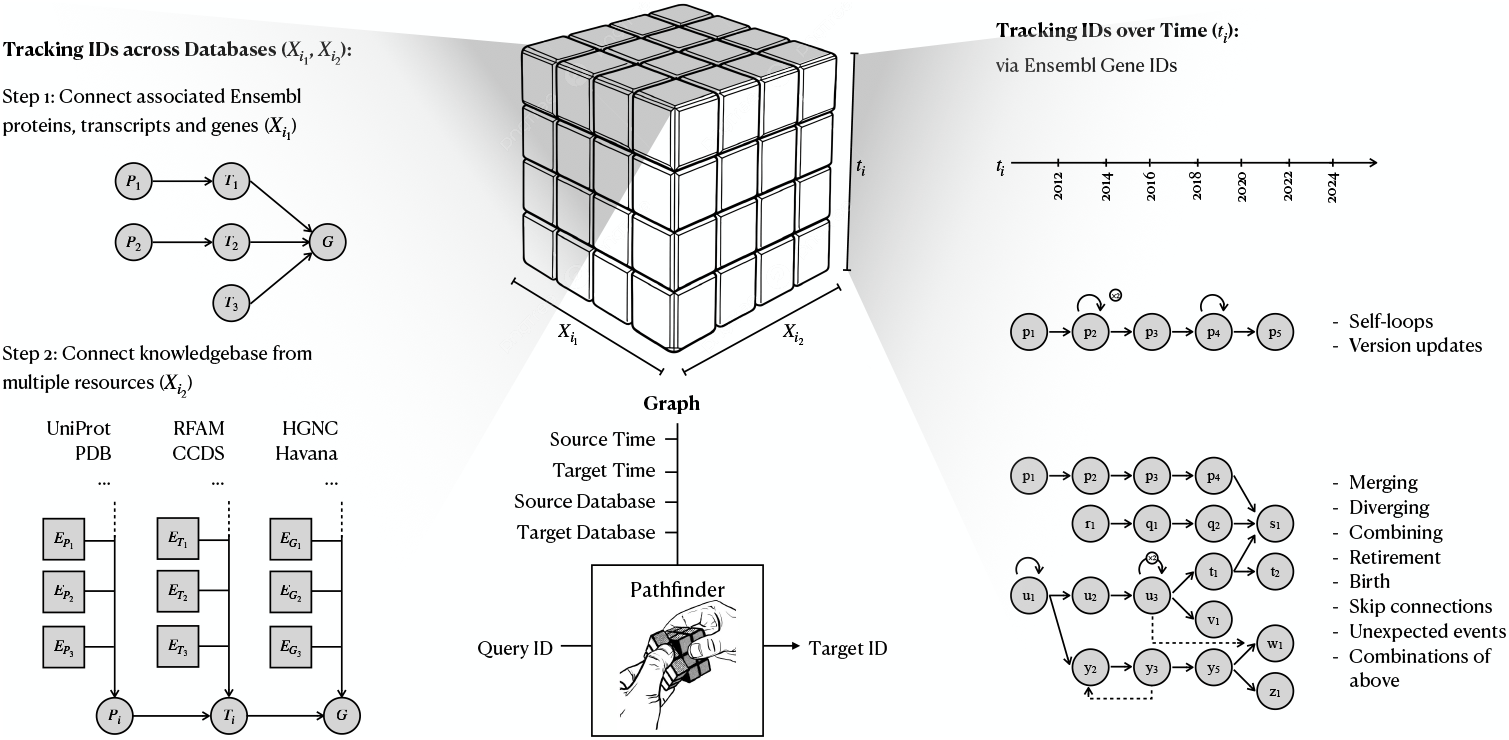
The IDTrack coordinate system: a snapshot graph coupling Ensembl time travel with cross-namespace edges. Within each release, IDTrack links Ensembl entity layers and allowlisted external databases into a joint graph, while across releases the stable-ID history defines a time axis with structured events (version updates, merges/splits, retirements/births, and rare non-local transitions) that make mapping inherently set-valued rather than functional. By embedding these relations in a snapshot-bounded substrate, a conversion becomes a contract-constrained path query that time-travels an input to a target release and optionally projects it into a target namespace, with returned targets accompanied by outcome semantics and, when requested, auditable traversal paths.

IDTrack constructs a directed multigraph 𝒢_𝒞_ = (V, E_H_ ∪ E_X_ ∪ E_L_) whose nodes represent identifiers typed by *Ensembl* entity layer and assembly context (including versionless base identifiers as explicit nodes) as well as external namespaces, and whose multiedges preserve distinct evidence channels without premature conflation. History edges E_H_ are extracted from the *Ensembl* Core stable-ID lineage tables (`stable_id_event` joined with `mapping_session`) within ℛ_𝒞_ and are annotated by release endpoints and the upstream combined mapping score, so when multiple admissible histories exist the pathfinder can prefer higher-evidence transitions under a declared scoring rule rather than by arbitrary tie-breaking. Cross-reference edges E_X_ are extracted from the xref join (`object_xref, xref, external_db, identity_xref, external_synonym`), filtered at build time by Λ_𝒞_, and recorded as external→*Ensembl* attachments with explicit release and assembly validity, with synonym display labels encoded as a separate namespace via a dedicated prefix to prevent conflating accessions with free-text aliases. Cross-layer edges E_L_ connect entity layers along biological containment (translation→transcript→gene) using *Ensembl*’s `relationcurrent` table, allowing traversal to carry evidence across layers while constraining reported targets to the requested layer and assembly context. Every external/cross-layer edge is annotated with set-valued validity metadata, and we cache the release support ℛ(e) and assembly scope 𝒜(e) so that admissibility checks reduce to set membership tests during traversal rather than being deferred to a post hoc filter of a flattened join. With an allowlist predicate allow_𝒞_ (enforced mechanically at build time) and cached membership tests, edge admissibility becomes a local Boolean predicate evaluated inside the pathfinder.

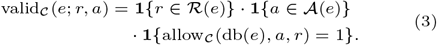

Because assemblies are part of the coordinate system, construction can retain multiple assemblies inside the snapshot window while enforcing a declared primary-assembly view for outputs, preventing build-scoped cross-references from being silently merged into a scientifically ambiguous feature space. To keep traversal tractable, 𝒢_𝒞_ is accompanied by deterministic derived summaries such as reverse views, combined-edge dictionaries keyed by (database, assembly, release), compact node activity ranges, origin-trio caches, and hyperconnective-hub filters, each computed once from the frozen substrate so repeated query-time checks become dictionary/set membership tests rather than repeated neighbor scans that would otherwise dominate path search.

### Pathfinder conversions with explicit ambiguity and audit paths

Given the canonical query node q, conversions are computed as a staged, contract-bounded enumeration of admissible path witnesses on 𝒢_𝒞_, where temporal orientation is inferred from cached node activity ranges when `from_release` is unspecified and the walk time-travels to `to_release` before optionally projecting into a requested external namespace, after which the reachable target set is returned with explicit (1→0)/(1→1)/(1→n) outcome semantics (Figure 2). An admissible conversion route can be represented by a path witness that interleaves history edges (timestamped by release endpoints) with external edges (validated by membership in cached release/assembly sets), so admissibility is checkable from the witness rather than assumed by an endpoint.

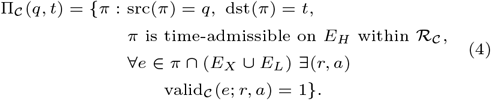

Here time-admissible means that along E_H_ the walk follows a monotone sequence of history events whose releases lie in ℛ_𝒞_ and that the reached node is active at `to_release` under its cached activity range. Search is staged with progressive relaxation, because the relevant question under drift is not only “what is reachable” but also “under what minimal relaxation of declared bounds does reachability appear,” which makes failure and ambiguity interpretable as coordinate-system properties rather than as unstable search behavior. IDTrack first attempts backbone-only traversal within the declared window, then permits bounded external “beam-up” steps by enumerating synonym neighbourhoods under increasing limits on synonym depth and external jump budget, and if no path exists it repeats the same schedule under a relaxed multi-transition mode that allows the inferred starting release to shift on external nodes while keeping ℛ_𝒞_ and Λ_𝒞_ fixed. Equivalently, the staged search induces a nested family of feasible sets 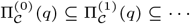 in which each stage expands admissibility only by relaxing an explicit bound, making “no path” a contract-level statement rather than a runtime accident. Candidate paths are ranked by a deterministic lexicographic objective that treats assembly downgrades and external beam-ups as first-order penalties and uses *Ensembl* history evidence only after these constraints, so the algorithm prefers minimal-policy explanations over opportunistic transitive closures while remaining stable under reruns.

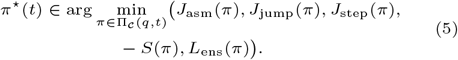

Here J_asm_ counts required assembly downgrades given per-edge feasible assembly sets, J_jump_ counts external beam-ups, J_step_ counts traversed external edges, S(ω) is the reduced history-edge evidence score, and L_ens_ counts *Ensembl*-history steps, making canonical selection a deterministic policy statement rather than an implementation artifact. If a target external namespace is requested, the final projection step performs a bounded synonym search from the reached *Ensembl* gene into the requested database constrained to the target release (confidence 0), and if no such attachment exists it falls back to a release-agnostic synonym search whose last-node validity is still checked at the target release (confidence 1) or, optionally, returns the *Ensembl* gene itself as an explicit no-target fallback (confidence ∞). When `strategy=“best”`, a second lexicographic filter selects a single globally best (*Ensembl* gene, target ID) pair using final-conversion confidence and assembly-priority metrics with deterministic tie-breakers (prefer identity with the query when possible, otherwise prefer the synonym-richer *Ensembl* node), whereas `strategy=“all”` returns all scored targets so downstream analyses can propagate uncertainty explicitly. Let N_π_ := Σt | Π_𝒞_(q, t)| and 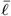 be the number and mean length of admissible witnesses enumerated under the staged bounds (synonym depth, jump budget, hub filtering) within ℛ_𝒞_, then per-query time is 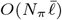 and space is 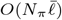, so algorithmic complexity becomes a declared property of 𝒞 rather than an opaque property of a queried endpoint. When enabled, audit payloads return the concrete traversal path that justifies each outcome, including the external hop segment when present, making conversions inspectable as evidence sequences that can be archived, quality-controlled, and defended as part of Methods metadata rather than rediscovered through debugging.

### From conversions to harmonized feature spaces

Having defined the single-identifier mapping relation, IDTrack treats harmonization as the application of the same contract-defined operator to entire feature vocabularies, because integration and model training implicitly assume a shared coordinate system even when per-dataset preprocessing choices differ across time, assembly, and namespace. For dataset i with feature set ℱ_i_, we compute mapped candidates 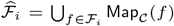 (optionally after best-selection) and record for each f whether the outcome is 1→0, 1→1, or 1→n, because these outcome semantics are the only stable diagnostics available under drift when a ground-truth mapping is ill-defined.

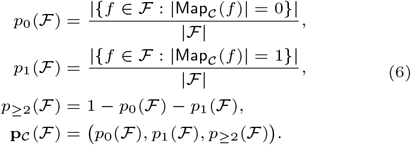

Across datasets we define a target feature space either as a union 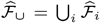 or an intersection 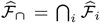, and we quantify how this set-level choice redistributes missingness and ambiguity under the same contract. To prevent many-to-one collapses (n→1) from being silently absorbed by sparse-matrix join logic, the harmonizer detects collisions where distinct source identifiers map to the same target identifier, resolves them deterministically under an explicit preference rule, and records discarded sources together with the contract identity and per-dataset matchings in a mapping ledger that can be archived as part of Methods metadata. When source metadata are missing, source inference proposes plausible origin coordinates by aggregating the cached node-trio contexts (db, assembly, release) observed across the identifier list, and restricted-network operation supports building the same snapshot artifacts on HPC systems where compute nodes lack direct internet access.

### Availability and implementation

IDTrack is available as a Python package ([pip install idtrack](https://pypi.org/project/idtrack/)), with documentation at https://idtrack.readthedocs.io/ and source code at https://github.com/theislab/idtrack.

## Results

### Release drift becomes measurable once time is treated as a first-class coordinate rather than an implicit default

The structure of identifier drift becomes empirically accessible only when raw data are held fixed while annotation coordinates are systematically varied, a controlled setting that transforms instability from an anecdotal complaint into a measurable property of the release axis whose outcome profiles can be attributed unambiguously to coordinate-system evolution rather than to biological heterogeneity or processing variation. We operationalize this principle by treating the *Cell Ranger* feature vocabulary as part of the data-generating process: quantifying the same *pbmc_1k_v3* FASTQs (18) across *Ensembl* references spanning releases 82–114 on GRCh38, then time-traveling each source-release gene-identifier vocabulary into every target release under a fixed IDTrack snapshot boundary at release 114 and classifying outcomes as 1→0 (no admissible target), 1→1 (unique target), or 1→n (branching ambiguity) to obtain a release×release drift matrix whose structured asymmetries make directionality and release-local change visible rather than averaged away (Figure 4). Across this controlled window, IDTrack achieves near-complete reachability (typically > 99%) with sub-percent vocabulary loss, and surfaces the residual ∼0.1% branching ambiguity as an explicit outcome class rather than absorbing it silently—demonstrating that high-fidelity conversion and transparent ambiguity reporting are complementary rather than competing objectives, because the same infrastructure that maximizes reachability also makes the rare edge cases inspectable instead of hidden in downstream joins. To connect these discrete outcomes to biological signal, we weight conversions by pseudo-bulk UMI totals and stratify genes into expression quantiles, mapping *Cell Ranger* gene symbols from release 85 to *HGNC* symbols at target release 105 so direct target-database matches (1→1 TDM, indicating reachable HGNC symbols) are separated from alias-to-target resolutions (1→1 ATM, indicating deterministic fallback to the *Ensembl* backbone when no HGNC attachment is admissible) and from problematic outcomes; ATM outcomes concentrate in the low-expression tail (3.08% of UMIs in the 0–50% bin versus ≤ 0.11% above the 90th percentile) while UMI-level loss remains negligible (Figure 5), showing that HGNC coverage gaps disproportionately affect low-abundance genes that dominate vocabulary size but contribute marginally to biological signal. Mapping gene identifiers into a fixed target release retains essentially most UMI mass (> 99.99%) and preserves pseudo-bulk correlations (ρ_Pearson_ ≥ 0.993) even across ∼20-release gaps, yet the feature vocabulary itself exhibits measurable drift (Jaccard as low as 0.945)—which motivates treating harmonization as a publishable coordinate transform whose boundary coordinates, ambiguity semantics, and outcome profiles must be declared as Methods metadata rather than absorbed as ephemeral lookup whose provenance cannot be reconstructed (see Suppl. Sections 2.1 and 6.7 for dataset coordinates and conversion semantics).

**Fig. 4.**
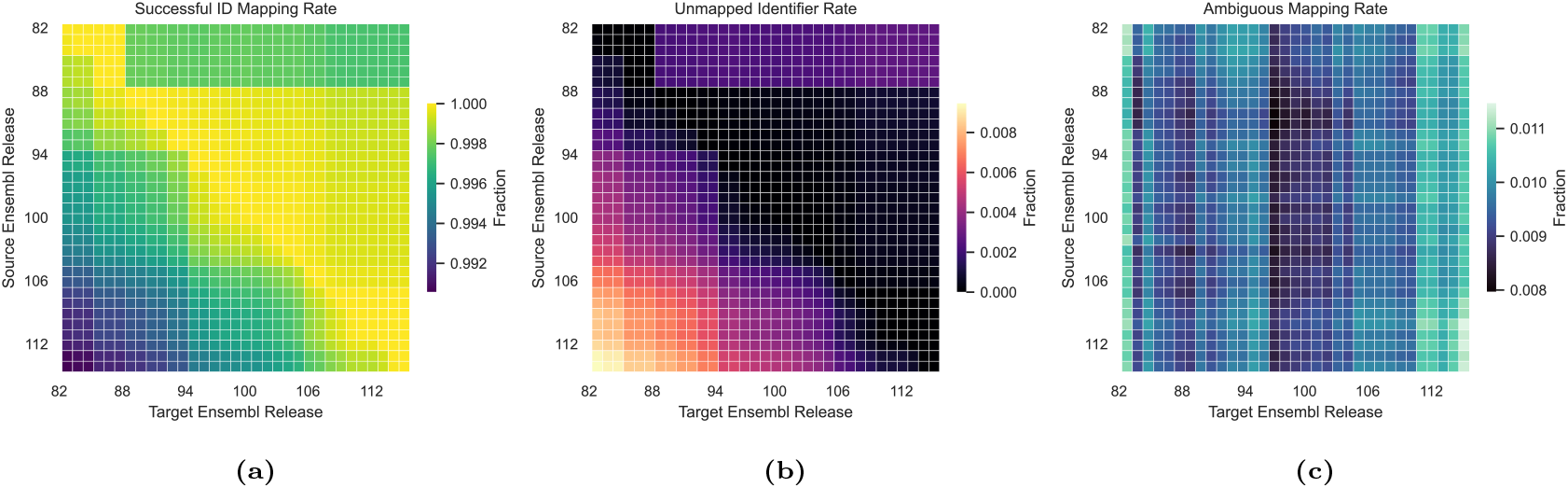
Release×release time-travel matrix quantifying identifier drift under a fixed IDTrack contract. The 10x Genomics *pbmc_1k_v3* FASTQs are quantified against *Ensembl* references spanning releases 82–114 on GRCh38, producing release-specific gene-ID vocabularies whose pairwise convertibility is then assessed by mapping each source-release vocabulary (rows) into each target release (columns) using a snapshot-bounded IDTrack graph with boundary at release 114. For each source→target pair, outputs are classified by mapping arity as 1→0 (no admissible target under the snapshot boundary), 1→1 (unique target, subdivided into direct matches and deterministic renames), and 1→n (multiple targets induced by branching history such as gene merges or splits), and the corresponding fractions over the input vocabulary are reported. (a) Reachable fraction into the target release; high values across the matrix indicate that identifier connectivity is generally preserved even across substantial release gaps. (b) Unmapped fraction (1→0); values remain sub-percent throughout, demonstrating that time-axis-aware mapping recovers nearly all identifiers within the GRCh38-era release window. (c) Ambiguous fraction (1→n); the ∼0.1% persistence of ambiguity underscores that reachability alone does not guarantee a unique feature space without declared disambiguation policy. Structured asymmetries between forward-time and reverse-time mapping directions reflect the irreversibility of merge and retirement events in *Ensembl* stable-ID lineage.

**Fig. 5.**
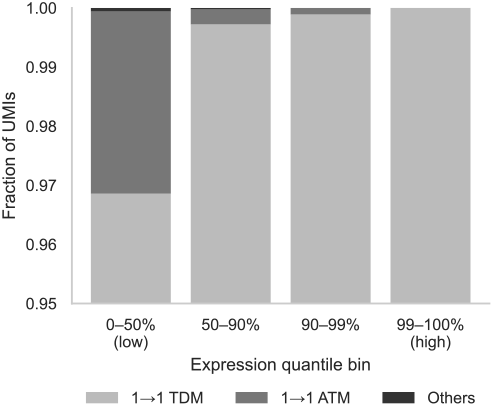
UMI-weighted distribution of mapping outcomes across expression quantiles reveals that identifier instability concentrates in the low-expression tail. Using the *pbmc_1k_v3*dataset, *Cell Ranger* gene symbols from *Ensembl* release 85 are mapped to HGNC symbols at target release 105 under a snapshot-bounded IDTrack graph, and each gene is weighted by its pseudo-bulk UMI total to report biological impact rather than vocabulary impact. Genes are stratified into four expression-quantile bins (0–50%, 50–75%, 75–90%, 90–100% of UMI totals), and for each bin the fraction of UMIs is reported according to mapping outcome: direct target-database matches (1→1 TDM, indicating reachable HGNC symbols at the target release), alias-to-target resolutions (1→1 ATM, indicating deterministic fallback to the *Ensembl* gene backbone when no HGNC attachment is admissible under the contract), or problematic (Others, combining 1→0 loss and 1→n branching ambiguity). The concentration of ATM outcomes in the low-expression tail (3.08% of UMIs in the 0–50% bin versus ≤0.11% above the 90th percentile) reflects the biological pattern that highly expressed housekeeping genes tend to have stable HGNC coverage, whereas low-abundance genes—which dominate vocabulary size but contribute marginally to quantitative signal—more frequently require *Ensembl* fallback due to incomplete HGNC attachment. Because UMI weighting emphasizes biological signal contribution rather than feature-list length, this analysis shows that harmonization preserves nearly all signal mass even when vocabulary-level metrics suggest substantial drift.

### External evidence must be bounded and ranked so completeness and ambiguity are declared policy rather than hidden defaults

The trade-off between mapping completeness and outcome ambiguity constitutes a fundamental design axis for any reproducible identifier conversion system, because external cross-references—the only mechanism capable of bridging disconnected components in a time-bounded history graph— can simultaneously restore reachability and inflate ambiguity through overlapping attachments and high-degree hubs whose transitive closure would render results unreviewable. IDTrack addresses this tension by treating external inclusion as explicit policy rather than implicit convenience: a version-controlled YAML allowlist specifies which external databases participate under which release and assembly coordinates, bounded search parameters (synonym depth limits, jump budgets, hyperconnective hub filtering) constrain how external bridges propagate through the graph, and deterministic path ranking resolves ties so that the completeness–ambiguity trade-off becomes a stable, reviewable part of the contract rather than a hidden property of endpoint state (Figure 6). Within this controlled framework, randomized stress tests quantify how outcome profiles depend on namespace semantics and target-release choice under a fixed snapshot boundary: sampling identifiers from gene, transcript, protein, and curated-nomenclature namespaces yields distinct 1→0/1→1/1→n distributions that reflect identifier granularity and cross-reference density rather than algorithmic instability, while holding queries and policy fixed and varying only the target *Ensembl* release shifts the 1→1 frequency in patterns that make drift a measurable diagnostic (Figure 7). These experiments validate that IDTrack’s output semantics are interpretable as functions of declared policy coordinates: different namespaces produce different profiles not because the algorithm is unstable but because each namespace occupies a different position in the identifier lattice, and different target releases produce different profiles not because the snapshot drifts but because the coordinate system itself evolves—which is precisely why “latest” must be replaced by an explicit, publishable boundary (see Suppl. Sections 6.3 and 6.7 for allowlist schema and search algorithm specifications).

**Fig. 6.**
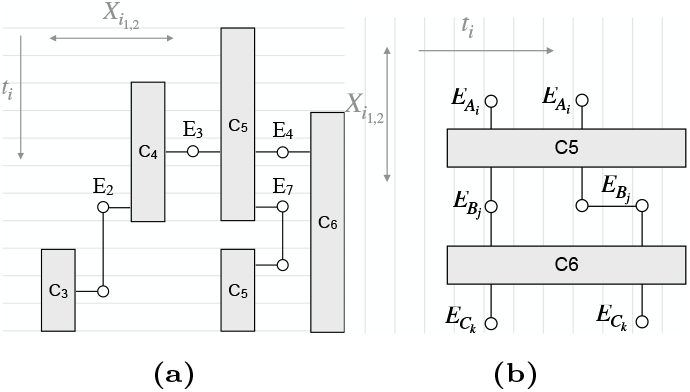
Controlled external bridging reconnects disconnected history clades while preserving explicit ambiguity semantics. *Ensembl* stable-ID history forms a directed acyclic graph where some components become disconnected under pure history traversal due to identifier retirements, assembly-specific lineages, or curation gaps that interrupt temporal continuity. (a) A single allowlisted external identifier (e.g., an HGNC symbol or RefSeq accession) can serve as an auditable re-entry point that links otherwise disconnected clades (here C3–C6), restoring temporal reachability while keeping the witness path explicit and the bridging evidence attributable to a declared external source rather than hidden in an implicit join. (b) When multiple overlapping external attachments exist, alternative bridge routes emerge that create explicit 1→n ambiguity under the declared contract; IDTrack bounds this expansion through configurable search parameters (synonym depth limits, external jump budgets, hyperconnective hub filtering with default threshold of 20) and resolves ties through deterministic path ranking, so improved reachability manifests as reportable branching rather than as an unreviewable transitive closure whose ambiguity cannot be distinguished from biological multiplicity. This design makes the completeness–ambiguity trade-off a reviewable property of the contract (allowlist contents, search bounds, ranking policy) rather than an implicit artifact of whichever cross-references happen to be returned by an endpoint at query time.

**Fig. 7.**
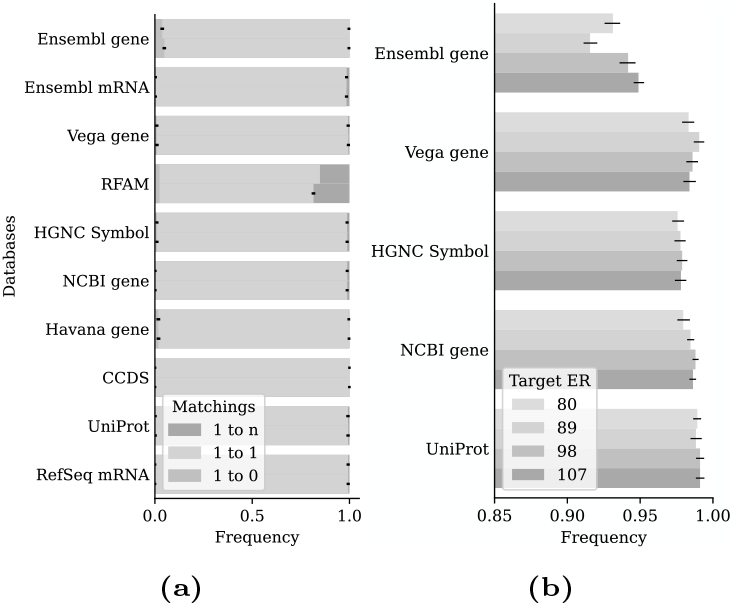
Randomized stress tests demonstrate that IDTrack’s pathfinder produces stable outcome profiles regardless of whether source-release provenance is available, and that target-release choice has minimal impact on matching success. To validate robustness under realistic conditions where identifier provenance may be incomplete, we sample 1,000 identifiers from each external database and *Ensembl* namespace (gene, transcript), convert them to *Ensembl* gene targets, and repeat the experiment six times with error bars showing standard deviation. (a) For each namespace, paired bars compare conversion outcomes when the source *Ensembl* release is known (upper bar) versus unknown (lower bar) to the algorithm; the near-identical 1→0/1→1/1→n distributions across conditions confirm that IDTrack recovers equivalent mappings even without source-release metadata, providing practical flexibility for legacy datasets whose annotation coordinates are undocumented. (b) Holding query identifiers and policy fixed while varying only the target *Ensembl* release shows that 1→1 matching frequency remains stable across release choices, demonstrating that the pathfinder’s accuracy is robust to target-release selection rather than dependent on a specific coordinate configuration. These results confirm that IDTrack’s output semantics are interpretable functions of declared policy coordinates rather than algorithmic artifacts, and that the pathfinder recovers consistent mappings even when source provenance is unavailable—precisely the condition that makes “latest” endpoints unreliable and motivates explicit, publishable snapshot boundaries.

### Atlas-scale harmonization becomes a publishable audit ledger once time is pinned as a contract

The scale and heterogeneity of multi-study atlas integration transform identifier harmonization from a preprocessing convenience into a reproducibility-critical infrastructure component, because every downstream integration, annotation, and model-training claim inherits the feature vocabulary that harmonization produces, and the mapping ledger therefore becomes the archiveable artifact upon which scientific interpretability depends rather than an ephemeral lookup whose provenance evaporates after execution. We operationalize this principle by applying IDTrack to the Human Lung Cell Atlas (19; 20; 21), harmonizing 27 datasets compiled under heterogeneous annotation coordinates into a single declared target *Ensembl* release (107) and reporting per-dataset typed outcomes (1→0/1→1/1→n) together with conversion throughput (Table 1), because multi-study integration whose constituent datasets reference different *Ensembl* releases makes naive feature joins scientifically underspecified even when downstream analysis code is version-controlled and containerized. The ledger keeps ambiguity explicit and distinguishes nomenclature updates from missingness by mapping to *HGNC* symbols when reachable under the declared contract and falling back deterministically to the *Ensembl* gene backbone when no *HGNC* attachment is admissible, so downstream vocabulary reuse remains defensible because the decision pathway for each identifier is reconstructible rather than absorbed as an implicit coercion whose provenance cannot be inspected. Across all 27 datasets, 94,538 unique input identifiers reconcile to 56,536 unique *Ensembl* gene IDs with 38,002 synonym consolidations recorded explicitly rather than silently absorbed as many-to-one merges, and throughput remains in the 𝒪 (10^2^) conversions-per-second regime (median 98 it/s for *Ensembl* targets and 133 it/s for *HGNC* targets), demonstrating that contract-driven harmonization scales to atlas-level workloads without requiring infrastructure beyond a standard computational environment. This atlas-scale application operationalizes the paper’s central claim: by pinning the time axis and evidence channels as explicit contract coordinates, IDTrack converts identifier mapping from private preprocessing into a versionable audit ledger whose decisions survive the upstream release cycle and can be inspected, compared, and archived alongside the integration workflow (see Suppl. Sections 2.2 and 6.12 for dataset retrieval coordinates and harmonizer implementation specifications).

**Table 1.** Human Lung Cell Atlas identifier harmonization demonstrates atlas-scale contract-driven mapping with explicit outcome semantics. The 27 HLCA constituent datasets (19) are harmonized under a fixed IDTrack contract targeting *Ensembl* release 107, with each row reporting input vocabulary size and typed outcome profiles for two target configurations: *HGNC* symbols (with deterministic *Ensembl* fallback) and direct *Ensembl* gene IDs. For *HGNC* targets, the 1→1 column subdivides into TDM (target-database match, indicating a reachable *HGNC* symbol at the target release) and ATM (alias-to-target match, indicating a deterministic fallback to the *Ensembl* gene backbone when no *HGNC* attachment is admissible under the same contract), so nomenclature change is distinguished from structural missingness in the archived ledger. For *Ensembl* targets, 1→1 counts unambiguous gene-level mappings and 1→n counts branching ambiguity that requires explicit disambiguation policy rather than implicit row reduction. The 1→0 column (shared across both targets) reports identifiers unreachable under the declared snapshot boundary, representing genuine upstream gaps rather than silent drops. Throughput (it/s) confirms that contract-driven conversion operates in the 𝒪 (10^2^) regime without specialized infrastructure, making atlas-scale harmonization tractable as part of standard integration workflows. Across all datasets, 94,538 unique input identifiers reconcile to 56,536 unique *Ensembl* gene IDs with 38,002 synonym consolidations recorded explicitly rather than silently absorbed as many-to-one merges, and the per-dataset profiles enable targeted inspection of studies with anomalous loss or ambiguity rates.

| Dataset | Input IDs | Target HGNC Names |  |  |  | Target Ensembl IDs |  |  | Performance (it/s) |  |
| --- | --- | --- | --- | --- | --- | --- | --- | --- | --- | --- |
|  |  | 1→1 |  | 1→n |  | 1→1 | 1→n | 1→0 | Ensembl IDs | HGNC Names |
|  |  | TDM | ATM | TDM | ATM |  |  |  |  |  |
| *Kaminski_2020* | 45947 | 33446 | 12331 | 59 | 13 | 45353 | 496 | 98 | 150.37 | 193.04 |
| *Meyer_2021* | 20922 | 20532 | 368 | 3 | 2 | 20692 | 213 | 17 | 75.08 | 94.23 |
| *MeyerNikolic_unpubl* | 33582 | 25109 | 8342 | 15 | 10 | 33186 | 290 | 106 | 114.64 | 167.62 |
| *Barbry_unpubl* | 16859 | 15155 | 1679 | 1 | 4 | 16738 | 101 | 20 | 67.29 | 83.67 |
| *Regev_2021* | 30983 | 23466 | 7387 | 8 | 9 | 30648 | 222 | 113 | 98.64 | 135.73 |
| *Thienpont_2018* | 27958 | 22394 | 5342 | 8 | 43 | 27541 | 246 | 171 | 98.29 | 123.97 |
| *Budinger_2020* | 26316 | 21441 | 4807 | 6 | 6 | 26066 | 194 | 56 | 94.27 | 136.63 |
| *Banovich_Kropinski_2020* | 33694 | 25229 | 8131 | 15 | 57 | 33096 | 336 | 262 | 115.44 | 145.69 |
| *Sheppard_2020* | 27147 | 21915 | 5020 | 6 | 40 | 26742 | 239 | 166 | 96.21 | 121.66 |
| *Wunderink_2021* | 21819 | 18856 | 2907 | 2 | 1 | 21597 | 169 | 53 | 72.65 | 101.13 |
| *Lambrechts_2021* | 33538 | 25091 | 8342 | 15 | 10 | 33168 | 290 | 80 | 115.40 | 143.91 |
| *Zhang_2021* | 18474 | 18364 | 99 | 1 | 0 | 18305 | 159 | 10 | 68.06 | 99.36 |
| *Duong_lungMAP_unpubl* | 27678 | 21617 | 5994 | 6 | 10 | 27447 | 180 | 51 | 99.63 | 125.62 |
| *Janssen_2020* | 33538 | 25091 | 8342 | 15 | 10 | 33168 | 290 | 80 | 115.16 | 143.96 |
| *Sun_2020* | 26578 | 21097 | 5417 | 6 | 8 | 26367 | 161 | 50 | 85.62 | 119.21 |
| *Gomperts_2021* | 31229 | 24929 | 5910 | 14 | 28 | 30607 | 274 | 348 | 104.83 | 153.75 |
| *Eils_2020* | 32738 | 24546 | 7519 | 17 | 44 | 31842 | 284 | 612 | 110.71 | 136.92 |
| *Schiller_2020* | 32104 | 25106 | 6431 | 8 | 26 | 31308 | 263 | 533 | 110.80 | 140.54 |
| *Misharin_Budinger_2018* | 27181 | 21945 | 5009 | 7 | 43 | 26753 | 251 | 177 | 95.28 | 120.93 |
| *Shalek_2018* | 25328 | 21154 | 3642 | 7 | 33 | 24638 | 198 | 492 | 90.33 | 133.17 |
| *Schiller_2021* | 17533 | 17330 | 193 | 0 | 0 | 17384 | 139 | 10 | 58.55 | 81.76 |
| *Peer_Massague_2020* | 19222 | 17897 | 1247 | 4 | 11 | 19004 | 155 | 63 | 70.76 | 87.07 |
| *Lafyatis_2019* | 22164 | 21693 | 442 | 6 | 2 | 21885 | 258 | 21 | 78.39 | 114.67 |
| *Tata_unpubl* | 31915 | 24230 | 7632 | 11 | 6 | 31599 | 280 | 36 | 111.79 | 140.47 |
| *Xu_2020* | 32738 | 24546 | 7519 | 17 | 44 | 31842 | 284 | 612 | 109.45 | 137.10 |
| *Sims_2019* | 60725 | 41583 | 16431 | 985 | 265 | 57434 | 1830 | 1461 | 114.41 | 157.24 |
| *Schultze_unpubl* | 24532 | 20655 | 3820 | 5 | 6 | 24290 | 196 | 46 | 89.73 | 113.41 |

## Discussion

The central insight animating IDTrack is that identifier harmonization is not a timeless dictionary lookup but a time-indexed coordinate transformation, wherein every feature label denotes a point in the joint space of namespace, *Ensembl* release, and genome assembly (often also entity layer), and the mapping layer therefore inherits a time axis that silently advances even when analysis code, container images, and downstream statistics remain unchanged. This temporal inheritance constitutes the core reproducibility failure mode in modern reuse pipelines: endpoint-style mappers and “latest” services cannot solve the time-mapping problem because, without a frozen snapshot boundary and explicit ambiguity semantics, the feature space is scientifically underdetermined and reruns become comparisons across drifting coordinate systems rather than replications of a stable methodological substrate. IDTrack materializes release history as an assembly-aware graph snapshot and pairs it with a deterministic, contract-driven pathfinder that emits explicit 1→0/1→1/1→n outcomes (with optional audit paths), converting mapping from invisible string cleanup into a publishable method object whose boundary coordinates and ambiguity semantics can be cited, compared, and archived with the same discipline expected for reference genomes and software versions. The practical consequence for authors is straightforward: publish the mapping contract (snapshot boundary and release window, assembly context, enabled external namespaces, ambiguity strategy) and, whenever possible, archive the corresponding graph snapshot, because without these coordinates the feature space cannot be reconstructed and “re-run the pipeline” is not a reproducibility claim when the mapping substrate can drift under unchanged code (8). This contract framing clarifies limitations as boundary conditions rather than deficiencies: IDTrack is not a claim about biological truth but about methodological explicitness under upstream curation, so missing mappings can reflect authoritative gaps and ambiguous multi-target outcomes can be irreducible when histories branch or cross-references overlap—conditions that make silent coercion scientifically incorrect compared to policy-controlled uncertainty propagation. Point-in-time query interfaces remain valuable for exploratory lookup when time and assembly assumptions are acceptable as implicit, but they are structurally incapable of producing an auditable, drift-aware coordinate system; IDTrack’s reusable snapshots, batch utilities, and harmonization ledgers make mapping decisions comparable objects across datasets and across years, positioning the tool as complementary infrastructure whose niche is precisely where longitudinal reproducibility matters most (extended related work appears in Suppl. Section 4). Future work therefore scopes to packaging, configuration, and coverage rather than to redefining the scientific core: expand organism and assembly presets, broaden curated external coverage, refine diagnostics and default profiles that make common pipelines reproducible out of the box—preserving the invariant that mapping decisions must remain explicit, bounded, and auditable because the time axis will continue to advance and because the identifier layer will remain the silent dependency that determines whether atlas-scale integration produces replicable science or coordinated drift.

## Author contributions statement

K.I. conceived the study and developed the methodology. K.I. and E.E. implemented the software and performed the analyses. F.J.T. supervised the project. All authors wrote and reviewed the manuscript.

## Acknowledgments

We thank the reviewers for their valuable feedback.

## Supplementary Materials for IDTrack

## 1. Supplementary Overview

This supplementary document accompanies the main manuscript (Sections 1–4) and is intentionally text-only, because its role is to make the mapping contract reconstructible from concrete artifacts, code anchors, and explicit decision rules rather than to add new experimental claims. It expands related work as a capability taxonomy (rather than a benchmark), states assumptions and limitations as disciplined scope boundaries, and provides implementation-level method specifications for the two core objects that define IDTrack’s behavior: a snapshot-bounded, assembly-aware identifier graph and a contract-driven pathfinder that treats *Ensembl* releases as a time axis. Because identifier drift is structured history rather than random noise, this supplement is written as an audit layer for time-axis reasoning, defining the graph and pathfinder as the method’s scientific objects and treating every configuration knob as a coordinate-system choice that deterministically changes reachability, ambiguity, and output semantics.

Operationally, the main paper is the compressed argument and evidence, while this supplement is the contract specification: when the main text says “snapshot boundary”, “time axis”, “controlled bridging”, or “explicit ambiguity semantics”, the sections below name the corresponding parameters, default values, and decision rules that determine reachability, ambiguity, and output structure. The intended reader outcome is that methodological explicitness can be evaluated without supplementary figures and without rerunning analyses, because each claim is tethered to an artifact identity and to the public API surface that materializes it.

Because IDTrack’s scientific contribution is to turn identifier mapping into a publishable practice rather than a private preprocessing script, this supplement keeps the application surface explicit: the same contract object underlies dataset harmonization before integration, legacy gene-list rescue, reproducible reuse of shared matrices, cross-database interoperability when collaborators mix identifier conventions, and traceability in variant/regulatory interpretation, proteomics aggregation, perturbation pipelines, and clinical reporting. We treat the external allowlist as a method knob rather than as a headline feature, because its scientific role is to bound which cross-reference channels are allowed to influence reachability and ambiguity under a fixed snapshot, and because the main differentiator remains time-axis tracking implemented by the graph and pathfinder.

For practical reuse, the intended reading mode is checklist-like: a reader should be able to extract a complete contract from Methods and verify it against concrete artifact identifiers (graph snapshot filename plus configuration identity and hashes), which is the minimum required for FAIR reuse when upstream resources drift (1). Even without adopting IDTrack, the same reporting discipline applies ecosystem-wide, because under drift any conversion workflow that fails to publish its boundaries is scientifically underspecified by construction. For navigation, Suppl. Sections 3–5 provide contract framing, capability context, and disciplined scope boundaries for treating mapping as a publishable coordinate system under drift. Suppl. Sections 6–7 then provide code-anchored method specifications and a provenance ledger plus copy/paste reporting template that make mapping claims reconstructible.

## 2. Dataset availability

The manuscript-facing experiments use two public datasets whose retrieval coordinates are part of the reproducibility contract.

### 2.1. 10x Genomics PBMC 1k (v3 chemistry)

We use the 10x Genomics “1k PBMCs from a Healthy Donor” dataset (v3 chemistry, 3’ gene expression; v3.0.0), downloaded as `pbmc_1k_v3_fastqs.tar` from https://cf.10xgenomics.com/samples/cell-exp/3.0.0/pbmc_1k_v3/pbmc_1k_v3_fastqs.tar (2). The dataset landing page is https://www.10xgenomics.com/datasets/1-k-pbm-cs-from-a-healthy-donor-v-3-chemistry-3-standard-3-0-0 (2). We verify integrity via the published MD5 checksum `n265ebe8f77ad90db350984d9c7a59e52` (2). The vendor-reported design targets approximately 1,000 cells (expected 1,222) with a mean of 54,502 reads per cell, and our *Cell Ranger* runs (3) use an expected cell count of 1,000 while disabling BAM generation so that downstream differences are attributable to reference-release choice rather than to extraneous output artifacts. The main manuscript’s PBMC figures therefore treat this fixed FASTQ archive as the shared raw input whose feature-space drift is induced solely by reference and mapping choices.

### 2.2. Human Lung Cell Atlas (HLCA)

We use the Human Lung Cell Atlas (HLCA) as described by Sikkema et al. (4), and we follow the authors’ public distribution for both the integrated atlas and the reference model used for mapping new data. The integrated HLCA collection (raw and normalized counts, integrated embedding, annotations, and metadata) is available via cellxgene at https://cellxgene.cziscience.com/collections/6f6d381a-7701-4781-935c-db10d30de293 (5), and the collection snapshot used in this work contains 2,282,447 cells. For reference-based mapping experiments we use the HLCA reference model and full embedding released on Zenodo (version 1.1) at https://doi.org/10.5281/zenodo.7599104 (6), which provides the reference model and model extensions (scArches surgery adapters) needed to map new datasets into the atlas coordinate system. For reproducibility notebooks that operate on per-study `.h5ad files`, we assume the HLCA reproducibility layout (environment variable `HLCA_BASE_PATH`) and treat the exact study-to-file mapping as part of the experiment code rather than as implicit filesystem knowledge.

## 3. From Lookup Tables to Mapping Contracts

### 3.1. Gaps (mapping as a declared coordinate system rather than a lookup table)

Under identifier drift, conversion is not a timeless dictionary lookup but a coordinate transformation in the joint space of namespace × *Ensembl* release × genome assembly × entity layer (gene/transcript/protein), so every feature-level claim inherits hidden degrees of freedom unless the boundary is published as Methods metadata rather than assumed by a default endpoint. The recurring ecosystem gaps are therefore not about effort or curation quality but about missing contract surfaces: release semantics are often implicit (“latest” servers or unlabeled tables), assembly semantics are under-modeled (build-scoped resources treated as interchangeable), entity-layer normalization is ad hoc (genes mixed with versioned transcripts/proteins), external cross-reference participation is rarely declared as a policy object, and one-to-many ambiguity is commonly coerced by downstream joins instead of surfaced as an explicit outcome class. Because *Ensembl* stable identifiers can be retired, merged, split, and versioned while symbols and external cross-references evolve asynchronously, a single input string rarely denotes a unique entity without a declared coordinate system and a declared disambiguation policy, and the absence of that declaration turns mapping into an unreported experimental condition that drifts between reruns under unchanged analysis code. The practical consequence is that reproducibility is usually attempted at the wrong level: pipelines version-control analysis code while allowing the mapping layer to float with endpoint state, which is precisely where cross-study integration, long-horizon reuse, and atlas building are most sensitive to silent feature non-equivalence. IDTrack’s scientific motivation is therefore not to replace point-in-time mappers, but to make boundary choices publishable by snapshot-bounding the *Ensembl* history substrate, treating assembly as a first-class coordinate, controlling which external namespaces contribute evidence under a versionable allowlist, and returning auditable 1→0/1→1/1→n outcomes (with optional paths) that can be archived alongside the analysis.

### 3.2. Application surface (where drift becomes a scientific confound rather than a nuisance)

In atlas construction and single-cell integration, cohorts processed across years routinely mix *Ensembl* releases, GRCh37/GRCh38-era annotations, and symbol conventions, so integration quality, marker interpretability, and cross-study differential expression can be altered by silent feature non-equivalence rather than by biology, particularly when raw-read reprocessing into a single reference is infeasible (7; 8; 9). In reference mapping and label transfer, the learned correspondence between query and atlas cells depends on the exact feature vocabulary used during training, so drift in upstream identifier normalization can shift embedding geometry, perturb classifier weights, and complicate the interpretation of “model updates” when the effective transformation was a moving mapping backend (10). In spatial transcriptomics, multiome, and regulatory pipelines, feature identity couples gene models to assembly-scoped coordinates through peak-to-gene links, gene activity matrices, transcript selection, and variant-to-gene mapping, so conversions that ignore release and build context can change regulatory assignments and cross-modality alignment even when the statistical method is unchanged (11; 12; 13). In long-horizon reanalysis, meta-analysis, and the reuse of published count matrices where FASTQs and aligner parameters are unavailable, the mapping layer determines which historical identifiers are treated as equivalent, which are declared unmappable, and which are surfaced as branching ambiguity, thereby controlling what can be pooled without pretending the past conforms to the present. In legacy gene-list rescue for replication and functional interpretation (older microarray signatures, pathway gene sets, curated marker panels), symbol reuse and withdrawn names can create false agreement across studies unless symbol history and snapshot boundaries are recorded, making identifier normalization a prerequisite for scientifically meaningful comparisons rather than a clerical step. In perturbation and screening workflows (CRISPR libraries, Perturb-seq, pooled functional genomics), target definitions are often authored in symbols or legacy IDs at library design time, so drift-aware mapping is required to preserve what “the same target” meant historically while still enabling modern annotation and cross-study aggregation (14; 15). In proteomics and proteogenomics, protein accessions, isoforms, and gene-level identifiers form intrinsically many-to-many relationships across *UniProt/RefSeq/Ensembl* layers, so pipelines that collapse mappings implicitly can propagate inconsistent aggregation choices into quantitative comparisons, differential protein inference, and cross-omics integration (16; 17; 18). In clinical and translational reporting, assay pipelines and gene panels must remain traceable across updates to reference annotations, so unreported mapping boundaries complicate audit trails during re-interpretation, multi-site harmonization, and regulatory reproducibility when laboratories adopt different identifier authorities or genome builds (19). In variant annotation, QTL interpretation, and regulatory genomics, mappings between variants, transcripts, and genes are conditioned on assembly and transcript models, so converting identifiers across databases without explicit build and release semantics can alter variant-to-gene links, causal-gene prioritization, and downstream pathway interpretation even when summary statistics are unchanged (20). In knowledge graphs, pathway/network resources, and machine-learning feature engineering, identifier normalization is the join operation that fuses heterogeneous databases into a single graph or matrix, yet drift can change gene-set membership, duplicate or split nodes, and rewire edges, meaning that “the same model” is not scientifically comparable unless the mapping transformation is pinned and reported (21; 22). In data portal ingestion and large collaborative consortia, identifiers are the boundary object exchanged across teams, so a publishable mapping contract supports staged updates under explicit snapshot changes, prevents silent divergence between pipelines, and enables reviewers to reconstruct the exact feature space that underlies a reported result. Across these settings, the scientific gap is not that mapping is hard, but that mapping is treated as an internal convenience whose assumptions are rarely reported at the granularity needed to make cross-time reuse defensible, especially when dataset reuse outlives the release cycle of upstream resources.

### 3.3. Failure modes that remain invisible without a mapping contract

The most common failures are systematic rather than dramatic: stripping transcript/protein version suffixes, defaulting to “current” endpoints, mixing GRCh37- and GRCh38-scoped resources, and treating symbols as stable keys, which together can create consistent but hidden feature divergence across cohorts and across reruns of the same pipeline. Even when a mapper returns a table, ambiguity is typically resolved implicitly by row-reduction logic outside the mapper (first hit wins, duplicates dropped, many-to-many exploded and then filtered), so the effective mapping policy lives in downstream joins and aggregation code rather than in a reportable object whose semantics can be audited. Because external namespaces differ in granularity and curation (gene IDs vs transcript/protein accessions; curated nomenclature vs computed cross-references), broad external inclusion can improve connectivity while amplifying ambiguity and hyperconnectivity, making results depend on invisible configuration choices and on transitive paths that are not scientifically intended. A drift-aware mapping layer therefore needs explicit missingness semantics (distinguishing “no corresponding node” from “no path under the declared window”), explicit ambiguity semantics (one-to-zero (1→ 0), one-to-one (1→1), one-to-many (1→n) under a declared strategy), and optional explainability that renders mappings reviewable as structured evidence rather than as ad hoc narrative. IDTrack’s contract framing operationalizes these requirements as parameters (snapshot boundary and window, assembly context, external inclusion policy, ambiguity strategy, explainability toggle) and as archiveable artifacts (cached graph snapshot plus configuration identity), so conversion becomes an auditable, rerunnable transformation rather than an undocumented pre-processing step. The point of this section is not to argue that other tools should behave like IDTrack, but to make the scope boundary explicit: tools optimized for convenience lookup are excellent at answering “what does this mean today” while being structurally ill-suited for publishing “what coordinate system produced the feature space in this analysis”.

### 3.4. Axes and rules (capability-first, not accuracy-first)

This extended comparison is written as a capability taxonomy because benchmarking “accuracy” across endpoints is ill-defined under drift, and because tool outputs depend on unreported choices (release hosts, xref inclusion, normalization, and row-reduction) that preclude a timeless ground truth for the conversion problem. We compare tools by whether they expose time-awareness (explicit release semantics rather than implicit “latest”), assembly-awareness (explicit build scoping), entity-layer specificity (gene vs transcript vs protein with version handling), and ambiguity semantics (one-to-zero/one-to-one/one-to-many, i.e., 1→0/1→1/1→n under a declared policy rather than post-hoc filtering). We additionally consider auditability (inspectable provenance or paths), artifact discipline (cacheable substrates that can be archived), configuration explicitness (versionable inclusion/exclusion of external namespaces), and pipeline integration (batch conversion plus diagnostics rather than interactive lookup), because these are the properties required when mapping is a published methodological transformation. Throughout, we treat “supports archives” as necessary but insufficient, because pinning an archive host answers “which release did I query” but does not answer “what mapping policy did I apply” when one-to-many relationships and transitive cross-references are present. We likewise treat “can run locally” as necessary but insufficient, because a local cache without a declared snapshot identity and ambiguity policy is still a moving mapping substrate when rebuilt from updated sources.

### 3.5. Fairness rule

Each tool below is described by what it was designed to do well— querying authoritative live services, providing broad aggregation, enabling enrichment workflows, distributing versioned annotation packages, or abstracting mapping backends—because that design center determines appropriate expectations about drift, offline reproducibility, and output semantics. We then state the boundary it does not attempt (a scope choice rather than a defect) and use that boundary to motivate IDTrack’s niche as a publishable mapping contract for drift-aware harmonization, so the comparison stays respectful while still making the scientific gaps explicit. This framing also makes complementarity concrete, because the question is not which tool “wins”, but which tool produces an auditable coordinate system that can be carried across datasets and across years.

### 3.6. Reporting rule (contract discipline even without IDTrack)

Because drift makes conversion a coordinate transformation rather than a timeless lookup, any workflow that uses an external mapper should report the endpoint boundary (service/host, query time, and data version or release identifier), the assumed assembly/build context, the identifier layer and normalization applied (e.g., version-suffix handling), and the ambiguity policy used to reduce one-to-many outputs. When these elements are left implicit—current endpoints, hidden allowlists, default schema joins, and silent duplicate suppression—the mapping layer remains a hidden dependency even when the downstream analysis code is fully version-controlled, and reviewers cannot reconstruct which feature space the results actually used. IDTrack’s position is that these parameters should be elevated into first-class metadata and, whenever possible, into archiveable artifacts, because only then can mapping be rerun without requerying moving endpoints and only then can ambiguity be interpreted as an output rather than as an implementation artifact. Even when IDTrack is not used, adopting this contract discipline makes the mapping layer reviewable, which is the minimal standard required when identifier choices condition claims about differential expression, cell-type labeling, network connectivity, or variant-to-gene interpretation.

## 4. Extended Related Work

This section expands ecosystem comparison as a capability taxonomy rather than a benchmark, because under drift endpoint outputs are conditioned on unreported choices and a timeless ground truth for conversion is ill-defined. For each tool family we state its design center and the boundary it does not attempt (a scope choice rather than a defect), and we use that boundary to motivate IDTrack’s niche as a publishable mapping contract under *Ensembl* release drift and assembly heterogeneity. The objective is not to rank tools but to make complementarity explicit: point-in-time query interfaces and aggregation services remain essential for exploration and annotation, while contract-driven graph substrates are required when mapping decisions must survive as Methods metadata across years and datasets.

### 4.1. BioMart/biomaRt/pybiomart as *Ensembl*-centric query interfaces

*BioMart* is a mature query framework over *Ensembl*-style relational schemas, exposing filters and attributes for genes, transcripts, proteins, and cross-references, and it remains an efficient choice when the scientific requirement is point-in-time retrieval from a chosen host (including *Ensembl* archive sites) and when the deliverable is a flat table that can be joined into an analysis pipeline (23; 24). In practice, *biomaRt (R)* and *pybiomart (Python)* workflows materialize local annotation snapshots—symbols, synonyms, xrefs, gene biotypes, coordinates, and various per-gene attributes—and they are widely used to translate between identifier columns, enrich gene lists with metadata, and export mapping tables that serve as downstream caches. The strength is schema access without local database staging: users specify a dataset, declare a query as filters plus attributes, and retrieve a representation of a particular release if the archive host is pinned and the query specification is preserved, which is operationally reproducible at the level of rerunning the same query against the same host.

However, even with archive pinning, the dominant interaction is still query-to-table rather than contract-to-artifact, because *BioMart* does not force users to declare mapping semantics as a portable object that records assembly scoping, transcript/protein version handling, cross-reference source inclusion, and an interpretation of one-to-many relationships beyond row duplication. Ambiguity is therefore usually resolved implicitly by table-merging behavior—duplicate rows are exploded or collapsed according to how joins and groupings are written—and many pipelines silently coerce a one-to-one view by taking first matches, dropping duplicated keys, or aggregating without recording which rule was applied, making the effective mapping policy an implementation detail. Cross-release harmonization is possible but typically requires analysts to coordinate multiple archive queries and reconcile schema and attribute differences across releases, which is precisely where merges, splits, and conflicting cross-references become scientifically consequential and where ad hoc row-reduction rules are most likely to diverge between teams. IDTrack is positioned orthogonally: it builds a snapshot-bounded, assembly-aware history graph once, it makes external cross-reference participation an explicit inclusion policy rather than an implicit set of selected xref columns, and it returns standardized one-to-zero/one-to-one/one-to-many outcomes (with optional audit paths) so ambiguity is a reported semantic rather than a hidden join artifact. In practice, the approaches compose well: *BioMart* remains a convenient way to fetch rich annotation attributes once IDTrack has defined a release-pinned feature space, while IDTrack provides the contract object that makes the mapping layer itself reproducible, auditable, and reviewable.

### 4.2. MyGene.info as a broad identifier aggregation service

*MyGene.info* is a gene-centric aggregation service built on the *BioThings* stack, providing a high-level API that resolves heterogeneous user inputs (symbols, aliases, accessions, *Entrez* IDs, *Ensembl* IDs) into normalized gene records with cross-references and annotations across many upstream sources (25; 26). Its practical value is speed and breadth: batch endpoints accept mixed-namespace queries at scale, return structured hits with per-record scores and per-source fields, and often surface multiple candidate records when the input is under-specified, which moves ambiguity from silent failure into an explicit set of candidates that users can inspect. Because *MyGene.info* exposes metadata about the aggregated build and source versions, a pipeline can at least record the service-defined snapshot that generated a mapping table, which is a meaningful reproducibility improvement relative to unlabeled ad hoc scrapes from web interfaces.

However, the time axis in *MyGene.info* is the evolving aggregate build rather than an explicit *Ensembl* release coordinate system, and cross-references reflect asynchronous updates across sources, so the same query can yield different xref sets as upstream resources advance even when client code is unchanged. This is not a flaw—*MyGene.info* is optimized for providing current, broad coverage—but it means that turning its outputs into a publication-grade mapping contract requires additional discipline: build metadata must be captured and downstream row-selection rules must be stated when multiple hits, multiple xrefs, or conflicting evidence appear. Furthermore, because the service is gene-centric, conversions that depend on transcript/protein granularity, accession-version semantics, or *Ensembl* release-history time travel (including merge/split events) are not naturally represented as auditable paths through a time-bounded substrate, and pipelines can unknowingly mix gene-level and transcript-level reasoning when inputs include versioned accessions. IDTrack therefore does not compete on being a general aggregation service; it competes on making boundary choices explicit (snapshot boundary and window, assembly context, external inclusion policy) and on returning explicit one-to-zero/one-to-one/one-to-many outcomes with optional explainability paths, so longitudinal and atlas-scale integrations can publish the mapping transformation that defined their feature space. In an IDTrack-centered workflow, *MyGene.info* remains valuable for exploratory lookup and namespace discovery, but its outputs are most responsibly treated as upstream suggestions whose build identity and ambiguity handling must be recorded, because otherwise they reintroduce hidden drift into the mapping layer. Operationally, IDTrack exposes *MyGene.info* as an optional convenience backend for cross-namespace lookups, but it treats live aggregation as helper evidence rather than as the published substrate, keeping the contract anchored in snapshot-bounded *Ensembl* history graphs.

### 4.3. Gene nomenclature authorities as symbol-history references

Nomenclature authorities form a distinct class of identifier resources, where the primary object is not a mapping service but a curated policy for gene naming, symbol approval, and symbol-history tracking, exemplified by the *HGNC* resources for human genes (27; 28). These resources are crucial because gene symbols are pervasive in experimental reporting, legacy matrices, and wet-lab protocols, yet they are not stable identifiers: symbols can be updated, withdrawn, reassigned, or reused, and alias lists encode many-to-one and one-to-many historical relationships that are invisible when symbols are treated as immutable keys. *HGNC* in particular provides stable *HGNC* identifiers, curated approved symbols, previous symbols, alias symbols, withdrawal semantics, and cross-references to resources such as *Ensembl, NCBI* Gene, and UniProt, making it a principled starting point for interpreting symbol-bearing inputs when the goal is to map them into stable identifier coordinates. Across organisms, analogous committees and knowledge bases (for example MGI, FlyBase, ZFIN, RGD) provide approved symbols, synonym sets, and withdrawal policies, but these policy boundaries differ, so “symbol normalization” is only well-defined when the authority and its snapshot assumptions are declared. As a result, symbol-to-identifier conversion is scientifically safe only when it is treated as time- and policy-dependent— which authority and which snapshot were assumed, whether withdrawn symbols were allowed, and how alias collisions were handled—because otherwise apparent cross-study agreement can reflect symbol reuse rather than biological consistency. Nomenclature databases typically provide stable internal IDs (e.g., *HGNC* IDs) and explicit history fields that support defensible symbol interpretation, yet they do not attempt to provide *Ensembl* release-history time travel, assembly-aware reconciliation, or a unified substrate that spans transcript and protein layers across external namespaces. IDTrack treats symbol-based namespaces as externals whose participation is explicitly allowlisted and interpreted under an *Ensembl* snapshot boundary, which allows symbol inputs to be time-traveled into a declared *Ensembl* coordinate system while still requiring the user to report the symbol authority boundary when that matters for interpretability. In practice, nomenclature authorities and IDTrack are complementary: nomenclature databases define what a symbol should mean under a curated naming policy, while IDTrack defines how symbol-bearing datasets are reconciled with release- and assembly-explicit *Ensembl* histories so that symbol normalization does not silently change the feature space of an integration workflow.

### 4.4. g:Profiler as analysis-first tooling with conversion as a convenience

*g:Profiler* is designed around functional profiling and pathway analysis (*g:GOSt*), with identifier conversion (*g:Convert*) serving as an enabling step that makes enrichment workflows accept heterogeneous user inputs without requiring users to manually assemble mapping tables (29). In typical use, the conversion layer is valuable precisely because it is frictionless: it tolerates mixed identifier types, performs normalization over a broad set of supported namespaces, and returns outputs that can be fed directly into enrichment and reporting workflows with minimal user configuration. Because the tool is oriented toward analysis rather than substrate publication, the conversion output is naturally shaped as an intermediate table keyed to the service’s internal data versioning, which is appropriate when the scientific objective is to obtain an interpretable enrichment result under a known backend snapshot. The *g:Profiler* ecosystem also emphasizes reproducibility within its own release system by documenting data versions and offering archive access, which is useful when the analytic goal is to reproduce the same enrichment analysis under the same backend state rather than to define a reusable mapping contract that spans years of heterogeneous dataset provenance.

Under drift-aware dataset harmonization, however, the mapping problem is more constrained and more demanding: the coordinate system must be an explicit *Ensembl* release and assembly context, the mapping must be a reusable artifact that can be rerun offline, and ambiguity needs to be surfaced as an output semantic rather than handled implicitly as a list-cleaning step. Because *g:Profiler ‘s* primary deliverable is enrichment rather than a mapping substrate, conversions are commonly consumed without an explicit contract object that records assembly semantics, external inclusion policy, and standardized one-to-zero/one-to-one/one-to-many outcomes, so drift and ambiguity can re-enter the pipeline through unreported defaults. IDTrack targets this different objective by treating conversion itself as a publishable contract with snapshot-bounded *Ensembl* release history, explicit assembly handling, and versionable external allowlists, and by returning auditable outcomes (and optional paths) that can be attached to integration diagnostics. In practice, the tools are complementary: an analysis can use IDTrack to produce a release-bounded identifier set (with explicit ambiguity handling) and then apply *g:Profiler* for enrichment within that explicitly defined feature space, ensuring that interpretive biology is not confounded by a drifting or implicit mapping layer.

### 4.5. gget as a pragmatic API-wrapper ecosystem tool

*gget* provides a command-line and *Python* interface that wraps multiple genomic reference resources behind a uniform scripting surface, lowering friction when the task is to fetch current metadata, sequences, or cross-references in notebooks and lightweight pipelines (30). Its design center is integration-by-ergonomics: established endpoints such as *Ensembl, NCBI, UniProt*, and *Enrichr* are exposed as compact subcommands and Python calls, making common retrieval tasks easy to share and easy to embed in exploratory analysis code without having to learn each upstream API separately. For identifier-related tasks, this wrapper posture is often sufficient when the scientific question is exploratory or when the mapping step is not intended to survive as a Methods-level artifact beyond the immediate analysis session, because users typically want an up-to-date interpretation of an identifier rather than a frozen coordinate system.

The drift-aware boundary is that wrapper convenience is not a reproducibility contract: if the wrapped endpoint changes its default release, schema, or cross-reference logic, the mapping changes, and unless users pin hosts, capture response metadata, and archive outputs, the transformation cannot be reconstructed reliably at manuscript review time even when the wrapper code is public. Moreover, because *gget* spans heterogeneous resources with different assumptions, assembly and identifier-layer semantics can vary across calls (gene vs transcript vs protein, versioned accessions vs unversioned IDs, build-scoped endpoints vs general endpoints), and pipelines that do not make these distinctions explicit can silently mix incompatible coordinates. IDTrack therefore treats endpoint access as a graph-building or optional-helper phase whose products are pinned under a snapshot boundary, and it surfaces ambiguity and failure modes explicitly so downstream integration can treat mapping as declared policy rather than absorb it as an invisible side effect. In practice, *gget*-style wrappers remain excellent for answering “what does this identifier mean today”, while IDTrack is designed for the complementary question “what mapping contract produced the feature space in this paper, and can we rerun it exactly from archived artifacts”, which is the drift-aware niche this manuscript argues should be treated as a Methods-level responsibility.

### 4.6. *Ensembl* REST as an upstream metadata endpoint

The *Ensembl* REST API is a canonical upstream interface for programmatic access to *Ensembl* reference data, exposing endpoints for stable-ID lookups, cross-references, sequences, genomic overlaps, and stable-ID history queries, and it is often the most direct way to retrieve authoritative metadata in workflows that are already *Ensembl*-centric (31; 32). A key strength is explicitness at the request level: a client can specify a species, identifier, coordinate, and content type, retrieve structured JSON that reflects *Ensembl*’s current schema, and integrate these queries into pipelines without staging the full *Ensembl* MySQL databases locally. *Ensembl* also maintains special endpoints for assembly contexts such as the GRCh37 REST server and archive sites for past releases, which makes it possible to query a chosen coordinate system when the scientific requirement is to interpret identifiers and annotations in the context in which a dataset was originally produced. However, REST remains an endpoint for retrieval rather than a substrate for publication, because a sequence of API calls does not, by itself, define a reusable mapping contract that spans multiple releases, records the admissible time window, and standardizes how one-to-many histories and cross-reference conflicts should be represented in outputs. In practice, analysts often use REST to answer per-identifier questions or to populate lookup tables on the fly, and these tables then become implicit mapping layers whose provenance is incomplete unless the exact host, release, assembly context, and response metadata are captured and archived. The drift-aware consequence is that even small default changes—which release a host points to, how xref sets are curated, how synonyms are normalized, or which transcripts are considered canonical—can silently alter downstream feature identity while leaving pipeline code unchanged, making cross-study comparability brittle. IDTrack instead uses *Ensembl*’s release-history resources to build a snapshot-bounded graph artifact that encodes stable-ID lineage as a time axis, enforces assembly-aware reachability rules, and produces auditable conversion outcomes with optional paths, so the mapping layer can be rerun offline and inspected as a method object rather than recreated from live queries. Seen this way, *Ensembl* REST is indispensable upstream metadata infrastructure, while IDTrack is a downstream reproducibility layer that materializes the particular subset of *Ensembl* history and cross-reference evidence that a paper chooses to treat as its coordinate system.

### 4.7. Genome coordinate liftover tools as assembly-bridge infrastructure

Genome coordinate conversion tools such as the *UCSC liftOver* utility and *CrossMap* implement assembly-to-assembly remapping for genomic intervals and variants by applying chain files derived from whole-genome alignments, providing a practical bridge when the scientific object is a locus in one build that must be located in another build without rerunning upstream alignment (33; 34). These tools are foundational in variant annotation, peak calling reuse, and interval-based reproducibility, because they translate the spatial coordinate system of the genome while preserving the semantics of sequence-based mapping under a specified chain file. Their scope boundary is that coordinate liftover is not identifier time travel: a lifted interval can overlap different gene models across releases, and gene, transcript, and protein identifiers may change even when the underlying genomic sequence is conserved, so a coordinate bridge does not by itself define equivalence of biological entity identifiers. Moreover, liftover outputs rarely carry a downstream-friendly ambiguity semantic beyond success/failure or multi-mapping loci, so the interpretation of interval-to-gene assignments still depends on which annotation release was used after liftover and how ambiguous overlaps were handled. IDTrack’s assembly awareness addresses a different layer of the problem by treating assembly as part of identifier meaning and by constraining reachability across *Ensembl* releases and external namespaces under a snapshot boundary, while explicitly not replacing alignment-based liftover when the scientific question is about genomic intervals rather than entity identifiers. In drift-aware workflows, the two are therefore complementary: liftover stabilizes spatial coordinates across builds, while IDTrack stabilizes identifier coordinates across release history and namespace boundaries so that feature-level matrices remain comparable when gene models and naming conventions change.

### 4.8. *UniProt* mapping services as protein-centric infrastructure

*UniProt* is the central protein knowledge infrastructure, maintaining curated protein records, accession and isoform semantics, rich functional annotation, and extensive cross-references, and its identifier mapping services provide practical conversion between *UniProt* accessions and many external identifier systems via web and API interfaces (16). This protein-centric posture is scientifically appropriate when the objective is to interpret proteins, isoforms, and functional annotations rather than to define a gene-level feature space, and *UniProt*’s curation and cross-reference breadth make it a common pivot point in proteomics and proteogenomics workflows. *UniProt* mapping is also operationally robust: batch jobs can map large identifier lists and return structured results, which supports reproducible reporting when the *UniProt* release version and input accession versions are recorded. However, *UniProt*’s time semantics are organized around *UniProt* releases and protein record histories rather than around *Ensembl* release history, and cross-references to *Ensembl*, RefSeq, and gene symbols update asynchronously, so *UniProt* mapping does not provide a native time axis that can project arbitrary historical *Ensembl* identifiers into a declared *Ensembl* snapshot. Furthermore, protein-to-gene relationships are intrinsically many-to-many across isoforms, splice variants, and gene families, so conversions that pass through protein identifiers can introduce implicit aggregation and evidence mixing that is correct in protein biology but can be an unintended confound when the goal is a stable gene-level vocabulary for cross-study expression integration. IDTrack does not replace *UniProt*; it treats *UniProt*-derived namespaces as optional externals within a snapshot-bounded *Ensembl* history graph, so *UniProt* participation is controlled by an explicit inclusion policy and interpreted under the same release and assembly boundary that defines the target feature space. Practically, *UniProt* mapping can be seen as authoritative protein annotation infrastructure, while IDTrack provides the release- and assembly-explicit contract needed when heterogeneous transcriptomic datasets must first be reconciled into a single *Ensembl* coordinate system before protein-centric interpretation is applied. This separation of responsibilities is operationally useful: IDTrack can time-travel heterogeneous identifiers into a declared *Ensembl* snapshot and expose ambiguity explicitly, after which *UniProt*-centric analyses can proceed within a stable feature space without making the mapping layer depend on a live mapping backend at review time.

### 4.9. NCBI Gene/*RefSeq* resources as curated sequence databases

*NCBI* Gene and *RefSeq* provide curated identifiers and reference sequences across genomes, transcripts, and proteins, with explicit accession-version semantics for *RefSeq* records and gene-centered organization that supports sequence-based workflows, clinical reporting, and broad cross-referencing to external resources (35; 36; 17). These resources are often the authoritative substrate when the object of analysis is a reference sequence (transcript or protein accessions with versions) rather than a particular *Ensembl* release snapshot, and their explicit versioning is a strong reproducibility feature when accession versions and genome builds are carried through to downstream pipelines. *NCBI* also exposes history and replacement metadata for genes and accessions, which supports principled handling of discontinued records and helps distinguish retirements from merges in workflows that need to audit identifier evolution.

However, cross-resource conversion between *NCBI* and *Ensembl* is not a trivial relabeling: the coordinate systems differ, annotation update cycles are asynchronous, assembly assumptions can diverge across time, and transcript/protein granularity introduces many-to-many mappings that reflect alternative splicing, record updates, and differing gene-model boundaries. As a consequence, a pipeline that maps *Ensembl* IDs to *RefSeq* (or vice versa) without explicitly stating which releases, assemblies, and normalization rules were assumed can silently change feature identity and even apparent gene membership when run later or in a different environment, because “the same” gene symbol or accession may point to a different model after curation updates. In practice, the most reproducible way to consume *NCBI* -to-*Ensembl* links is to treat the chosen snapshot (*NCBI* /*RefSeq* release, genome build, accession-version policy, and mapping table version) as part of Methods metadata and to archive the exact mapping files used, yet many pipelines consume cross-references through live services, cached but unlabeled files, or implicit defaults. IDTrack’s contribution is to make those boundary choices part of the conversion contract and to integrate *NCBI* -derived namespaces as explicit externals in a snapshot-bounded *Ensembl* history graph, so that conversions are rerunnable from local artifacts and ambiguous cases are surfaced as explicit outcomes rather than as dropped duplicates. In this framing, *NCBI* remains curated sequence infrastructure, while IDTrack provides a release- and assembly-explicit method for reconciling *NCBI* - linked identifiers with *Ensembl* time travel when the published unit is a shared feature space across heterogeneous studies.

### 4.10. Bioconductor-style static annotation packages

*Bioconductor ‘s* annotation ecosystem—including AnnotationDbi and organism packages such as `org.*`, build-specific resources such as `TxDb` and `EnsDb`, and a broader distribution model through *Bioconductor* releases—packages mapping tables and annotation databases as versioned, locally installable artifacts, which is a strong reproducibility model when package versions and genome builds are pinned and reported (37; 38). This packaging discipline aligns with contract thinking because it turns mappings into distributable objects rather than live queries, integrates naturally with *R* workflows where package versions are already provenance, and supports long-term reruns when computational environments are frozen at the package manager level.

At the same time, the dominant abstraction is still “select keys and columns from a point-in-time table”, so ambiguity is frequently surfaced as duplicated rows without an explicit semantic layer that distinguishes one-to-many biological history (merge/split) from one-to-many synonymy or from one-to-many transcript/protein relationships, and downstream code often resolves this implicitly. The limitation under drift is therefore not a lack of versioning but a lack of an explicit time axis: static mappings alone do not solve cross-release harmonization unless the version boundary is deliberately chosen and communicated across collaborating pipelines, and many packages do not represent branching *Ensembl* history events in a way that makes one-to-many outcomes first-class. Moreover, because these resources are typically scoped to a particular build and annotation snapshot, integrating datasets processed under different *Ensembl* releases or different assemblies still requires an explicit mapping policy that determines what is treated as equivalent, what is discarded, and how ambiguity is reported to downstream integration and QC. IDTrack targets this workflow niche in Python by making the snapshot boundary and historical window explicit, treating assembly as part of the coordinate system, and storing the resulting graph as a reusable artifact whose external namespace participation is controlled by a reportable inclusion policy. This design makes ambiguity and loss visible at scale through explicit outcome classes and diagnostics, which is particularly important in multi-dataset integration where silent feature drops and many-to-one collapses can change downstream model behavior without leaving an audit trace. In practice, *Bioconductor* -style packages remain an excellent approach when an analysis is R-native and the mapping boundary is fixed and shared, while IDTrack complements them by providing release-history time travel and audit-friendly ambiguity semantics in *Python* pipelines where cross-release harmonization is the dominant requirement.

### 4.11. BED as a cross-time mapping effort

*BED* (Biological Entity Dictionary) frames identifier conversion explicitly as a graph problem: it models biological entities and identifier relationships in a *Neo4j* graph data model and exposes R tooling to query and populate a local graph instance for a chosen organism and resource context (39). *BED* is motivated by practical integration pain points—heterogeneous identifier systems across resources, deprecated identifiers from former database versions, and the non-triviality of selecting a conversion path across multiple scopes—and it addresses these by explicitly encoding relationships in a local graph that can be queried for direct and indirect mappings rather than relying on single-source lookup tables. Because the model is implemented in a local *Neo4j* instance and the package includes import and caching utilities, *BED* can convert large identifier lists efficiently once the graph is populated, and its explicit substrate encourages users to treat mapping as traversal with inspectable intermediates rather than as opaque string replacement. Importantly, *BED* can represent deprecated identifiers and cross-resource links as first-class nodes, which is conceptually aligned with drift-aware thinking because retired identifiers remain part of the mapping space rather than being silently discarded.

At the same time, *BED*’s design goal is broad cross-resource dictionary construction within a user-defined context, rather than enforcing *Ensembl* release history as the explicit time axis with a snapshot boundary that can be encoded and rerun as a single parameterized artifact. In practice, drift-aware reproducibility therefore depends on the provenance of the constructed graph (which resource versions were ingested, how deprecated IDs were retained, and what update cadence was applied) and on the path-ranking policy used to choose between alternative routes, because a graph rebuilt from updated sources can change conversion behavior even if the query code is unchanged. Because *BED* is delivered as *R* tooling on top of a graph database, it is well suited for workflows that already operate in that ecosystem, but operational integration into pure-*Python* pipelines can require additional coordination around database lifecycle, environment pinning, and cross-language provenance capture. IDTrack differs by treating *Ensembl* releases as the explicit time axis, snapshot-bounding the backbone history at `snapshot_release` (with an explicit historical window start), controlling external participation via a reportable inclusion policy, and returning standardized one-to-zero/one-to-one/one-to-many outcomes with optional audit paths so ambiguity is surfaced rather than implicitly resolved. In other words, both tool families acknowledge that identifier conversion requires explicit context and can benefit from graph-based modeling, but IDTrack optimizes specifically for *Ensembl* time travel, assembly-aware reconciliation, and standardized outcomes that can be audited and reported in large *Python* integration workflows.

### 4.12. BridgeDb as an abstraction over mapping services

BridgeDb provides a standardized abstraction over identifier mapping services for genes, proteins, and metabolites, separating interface from backend so workflows can swap mapping resources without rewriting downstream code and enabling integration into pathway and network analysis tooling where identifier normalization is a recurring prerequisite (40). This separation is valuable infrastructure, particularly when teams need a uniform way to access multiple identifier systems and when mapping backends evolve, because it encourages modularity, promotes reuse across tools, and reduces the tendency for ad hoc converters to proliferate across pipelines. At the namespace layer, registry efforts such as *Identifiers.org/MIRIAM* provide persistent namespace identifiers and resolvable URIs for cross-resource referencing, improving interoperability at the level of naming and linking even when the underlying mapping services differ (41).

The drift-aware boundary is that abstraction and transitivity are not, by themselves, reproducibility contracts: without an explicit snapshot boundary, assembly context, and ambiguity policy, an abstracted mapping call can silently change as underlying services or mapping files change, and high-degree identifiers can dominate transitive connectivity in ways that are difficult to audit. For conversion-heavy integration tasks, this means that a *BridgeDb*-style interface still needs to be paired with rigorous provenance capture (exact mapping database/file versions, endpoint hosts, filtering rules, and row-reduction policy) if results are to remain comparable across time and across compute environments. IDTrack therefore constrains search semantics by release and node-type rules, elevates configuration into a versionable inclusion policy, and returns explicit ambiguity outcomes (and optional audit paths), treating the mapping substrate itself as a publishable artifact rather than an implicit property of a service layer. Practically, *BridgeDb* remains valuable for workflows whose core requirement is interoperable access across many identifier domains, while IDTrack targets settings where the scientific requirement is to publish a stable, rerunnable mapping substrate under *Ensembl* release drift and assembly heterogeneity, which is why this manuscript treats ecosystem comparison as capability context rather than as a benchmark race.

### 4.13. Synthesis: ecosystem value and IDTrack’s niche

Across the ecosystem, the dominant design centers are legitimate: some tools optimize for interactive lookup against authoritative live services, others optimize for analysis-first workflows where conversion is a helper step, others distribute versioned annotation tables for local pinning, and others provide interoperability interfaces that decouple clients from backends, so the absence of an explicit mapping contract is often acceptable when time and assembly assumptions are implicit and the analysis is not intended for cross-study comparability. The scientific gap emerges when identifiers become part of a reusable feature space—atlas integration, meta-analysis, cross-omics modeling, clinical reporting, large-scale knowledge-graph construction, and any long-horizon reuse of published matrices—because in those settings the mapping layer is no longer an internal convenience but a methodological transformation whose boundary choices determine what “the same gene” means across datasets and across years. IDTrack is positioned not as a replacement but as a reproducibility primitive for this subset of workflows, making the coordinate system explicit (namespace × release × assembly × entity layer), snapshot-bounding the evidence used for mapping, and promoting external namespace inclusion into a versionable inclusion policy rather than an implicit default.

The differentiator is the unit of publication: IDTrack is designed to publish a mapping substrate (snapshot-bounded graph artifact plus configuration identity) together with standardized outcome semantics (one-to-zero/one-to-one/one-to-many and optional audit paths), so mapping decisions can be reviewed, rerun, and cited with the same discipline expected for reference genomes, software versions, and statistical models. This contract framing clarifies composition with the ecosystem: point-in-time services remain excellent for exploratory lookup and annotation retrieval, enrichment platforms remain appropriate for functional interpretation once a stable identifier space is declared, and registry efforts remain valuable for naming interoperability, while IDTrack provides the pinned coordinate system that prevents upstream drift from masquerading as downstream biological variation. In applied terms, the same contract object supports legacy gene-list rescue, pre-integration harmonization across many studies, cross-database interoperability in collaborations, and time-consistent feature engineering for machine-learning workflows, because each reduces to publishing and rerunning an explicit mapping transformation rather than repeating an ad hoc lookup. The reviewer-facing claim becomes concrete: given the archived snapshot graph and its configuration identity, together with the declared snapshot boundary and window, assembly context, and ambiguity strategy, the mapping layer can be reconstructed exactly and its uncertainty semantics inspected, making drift a reported methodological axis rather than an invisible source of irreproducibility.

## 5. Assumptions and Technical Limitations

This section states assumptions and limitations as disciplined scope boundaries, because a mapping contract is only meaningful when its evidence substrate, coordinate system, and failure modes are explicit rather than implied. The intent is to make reviewability easier: readers should be able to distinguish upstream incompleteness, deliberate contract constraints, and algorithmic bounds, and to interpret 1→0 and 1→n outcomes as semantic outputs rather than undifferentiated failures.

### 5.1. Conceptual limitations: mapping is not ground truth

Identifier mapping in IDTrack is a transformation of symbolic coordinates rather than a claim about biological truth, and its “correctness” is therefore defined relative to a declared evidence substrate (snapshot-bounded *Ensembl* history plus an explicit external inclusion contract) rather than relative to a timeless ground truth that would require sequence-level and locus-level reasoning. Drift is partly irreducible because it reflects changing evidence, revised gene models, and shifting entity boundaries, so the existence of multiple plausible outputs is often the scientifically correct consequence of a well-bounded contract rather than an error to be hidden by post-hoc row reduction. In *Ensembl*, stable identifiers and version suffixes track curation decisions across releases, so the history graph legitimately encodes branching (splits, merges, retirements, reassignments, and version increments), and therefore 1→n and n→1 outcomes can be faithful reflections of upstream lineage semantics rather than pathologies of the traversal algorithm. IDTrack’s core claim is therefore conditional: given an explicit coordinate system (release window, snapshot boundary, primary assembly, and allowlisted externals) it reports what is reachable under node-type and activity constraints, and any interpretation of the result as “the” identity of a biological entity requires additional domain reasoning that the software does not and cannot supply. Under this framing, the choice between `strategy=‘best’` and `strategy=‘all’` is not an accuracy toggle but an ambiguity-policy statement, because collapsing branching history into a single target is a downstream decision about how uncertainty is propagated into analysis objects, while returning the full set is the conservative representation of what the contract permits. Scoring and tie-breaking are deterministic but not probabilistic, and they are designed to make decisions reproducible rather than to express calibrated confidence, which is why path selection reduces to a lexicographic preference over fewer assembly-priority drops, fewer external jumps/steps, and better-supported lineage edges rather than to a learned or statistical model of correctness. Conversely, a 1→0 outcome is also conditional: it means “not reachable under this snapshot boundary and configuration” rather than “nonexistent”, and it can arise from excluded releases, excluded externals, upstream gaps, or deliberate pruning decisions (for example, ignoring hyperconnective hubs), which call for different scientific responses rather than a single undifferentiated failure label. IDTrack therefore treats explicit reporting as the conservative response, but it cannot eliminate ambiguity encoded in upstream resources, and it cannot substitute for study-specific decisions about how ambiguous cases should be handled in downstream modeling, statistical testing, interpretation, or atlas-scale feature definition. This is why the API separates “not found in the snapshot” (no corresponding node under the declared contract) from “found but not convertible” (no admissible path under node-type, time, and assembly constraints) and from “convertible on the backbone but missing a requested external target”, because these are distinct methodological failure modes with distinct implications for reproducibility and for what a reviewer can conclude. In practice, credibility is strengthened when ambiguous cases are treated as first-class outputs that are quantified, inspected, and handled by declared policy, because under drift the least defensible posture is silent coercion of 1→n into a single token that is then reported as a stable feature space. As an honest boundary, this also implies that when a workflow’s primary requirement is unconditional uniqueness (a single identifier per input under all circumstances) or when a point-in-time lookup with minimal provenance is sufficient, lighter converters and endpoint-native services can be more appropriate, whereas IDTrack is optimized for the workflows that are willing to treat ambiguity and loss as explicit, reportable trade-offs in exchange for a rerunnable mapping contract.

### 5.2. Scope boundaries and representation assumptions: what is modeled (and what is not)

IDTrack’s scope is the reproducible transformation of identifier coordinates across the joint space of namespace × *Ensembl* release × genome assembly, and the central design choice is to materialize that space as a snapshot-bounded graph artifact rather than to rely on live queries against moving endpoints whose semantics can change without leaving an audit trace. Consequently, IDTrack is intentionally *Ensembl*-centric in its time semantics: time travel is defined over *Ensembl* releases, lineage edges are derived from *Ensembl* core history tables (`stable_id_event` contextualized by `mapping_session`), and the tool does not attempt to impose a universal cross-resource timeline whose release schedules, archival policies, and curation semantics are not aligned. External identifiers are incorporated only insofar as *Ensembl* exposes them in its cross-reference layer for a given release and assembly, so a conversion “to an external database” should be interpreted as “to the external accessions that *Ensembl* linked under this *Ensembl* context”, not as a claim that the output is pinned to a specific external-database release or that it reflects external authority independent of *Ensembl*’s integration choices. The backbone is *Ensembl* genes by construction (`DB.backbone_form=‘gene’`), because gene-level entities are the dominant feature unit in many integration workflows, but this abstraction can be suboptimal when the scientific unit is transcript isoform identity, protein sequence identity, locus-level coordinate equivalence, or regulatory element identity where non-gene objects must be treated as primary rather than auxiliary. Transcripts and translations are represented to improve connectivity and auditability, yet IDTrack does not expose transcript-or protein-level time travel as a primary contract object because non-gene history trees are explicitly removed from the traversal substrate (see `GraphMaker.remove_non_gene_trees in idtrack/idtrack/_graph_maker.py`), so the intended unit of longitudinal reasoning is gene-level harmonization rather than isoform-level lineage reconstruction. Likewise, the core history tables include additional entity types (for example, `stable_id_event.type` can include `rnaproduct`) but IDTrack’s supported forms are explicitly limited to `gene/transcript/translation`, so users should not assume that every *Ensembl* object class, annotation track, or regulatory feature is representable in a snapshot. Assembly awareness in IDTrack is a coordinate constraint on identifier meaning rather than a substitute for sequence-level remapping, so the package is not a replacement for chain-file *liftOver*, alignment-based reconciliation, or variant normalization pipelines when the scientific question is about genomic intervals, variant coordinates, or sequence equivalence across builds. IDTrack also does not perform cross-species mapping (orthology/paralogy resolution), because the contract is defined within an organism-specific *Ensembl* release history and any cross-species projection requires additional biological models, nontrivial uncertainty quantification, and explicit orthology database versioning that lie outside the identifier-stability problem we target. The package is therefore deliberately opinionated about what it treats as admissible evidence for “sameness” across time: it treats stable-ID lineage and explicitly enabled cross-references as evidence, and it does not attempt to infer equivalence from sequence similarity, genomic overlap, semantic similarity of names, or downstream functional annotations. These boundaries imply an honest division of labor: when the goal is a point-in-time, single-endpoint conversion with minimal provenance requirements, lightweight mappers or pinned annotation packages can be more appropriate and operationally cheaper, whereas IDTrack is optimized for settings where the mapping layer must survive reuse, review, and re-execution as a published artifact under drift. Stated differently, IDTrack trades universality and maximal transitive connectivity for a contract in which the coordinate system is explicit, reproducibility is enforceable by caching and archiving, and ambiguity is made visible rather than absorbed into undocumented heuristics.

### 5.3. Identifier-system assumptions: syntax, versions, and name collisions

At the representation layer, IDTrack assumes that identifiers can be normalized to canonical node labels that exist in the snapshot graph, which is straightforward for *Ensembl* stable IDs but more fragile for human-facing names (gene symbols and aliases) whose semantics are context-dependent, can collide, can be reused, and often represent many-to-many relations to *Ensembl* gene models even within a single release. *Ensembl* node labels are stored as `<stable_id>.<version>` with a single delimiter (`DB.id_ver_delimiter=‘.’`), so inputs that omit a version, use alternative separators (underscore/hyphen), or contain extra delimiters are handled by bounded heuristic normalization (`TheGraph.node_name_alternatives`) but can still fail or, in rare cases, resolve to an unintended node when multiple spellings or multiple version policies coexist in a dataset. Because *Ensembl* identifiers are frequently distributed without version suffixes in downstream pipelines, IDTrack also connects versionless base IDs to versioned gene nodes when version information is present (via `base_ensembl_gene` nodes created in `GraphMaker`), but this convenience necessarily defines a coarser equivalence class across versions, so analyses that rely on precise version semantics should treat version-stripped inputs as lower-specificity evidence and should supply versioned IDs whenever available. More generally, organisms and releases can differ in whether stable identifiers carry versions at all, and IDTrack can operate under “without version” or synthetic “add version” policies (`DB.first_version` is used as a placeholder), which improves practical coverage but should be interpreted as a representation compromise rather than as an assertion that version semantics are biologically irrelevant.

For robustness in interactive workflows, resolution is case-insensitive and includes simple punctuation permutations, yet this convenience relies on strong invariants: lowercasing must not create collisions for canonical nodes (`TheGraph.lower_chars_graph` raises an error otherwise) and external nodes that collide only by case are merged deterministically at build time, so strictly case-sensitive namespaces are not faithfully representable without upstream curation. To avoid collisions between canonical external accessions and free-text synonyms, IDTrack encodes nodes derived from `external_synonym` under a dedicated prefix (`DB.synonym_id_nodes_prefix`) and retries resolution with this prefix, which improves robustness in practice but also means that synonym matching is limited to what *Ensembl* exposes in the chosen release window and should not be confused with a comprehensive alias ontology. Graph construction also records cases where an external identifier string coincides with a pre-existing *Ensembl* node (`misplaced_external_entry`) rather than silently merging them, but users should still be cautious when providing mixed lists that combine *Ensembl*-looking prefixes with external namespaces, because string-level coincidence does not imply semantic equivalence. When `from_release` is not provided, IDTrack infers plausible origin releases and travel direction from pre-computed activity intervals (`TheGraph.get_active_ranges_of_id` and `Track.should_graph_reversed`), but this inference is underdetermined for discontinuous identifiers, mixed-provenance lists, or identifiers that are simultaneously valid under multiple assemblies, so the scientifically safest practice is to supply explicit source metadata when available and to treat inferred origins as hypotheses rather than as ground truth. External namespaces introduce further assumptions: IDTrack treats the external identifier string as meaningful only in the context of a declaring database, so a bare token that is valid in multiple databases is ambiguous by construction, which is why external participation is governed by an explicit YAML allowlist rather than by implicit inclusion of all cross-references. Because externals can attach at different biological forms and with assembly- and release-scoped validity, a conversion to an external database is not a simple join but a constrained synonym search whose output set and internal confidence flag depend on the target release, the primary assembly, and whether release-agnostic fallbacks were invoked (`final_conversion_confidence`), so users should not interpret these values as calibrated probabilities. Practically, this means IDTrack is strongest when inputs are already near the *Ensembl* coordinate system (stable IDs or well-supported cross-references) and when the analysis is willing to treat unmapped or ambiguous identifiers as reported methodological outcomes, rather than expecting the tool to invent a unique answer for every string.

### 5.4. Upstream-data limitations: *Ensembl* history and cross-references are authoritative but incomplete

IDTrack’s outputs are bounded by the completeness and internal consistency of the upstream *Ensembl* evidence it encodes, and therefore missing or ambiguous mappings can reflect absent lineage links, incomplete cross-references, schema drift, or changing identifier semantics rather than algorithmic instability. History edges are assembled from `stable_id_event` (including `old_stable_id, new _stable_id, old_version, new_version, mapping_session_id, type`, and the *Ensembl*-documented “combined mapping score” `score`) and contextualized by `mapping_session` (including `old_db_name`, `new_db_name`, `old_release`, `new_release`, `old_assembly`, `new_assembly`), so the contract inherits both the power and the limitations of these curated tables. Because `mapping_session.old_release` and `new_release` are stored as short strings (VARCHAR) in the *Ensembl* core schema, release parsing and assembly normalization are necessarily defensive operations, and edge cases in naming, archival availability, or patch-level conventions can surface as missingness or require HTTPS/FTP-based retrieval rather than live MySQL access. Graph construction also assumes that mapping session is coherent within the queried window (for example, at most one row per new release for the same core database), which is a reasonable invariant for *Ensembl* core schemas but an explicit upstream dependency that can fail under archival or mirror inconsistencies. The upstream evidence also contains entity classes beyond IDTrack’s supported forms (for example, `stable_id_event.type` can include `rnaproduct`), so some lineage events visible in *Ensembl* are intentionally excluded from the snapshot graph and cannot be recovered by algorithmic tuning without changing the package’s representational scope. Because IDTrack enforces a discrete integer release axis for traversal and filters out certain nonstandard patterns (for example, stable-ID version 0 entries in some patch contexts), the snapshot may omit niche annotation objects that do not follow standard stable-ID conventions, and such omissions should be treated as coverage limits rather than as algorithmic failures. In practice, *Ensembl* history tables can contain edge cases that violate simple monotonic lineage intuitions (duplicate score rows, reassignment patterns where a stable ID appears to “come alive again”, and references to nodes absent from release-specific identifier lists). IDTrack implements conservative repairs to preserve graph invariants and to keep the contract traversable (ignoring duplicate weight rows, normalizing some birth/retirement patterns, and deleting nodes absent from all `ids` tables in the snapshot window; see warnings in `idtrack/idtrack/ _graph_maker.py`). These mitigations improve robustness and reproducibility, but they are not guarantees of biological correctness, because the tool cannot recover information that is absent upstream and it must sometimes choose between “drop an inconsistent record” and “allow a corrupted lineage to destabilize the contract”, with the former chosen to preserve auditability and deterministic behavior. External connectivity is similarly bounded by *Ensembl*’s cross-reference layer (`xref, object_xref, external _db, external_synonym, identity_xref`), meaning that external mappings reflect what *Ensembl* chooses to expose for a given organism, release, and assembly rather than an exhaustive reconciliation of all external resources and their context-dependent identifier semantics. Cross-references are often many-to-many and can include low-specificity hubs, so IDTrack treats external inclusion as a contract choice that can legitimately trade coverage for specificity through allowlisting and hyperconnectivity filtering rather than silently accepting maximal connectivity that would make ambiguity unreportable at scale. Because *Ensembl* schemas evolve, IDTrack includes compatibility handling for cross-reference metadata that change names across releases (for example, legacy `identity_xref` columns `target_identity/query_identity` versus modern *Ensembl*e_identity/xref_identity), but when upstream fields are missing the package must propagate missingness rather than fabricate a confidence signal. Finally, scalar weights carried on history edges (`stable_id_event.score`) are used by IDTrack only as deterministic selection metadata inside a preference ordering, and because score conventions differ across releases and do not cover external edges, these values should not be interpreted as calibrated measures of biological correctness. An implication is that upstream curation changes can change mapping outcomes even when IDTrack code is fixed, which is precisely why the snapshot boundary exists: it makes the upstream state a reportable, archiveable axis rather than an untracked moving dependency.

### 5.5. Algorithmic and computational limitations: bounded search, heuristic scoring, and scalability

IDTrack represents the snapshot as a directed multigraph implemented in *Python* (`networkx.MultiDiGraph`), whose internal adjacency model is a nested dict-of-dict-of-dict-of-dict keyed by node → neighbor → edge-key → edge-attribute-dict, and this choice makes the implementation inspectable and easy to audit but imposes nontrivial per-node and per-edge memory overhead compared to compressed sparse or database-backed graph representations. Edges carry not only scalar weights for *Ensembl* lineage transitions but also release- and assembly-scoped validity metadata (for example, per-edge `connection` dictionaries and derived `available_releases` sets), so memory footprint scales with both graph topology and metadata cardinality, and permissive allowlists or long release windows can produce large graphs even when the number of distinct queried identifiers is modest. For interpretability and determinism, IDTrack does not treat the graph as an unconstrained transitive closure problem, but as a constrained path query that respects node-type semantics, release activity intervals, and assembly-scoped edge activity, so a conversion is defined as “reachable under the declared contract” rather than “connected somewhere in a maximally expanded synonym network”. The pathfinder (`Track.path_search` in `idtrack/idtrack/_track.py`) performs depth-first traversal over *Ensembl* history edges and may “beam up” from non-backbone contexts through synonym paths via external nodes, and its worst-case search space can grow combinatorially under branching lineage and high-degree externals, which is why exploration is explicitly bounded by default limits (`DB.external_search_settings: jump_limit=2, synonymous_max_depth=2`) that intentionally trade completeness for tractability. IDTrack additionally detects hyperconnective external nodes (out-degree > `DB.hyperconnecting_threshold`, default 20; see `TheGraph.hyperconnective_nodes`) and can ignore them during synonym expansion, which is computationally necessary to prevent frontier explosion and semantically useful to avoid low-specificity hubs dominating connectivity, but it also implies that some mappings mediated primarily by such hubs are intentionally not pursued unless the contract is revised. Search is staged rather than exhaustive: `Track.get_possible_paths` first attempts backbone-only time travel with external jumps disabled, then progressively relaxes synonym depth and external jump limits only until at least one path exists (and only then, if needed, enables more permissive multiple-transition modes), so candidate enumeration is biased toward minimal external reliance and may omit alternative routes that are reachable only under more permissive settings. This staged design makes conversions fast and interpretable for common integration workloads, but it should be understood as a bounded search policy rather than as a guarantee that every mathematically possible path in the full graph was enumerated, and users who need maximal reachability must accept higher computational cost by widening the contract and increasing search limits. Candidate paths are ranked by deterministic heuristics that penalize assembly-priority drops and external dependence (see `Track.minimum_assembly_jumps` and the lexicographic ordering in `Track._path_score_sorter_single_target`). History-edge weights are then aggregated under a user-declared missingness policy (`remove na=‘omit’|’to 1’|’to 0’`), making `strategy=‘best’` a reproducible policy choice while also implying that “best” is defined by contract-consistent scoring rather than by a claim of biological optimality. Because external edges are not commensurately weighted with *Ensembl* lineage edges (external jumps are counted rather than scored) and because history weights are *Ensembl*-provided selection metadata rather than calibrated correctness measures, the reported confidence and ranking values should be interpreted as internal decision records that support auditability, not as probabilities of correctness. In degenerate cases where multiple paths are exactly tied under the scoring tuple, the selected “best” path can depend on deterministic but semantically arbitrary enumeration order, which is why `strategy=‘all’` and optional path return are the scientifically safer choices when a downstream analysis is sensitive to which representative path is chosen. At scale, the practical limitation is therefore not that IDTrack cannot be run in batch (the system is explicitly cache-first and supports bulk conversion and atlas harmonization), but that per-identifier cost can become high for highly ambiguous identifiers or overly permissive snapshots, so workflow design should control ambiguity channels (release window, allowlist breadth, and search limits) rather than treating “convert everything” as a free operation. Finally, convenience filters such as `deprioritize_lrg_genes=True` (which drops LRG * region-style targets when alternative mappings exist) encode gene-centric relevance assumptions that are sensible for atlas integration but may be inappropriate for reporting pipelines that treat LRG identifiers as primary endpoints, so such settings should be audited and, if necessary, disabled as part of the published contract. If a project’s primary requirement is extreme scale (tens of millions of nodes/edges), language-agnostic graph artifacts, or formal global-optimum guarantees under a fully expanded mapping graph, then a different backend (compiled graph libraries, database engines, or precomputed mapping tables) may be more appropriate, whereas IDTrack intentionally optimizes for auditability and contract-driven reproducibility in the regime most common for atlas integration and legacy-data rescue.

### 5.6. Coverage limitations: upstream availability and configuration

IDTrack’s practical coverage is constrained by upstream *Ensembl* availability (organism, assembly, and release windows) and by cross-reference coverage for enabled externals, so missing conversions can reflect absent upstream links, assembly scoping, archival gaps, or deliberate configuration choices rather than algorithmic errors. For the current distribution, default profiles are provided for human, mouse, and pig, and this limited set is deliberate because the package treats organism and assembly availability as a provenance contract (declared in `DB.assembly_mysqlport_priority` and `DB.supported_organisms` in `idtrack/idtrack/ _db.py`) rather than as an implicit promise that any identifier can be resolved without stating its coordinate system. Time coverage is bounded not only by user-chosen `ignore_before/snapshot_release` but also by upstream service constraints, because public MySQL mirrors expose only a limited historical depth per port (see `DB.mysql_port_min_release`), and older releases or assembly archives may require dump-based retrieval and may not be uniformly available across organisms. The snapshot boundary (`snapshot_release`) and the historical window start (`ignore_before`) jointly determine whether an older dataset’s coordinate system is representable, so choosing a boundary that excludes an assembly-handoff period or early release history deterministically increases 1→0 outcomes for legacy inputs, while choosing a wider window increases operational footprint and can amplify ambiguity channels by importing additional branching history. Assembly coverage is an explicit constraint: *Ensembl* assembly handoffs are species-specific and overlap only within certain release intervals, so selecting a primary assembly that does not overlap the relevant input history can make conversions impossible or can force reliance on external bridges that change ambiguity and must be reported as part of the contract. External coverage is contract-bound: the YAML allowlist specifies, per form and per assembly, which external databases are admissible at which releases, so a mapping can disappear either because it was never present upstream or because it was excluded by configuration, and in contract terms these cases should be reported differently. Because the allowlist is release-scoped, older releases can fall outside the configured YAML, and graph construction will then skip external edges for those releases while still building the *Ensembl*-only history substrate, which preserves time-travel functionality but reduces external reachability in precisely the historical regimes where legacy-data rescue is common. IDTrack also provides a non-default “include everything” posture (`narrow_external=False`) that attempts to incorporate all externals available in *Ensembl*, but this setting can dramatically increase graph size and ambiguity and is therefore best treated as an explicitly justified and reported choice rather than as a baseline configuration. The allowlist schema is intentionally simple (an explicit `Include: true/false` gate over release-scoped availability), which makes it reviewable but also means it cannot express fine-grained, study-specific scoring policies, database-specific synonym semantics, or curation-aware trust rankings beyond inclusion and exclusion. Because assembly codes are species-specific in *Ensembl* schema naming (for example, “38” denotes GRCh38 in human but denotes a different build number in other organisms), the correct scientific practice is to treat assembly as an organism-scoped parameter rather than as a globally meaningful integer. Extending coverage to a new organism is therefore an explicit engineering and curation task (add assembly/port configuration, generate and curate an external allowlist template, validate it against the target release window, and publish the resulting configuration), and this explicitness is part of the tool’s credibility posture rather than a limitation that should be hidden. Because allowlists and release windows are first-class inputs, changing them is methodologically equivalent to changing reference genomes or preprocessing pipelines, so analyses that claim comparability across studies should treat these changes as explicit protocol differences rather than silent configuration tweaks. If an analysis expects legacy releases, mixed-build inputs, or long-horizon reuse, then the snapshot window and primary assembly must be chosen accordingly (and reported), because choosing a boundary that excludes relevant history is equivalent to choosing a coordinate system that cannot represent the data.

### 5.7. Operational limitations: resources, endpoints, and caching

Graph construction requires network access (public MySQL mirrors when reachable, HTTPS/FTP release dumps when ports are blocked) and sufficient disk space to cache downloaded tables and derived artifacts, and while caching makes repeated conversions fast, it also means that the local repository becomes part of reproducibility metadata that must be managed, backed up, and referenced in publications. Operational availability is itself a dependency: public endpoints can change, be rate-limited, or be temporarily unreachable, and building snapshots on restricted compute environments can require explicit network routing or building the artifacts on a networked machine and then relocating them, neither of which changes mapping semantics but both of which affect reproducibility practice. In practice, initial builds can take minutes to hours depending on organism, release window, enabled externals, and network throughput, and the documentation’s baseline expectations (roughly 2–5 GB disk for cached databases and ≥ 4 GB RAM, with ∼ 8 GB RAM preferred for larger graphs) reflect the fact that the snapshot graph is an in-memory NetworkX object with nontrivial overhead. Because IDTrack stores cached tables in HDF5 (h5py) and persists the graph as a *Python* pickle, reproducibility across compute environments depends on a functioning scientific *Python* stack with compatible HDF5 libraries, which can require additional system-level setup compared to pure-text mapping tables and can therefore be a nontrivial deployment consideration on some platforms. HDF5 supports robust single-writer/multiple-reader patterns but not transparent concurrent multi-writer updates, so parallel snapshot builds targeting the same `<organism> assembly-<assembly>.h5` file or the same local repository can lead to contention or corruption unless explicit locking and isolation policies are used. The conservative practice is therefore one build process per local repository with explicit artifact promotion into shared storage, which aligns with the contract goal (stable, publishable artifacts) but should be understood as an operational constraint when designing atlas-scale pipelines. The pickle artifact is *Python*-specific and reconstructing complex objects requires compatible import paths and library versions, and because unpickling arbitrary data is not safe, graphs should be treated as trusted, local artifacts rather than as interchange formats for untrusted distribution. For long-horizon archiving, the robust approach is to treat the pickle as a derived acceleration artifact and to archive the contract-defining inputs (cached tables, YAML allowlist, build parameters, and the IDTrack software version) that allow the snapshot to be rebuilt under the same contract coordinates, because these inputs are the scientific substrate while the pickled object is an implementation convenience. Under the contract model, this is a feature rather than a burden, because the point is to turn a historically moving dependency into a stable on-disk artifact that can be rerun and audited, but it implies that projects must treat cache integrity (and cache identity) as part of Methods reporting rather than as a local implementation detail. Operationally, this creates a reproducibility posture choice: a user-scoped cache optimized for convenience is rarely sufficient for publication-grade reuse, whereas a project-scoped local repository that is archived (or at least hashed) alongside analysis code creates a reviewer-checkable handle on the mapping substrate and makes “what mapping did you actually run?” answerable years later. This posture also requires users to understand that graph snapshots are not purely keyed by organism and release window but also by assembly choice and external allowlist contents, so changing primary assembly or revising the YAML (even without changing code) is a scientifically meaningful change that typically requires rebuilding the snapshot and reporting the new contract coordinates. As a practical constraint, the exported graph filename currently encodes organism and release-window settings but not the chosen primary assembly or allowlist identity (see `GraphMaker.create_file_name` in `idtrack/idtrack/ _graph_maker.py`), so maintaining multiple snapshots side-by-side requires separate local repositories or explicit artifact management to prevent accidental reuse of the wrong contract. There is also an unavoidable resource trade-off between snapshot breadth and operational footprint: wider release windows, additional assemblies, and broader external allowlists increase graph size and can increase build time and memory use, while narrower snapshots reduce connectivity and can increase 1→0 outcomes or multiple-target ambiguity by construction, so responsible use involves choosing a boundary that matches the scientific need rather than defaulting to maximal inclusion. When compute nodes lack direct internet access, operational routing solutions (for example, ConnectionBridge) can restore the ability to build snapshots by proxying network requests, but this is infrastructural support rather than scientific novelty and it does not change the requirement that the snapshot boundary and allowlist contract must be published for results to be interpretable. Once a snapshot is built, conversions can proceed from local artifacts without further endpoint access, which is precisely the reproducibility posture the package is designed to enforce, but it also means that teams should plan for long-horizon artifact storage rather than assuming that upstream endpoints will remain stable. Finally, IDTrack also ships optional external-mapping backends (for example, *MyGene.info, g:Profiler, BioMart*) for convenience, but these rely on third-party endpoints whose versioning may be opaque or time-dependent, so they should be treated as exploratory utilities rather than replacements for the snapshot-bounded contract when publication-grade reproducibility is the goal.

### 5.8. Related-work comparison limitations

Capabilities of external tools depend on endpoints, versions, and user configuration, so this supplement avoids numerical benchmarking and instead compares along explicit axes (time and assembly semantics, ambiguity reporting, auditability, and versionable configuration) that remain meaningful under drift, which is not an argument that other tools are wrong but a statement that they were designed for different integration postures. Because many tools can be made more reproducible by manual caching and careful provenance capture, the relevant distinction in this manuscript is whether a tool makes the mapping contract first-class and reportable by design rather than whether a mapping exists in principle for a given identifier list. In drift-aware settings, a benchmark without declared boundaries is not only unfair but ill-defined, because the notion of a timeless “ground truth” mapping across releases collapses once entity definitions change and once different tools silently choose different boundary defaults. Moreover, “performance” is not a stable property when the mapping substrate itself is moving, because changes in upstream cross-reference coverage, release history, or endpoint behavior can shift both outcome distributions and computational cost without any change in a tool’s code, so comparative claims must be anchored to published version and endpoint coordinates to be scientifically interpretable. When reviewers or readers require tool-specific details, those details should be interpreted as scope descriptions rather than universal claims, and point-in-time behavior should be reported with endpoint/version boundaries and configuration snapshots for reproducibility rather than narrated as a timeless property of the method. In particular, a paper that uses any mapper should report what was queried, when (release/version and access date), under which assembly assumptions, and with what ambiguity policy, because otherwise the mapping step remains a hidden dependency regardless of which software produced it. Within that reporting posture, IDTrack’s primary contribution is not a claim of universal superiority but a way to make boundary choices explicit, rerunnable, and inspectable in *Python* workflows where cross-release integration is a dominant source of irreproducibility.

## 6. Extended Methods

### 6.1. Methodological roadmap, notation, and reproducibility surface

This Extended Methods section is written as a code-anchored specification of IDTrack’s mapping contract, with the explicit aim that every conversion claim in the manuscript is reconstructible from archived artifacts and a declared policy even when upstream endpoints and cross-reference coverage have drifted since publication. Because identifier conversion under drift is a coordinate transformation rather than a timeless lookup, we present the method as a small set of explicit coordinate tuples whose values are fixed either at build time (graph snapshot) or at query time (search and selection policy), so that reproducibility is enforced mechanically rather than delegated to narrative.

Method structure (contract coordinates and objects).

We separate a build-time contract 𝒞_build_ that defines the snapshot graph and its artifact identity from a query-time contract 𝒞_query_ that defines admissible traversals and deterministic selection, because these two layers jointly determine all mapping outcomes and must be reportable as Methods metadata.

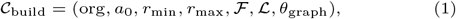

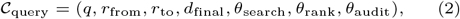

where org is the *Ensembl* organism name, a_0_ the primary assembly, [r_min_, r_max_] the inclusive release window, ℱ the included identifier forms, ℒ the external allowlist, and the θ blocks are the explicit bounds and ordering rules implemented in code (`idtrack/idtrack/_db.py`). Under this formalization, a study is reproducible if and only if the artifacts that materialize 𝒞_build_ (*HDF5* caches, allowlist YAML, and the serialized graph snapshot) are archived or re-creatable under a fixed software version, and a conversion is interpretable if and only if the values in 𝒞_query_ are reported rather than absorbed into implicit defaults.

Component map (what must be reproduced, in what order).

1. **Local repository and artifact identity**. All builds and queries are bound to a writable local repository whose on-disk layout is part of the scientific method because it is where the drift-sensitive substrate is frozen into inspectable files rather than being re-derived from moving endpoints (Section 6.2). Reproducibility therefore reduces to artifact identity (filenames, hierarchical keys, and content hashes) instead of relying on informal statements such as “queried *Ensembl*” whose semantics move with time.
2. **External allowlist policy**. External namespaces are opt-in via a versionable YAML allowlist that specifies inclusion by organism, form, assembly, and release, converting the connectivity-versus-ambiguity trade-off into an explicit, reviewable part of the contract rather than an implicit property of a queried endpoint (Section 6.3). The same mechanism also makes synonym-derived externals (from *Ensembl* `external_synonym`) explicit by encoding them as a separate namespace prefix, preventing semantic conflation between stable accessions and free-text labels (Section 6.5).
3. **Release window, assembly semantics, and acquisition invariance.** Snapshot construction is defined by an inclusive release window and a primary genome assembly, and acquisition is treated as contract-equivalent across *MySQL* mirrors versus HTTPS/FTP dumps once cached locally, so build semantics are invariant to infrastructure constraints (Section 6.4). Multi-assembly inclusion is explicit and conservative, because the graph must represent historically relevant identifiers without duplicating entire backbones for assemblies that are not required by the declared window and external policy.
4. **Core graph data structures** (schema-level semantics). The snapshot substrate is a typed directed multigraph whose nodes represent identifiers and whose edges represent either time-axis history transitions or release/assembly-scoped cross-namespace relations, with edge payloads designed to make temporal validity and evidence provenance machine-checkable (Section 6.5). These data-structure decisions are chosen to make time and namespace reasoning composable in a single substrate while keeping query-time checks reducible to constant-time set membership rather than ad hoc joins.
5. **Graph construction algorithms (deterministic, cache-backed)**. GraphMaker constructs history edges from `stable_id_event` and **`mapping_session`**, composes per-form graphs, adds cross-form and external edges with release validity metadata, normalizes casing collisions, and serializes the result under a deterministic filename encoding the contract boundary (Section 6.5). Build-time determinism is treated as a scientific requirement because it ensures that two users who share the same cached tables and configuration obtain byte-identical behavior at query time.
6. **Query optimization and cached summaries.** Deterministic cached properties (reverse view, combined edge dictionaries, activity ranges, hyperconnective hubs, and origin trios) act as derived indexes that shift repeated query-time checks from neighbor scans to dictionary/set lookups without changing semantics given a fixed snapshot (Section 6.6). These caches are computed as pure functions of the serialized snapshot, which preserves the contract posture while making atlas-scale workloads tractable.
7. **Identifier resolution and mapping (pathfinder)**. Conversions are posed as a constrained, time-respecting path enumeration on a typed identifier graph, with staged relaxation and bounded external “beam-up” jumps that introduce connectivity only when the backbone cannot explain a query, and with deterministic ranking so repeated runs under the same contract produce identical admissible explanations. The API then returns outcome classes explicitly (1→0, 1→1, 1→n) rather than silently coercing a single answer, so ambiguity and loss are propagated as first-class evidence under a declared policy rather than being hidden in downstream preprocessing (Section 6.7). The search and ranking parameters are part of θ_search_ and θ_rank_, so changing them is a methodological change rather than an implementation detail.
8. **API schema, integration, and downstream ledgers.** The public API surfaces machine-readable results whose fields preserve failure and ambiguity semantics across interactive, batch, and dataset-harmonization workflows, enabling deterministic QC and reporting without re-implementing mapping logic in downstream code (Section 6.11 and Section 6.12). This separation makes the mapping layer behave like an auditable instrument whose outputs can be aggregated, compared, and archived as structured evidence.
9. **Validation and quality control**. A developer-facing invariant test harness (**`TrackTests`**) and build-time consistency checks enforce that the constructed snapshot encodes a coherent time axis and that derived caches agree with raw graph structure, which matters because reproducibility requires detecting substrate anomalies rather than absorbing them silently (Section 6.10).

Notation (formal objects).

We denote by r_min_ = *`ignore_before`* and r_max_ = *`snapshot_release` = `ignore_after`* the inclusive release window that defines the snapshot boundary, and by ℛ = {r ∈ ℤ : r_min_ ≤ r ≤ r_max_} ∩ `g.graph[‘confident for release’]` the set of *Ensembl* releases explicitly represented in the built snapshot. We write the snapshot graph as a typed multigraph 𝒢 = (V, E_H_ ∪ E_X_, τ) where τ : V → 𝒯 is the node-type function (`DB.node_type_str`) and E_H_ and E_X_ are the history and cross-reference edge sets, each carrying explicit metadata described in Section 6.5. For complexity statements, we use N = |V |, M_H_ = |E_H_ |, M_X_ = |E_X_ |, and R = |ℛ|, and we distinguish operations that perform a single graph traversal (typically O(N + M_H_ + M_X_) over the reachable subgraph) from operations that enumerate admissible paths (worst-case exponential in the number of admissible branch points even when each local step is O(1) average).

Top-level pipeline (full-width pseudo-code; build once, query many).

### 6.2. Local repository contract (artifacts, cache keys, and provenance)

IDTrack binds every run to a writable local repository, because reproducible mapping under drift requires that upstream tables, the external inclusion policy, and the constructed snapshot graph are persisted as artifacts rather than treated as transient query results. Under the contract framing used throughout the manuscript, the local repository is the boundary where a mapping choice becomes a publishable object, because once the snapshot has been materialized as files, conversions become a deterministic computation over named artifacts rather than a moving dependency on external services.

Artifact inventory (what is cached).

The repository stores (i) external allowlist configuration files (**`<organism>_externals_template.yml` and `<organism>_exte rnals_modified.yml; idtrack/idtrack/_external_database s.py`)** and (ii) an organism/assembly-scoped HDF5 store of raw and derived *Ensembl* tables and xref joins (**`<organism >_assembly-<assembly>.h5; idtrack/idtrack/_database_ma nager.py`**). It also stores (iii) the serialized snapshot graph whose filename deterministically encodes the boundary and build flags (e.g. **`graph_<organism>_min<ignore_before>_max<ig nore_after>_narrow_forms…_extall.pickle; idtrack/idtrac k/_graph_maker.py:create_file_name`**). Downstream workflows may add project-scoped derived artifacts (e.g., harmonization ledgers persisted by **`HarmonizeFeatures; idtrack/idtrack/_har monize_features.py`**), but the reproducibility-critical minimum is the contract tuple recoverable from these three artifacts, i.e. (IDTrack version, org, a_0_, r_min_, r_max_, ℱ, ℒ, θ_graph_) plus the query-time policy fields used for the reported conversion claims.

**Algorithm S1.**
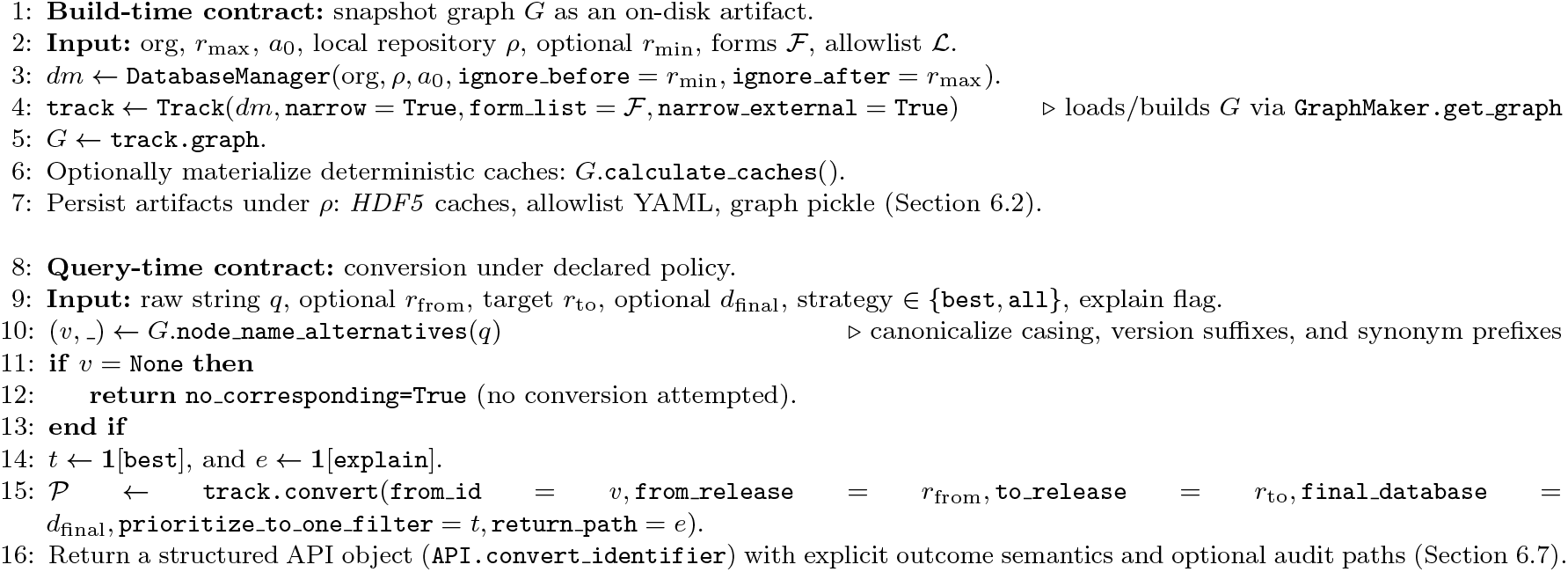
IDTrack separates a build-time contract that materializes the snapshot substrate from a query-time contract that fixes search and selection policy, making conversion reproducible as an artifact plus a declared coordinate tuple.

#### Deterministic cache keys (HDF5 schema)

Within **`<organism>_assembly-<assembly>`**.h5, cached frames are keyed by a deterministic hierarchy scheme returned by **`DatabaseManager.file_name`** whose string identity is a pure function of (r, **`df_type, df_indicator, form, usecols`**) together with the instance-level coordinates (org, a_0_, ρ) that determine the physical file path (**`idtrack/idtrack/_database_manage r.py`**). Concretely, raw MySQL tables are stored under keys of the form **`ensr_mysql`** <table> COL <c1> COL <c2>… (column subsets sorted and concatenated using COL), processed artifacts are stored as **`ensr_processed_<df_indicator>_<form>`**, and form-agnostic tables are stored as **`ensr_common_<df _indicator>`**, which makes cache identity fully inspectable without reading logs. DataFrames are serialized via a PyTables-free HDF5 round-trip layer implemented in idtrack/idtrack/_utils_hdf5.py that stores column labels, dtypes (including categoricals), index metadata, and data arrays as explicit datasets and attributes inside an HDF5 group, which is an implementation choice driven by reproducibility across environments where PyTables availability and dtype round-tripping are brittle.

#### Integrity and concurrency (single-writer discipline)

Because the HDF5 cache is a mutable scientific artifact, the conservative operational assumption is one writer process per **`<organism>_assembly-<assembly>.h5`** file and explicit isolation for parallel builds, since HDF5 supports single-writer/multiple-reader patterns but not transparent concurrent multi-writer updates (42). Similarly, the graph snapshot is serialized as a *Python* pickle (**`idtrack/idtrack/_graph_maker.py`**), which makes it a high-performance, *Python*-native cache for trusted local reuse but not a safe interchange format for untrusted distribution, so long-horizon archiving should treat the pickle as a derived acceleration artifact and the cached tables plus configuration as the contract-defining substrate.

#### Complexity and performance (artifact lookup versus computation)

Once the HDF5 and graph snapshot artifacts exist, subsequent runs are dominated by local I/O and in-memory traversal rather than by network access, so the asymptotic behavior is controlled by N, M_H_, and M_X_ rather than by endpoint latency and is therefore interpretable as a property of declared contract boundaries. HDF5 hierarchy lookup is a bounded number of hash-table operations (string-keyed group/dataset lookups; amortized O(1) average under standard assumptions (43)), while the dominating cost is I/O proportional to serialized table size and the conversion algorithm cost described explicitly in Section 6.7, which together make performance claims mechanically attributable to artifacts and bounds rather than to informal runtime anecdotes.

### 6.3. External allowlist contract (YAML schema, lifecycle, and ambiguity control)

External namespaces can reconnect disconnected *Ensembl* histories but they also create ambiguity channels, so IDTrack treats external participation as a reviewable contract rather than as an implicit inclusion of all cross-references available on a given endpoint. Concretely, this contract is stored as a YAML allowlist (**`<organism>_externals_modified.yml`**) located under the local repository when configured, with a packaged fallback under **`idtrack/idtrack/default_config`**/ when absent, making external participation both configurable and recoverable (**`idtrack/idtrack/_external_databases.py`**).

Formal object (allowlist as an explicit coordinate constraint). We treat the allowlist as a partial function ℒ that maps a coordinate quadruple (org, f, d, a)—organism, form, external database name, and assembly—to an inclusion flag and an allowed release set, so the legal external edge set is defined by the contract rather than by whichever cross-references happen to be returned by an endpoint at query time. Equivalently, the allowlist defines the predicate

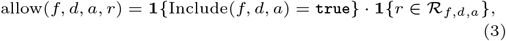

where ℛ_f,d,a_ ⊆ ℛ is the set of releases listed under that YAML entry, and any edge whose provenance tuple fails this predicate is excluded from the snapshot by construction.

#### Schema (exact hierarchy and semantics)

The YAML schema is hierarchical by design—**`organism → form → name db`**—and each external database entry carries a stable **`Database Index`**, a reserved **`Potential Synonymous`** field, and an Assembly block that maps assembly codes (as strings) to an explicit release list and an **`Include: true/false flag (idtrack/idtrack/_external_databases.py`**). This structure is methodological rather than cosmetic, because external availability is not constant across the time axis or across assemblies, so a scientifically meaningful allowlist must specify not only which databases participate but also under which coordinate context they are legally allowed to contribute edges to the snapshot.

#### Synonym namespaces (separating accessions from free-text labels)

*Ensembl* cross-references include both accession-like identifiers and synonym-like labels from **`external_synonym`**, and IDTrack encodes these evidence types as distinct namespaces by prefixing synonym IDs with **`DB.synonym_id _nodes_prefix`** and also prefixing their name db field in the extracted external table (**`idtrack/ idtrack/_database_manager.py:create_external_db`**). This design ensures that a raw string such as **`ACTB`** can be resolved deterministically as either a synonym node (**`synonym_id::ACTB`**) or as an accession from a different database, preventing accidental conflation while still allowing synonym nodes to act as bounded bridges during pathfinding under a declared allowlist policy.

Lifecycle (how a study should curate and report it). Operationally, the workflow is to generate a template that enumerates every observed external database by release and assembly (**`ExternalDatabases.create_template_yaml`**) and then curate a modified file by setting Include: true for the subset required by the study. As *Ensembl* releases evolve, the curated file can be validated for currency (**`ExternalDatabases.validate_yaml_file_up_to_date`**) and, once finalized, it becomes a deterministic input to graph construction (**`ExternalDatabases.load_modified_yaml`** and downstream **`DatabaseManager.get _db(‘external_relevant`**‘); **`idtrack/idtrack/_database_manager.py`**). The methodological consequence is direct: publishing the allowlist file (or at least its content hash and retrieval path) is publishing part of the mapping contract, because changing this file changes which edges exist, which paths are admissible, and therefore what “coverage” and “ambiguity” mean under the declared snapshot boundary.

Enforcement (why two runs differ only when the contract differs). At graph-build time the allowlist is enforced mechanically rather than narratively: the external cross-reference extraction is filtered to databases whose allowlist entry is **`Include: true`** for the relevant assembly and release, and **`GraphMaker.construct_graph`** then adds only those external nodes and edges, meaning that two runs that differ only in allowlist identity differ in reachable mapping space by construction (Section 6.5). For completeness, IDTrack exposes an “include everything” mode (**`narrow_external=False`**) that attempts to incorporate all external databases exposed by *Ensembl*, but this mode is treated as non-default because it grows M_X_ and the number of ambiguous paths, and therefore undermines the contract objective unless a study explicitly justifies and reports that choice.

#### Complexity and ambiguity control (why this belongs in Methods)

Because cross-reference extraction is filtered by **`name_db`** membership in the allowlist, the allowlist directly controls M_X_ and therefore controls both the branching factor of synonym expansion and the empirical distribution of 1→n outcomes, which is why we treat it as part of the scientific contract rather than as a convenience configuration. In practice, reporting ℒ is equivalent to reporting which joins are permitted in the identifier graph, and making that join policy explicit is precisely what converts a mapping step from an implicit preprocessing artifact into a reproducible, reviewer-auditable method.

### 6.4 Release window and assembly semantics (DatabaseManager view)

Snapshot construction is defined by an inclusive release window [r_min_, r_max_] and a primary assembly context, because *Ensembl*’s identifier semantics are indexed by (release, assembly) and therefore any conversion step that leaves these coordinates implicit is underspecified under drift. Concretely, **`API.build_graph(organism_name, snapshot_release, genome_assembly`**) instantiates a **`DatabaseManager`** with **`ignore_after=snapshot_release`** and an **`ignore_before`** default chosen as the earliest release supported by configured access methods (implemented as **`min(DB.mysql_port_min_release.values())`**; **`idtrack/idtrack/_api.py`**), so r_max_ is a declared boundary rather than a moving “latest” endpoint.

#### Assembly priority and multi-assembly inclusion

Assembly semantics are encoded in configuration rather than assumed implicitly: **`DB.assembly_mysqlport_priority`** specifies, per organism, which assemblies exist, which ports or dump resources can serve them, their priority order, and their release ranges, allowing IDTrack to declare a primary assembly for output while still retaining additional assemblies in the snapshot when they are historically necessary for reachability (**`idtrack/idtrack/_db.py`**). This matters in integration pipelines because mixed-build inputs are common (e.g., legacy GRCh37 identifiers mixed with GRCh38), and without explicit assembly handling a conversion step can silently collapse incompatible coordinate systems into a single output while erasing provenance, whereas IDTrack preserves assembly information at the node-type and edge-activity level and treats the chosen primary assembly as an output preference rather than as an excuse to discard assembly provenance.

#### Data acquisition under infrastructure constraints

Data acquisition is designed to be robust to infrastructure constraints while keeping semantics stable: **`DatabaseManager`** prefers public *MySQL* when reachable but falls back to HTTPS/FTP dump discovery when ports or archives are blocked. It caches both raw tables (e.g., **`stable_id_event, mapping_session, relationcurrent`**) and derived artifacts (e.g., **`idhistory`**) under deterministic HDF5 keys so repeated builds are contract-stable once artifacts exist locally (**`DatabaseManager.get_table, DatabaseManager.get_db, DatabaseManager.file_name; idtrack/ idtrack/_database_manager.py`**). Release discovery is therefore treated as part of the reproducibility substrate rather than as a deployment detail: **`DatabaseManager`** queries *Ensembl* REST for the latest release and caches it process-wide, probes core-schema availability across configured assemblies/ports, and resolves older schema name irregularities (e.g., patch-letter suffixes) so a declared r_max_ remains buildable even years later when live services have advanced.

#### Schema normalization across releases (preventing hidden parsing drift)

Because *Ensembl* schemas evolve, **`DatabaseManager`** normalizes known table-level differences during extraction, for example by detecting legacy **`identity_xref`** column names (target identity/query identity) and renaming them into the contemporary convention (***Ensembl*e_identity/xref_identity**) so that downstream joins are invariant to release-specific schema drift (**`idtrack/idtrack/_database_manager. py: create_external_db`**). Similarly, join-key columns are coerced to numeric types and malformed continuation lines are dropped when encountered in HTTPS/FTP dumps, because reproducibility requires that “the same release window” yields the same extracted relations instead of being sensitive to transport-specific parsing artifacts.

External mapping table as a reproducible relational object. For each release *r* and form *f*, external mappings are materialized by **`DatabaseManager.create_external_db`** as a de-duplicated relation.

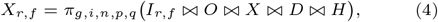

where *I*_*r,f*_ denotes the form-specific *Ensembl* identifier table at *(r, f)* (`ids`), *`O = object_xref, X = xref`, D* = **`external_db`**, H = **`identity_xref`**, and (*g, i, n, p, q*) correspond to **graph_id, id_db, name_db, *Ensembl*e_identity**, and xref identity. This relation is augmented by a synonym union term derived from **`external_synonym`** in which synonym display labels are prefixed and treated as a distinct namespace (Section 6.3), so that the eventual graph edges can record provenance and validity as explicit set-valued metadata rather than as implicit join logic.

#### Complexity (build cost is dominated by bounded tables, not by hidden heuristics)

The computational cost of acquisition is dominated by downloading and serializing bounded-size tables for each release in ℛ, so the asymptotic cost is O(Σ_r∈ ℛ_ |T_r_|) for table row counts plus I/O proportional to cache size, while the semantic definition of the snapshot remains controlled exclusively by (r_min_, r_max_, assembly, allowlist) rather than by adaptive heuristics. Because the method caches every retrieved and derived artifact under deterministic names, re-runs do not silently change unless a boundary-defining parameter changes, which is precisely the behavior required when a conversion claim is treated as a coordinate-dependent scientific statement rather than as an ephemeral lookup.

### 6.5 Graph schema and construction (TheGraph & GraphMaker)

The snapshot substrate is a typed directed multigraph **`TheGraph (a networkx.MultiDiGraph; idtrack/idtrack/_the_graph.py`**) in which every node is an identifier with an explicit node type and every edge encodes either time-axis history or a release/assembly-scoped cross-namespace relation, because the scientific object is not a timeless ontology but a time-parameterized mapping substrate whose boundary choices are inspectable and whose semantics are stable under archival. This data structure choice is pragmatic: NetworkX ‘s **`MultiDiGraph`** exposes adjacency as a nested mapping keyed by u ↦ v ↦ k ↦attribute-dict (44; 45), so direct adjacency and attribute lookup are amortized O(1) average while neighborhood iteration is O(deg(u)), and the multi-edge structure is essential because *Ensembl* history contains genuine branching events that cannot be represented as a simple function without losing provenance.

#### Formal schema (node and edge typing)

We view 𝒢 as *(V, E_H_ ⋃ E_X_, τ, α_V_, α_E_)* where *τ (v) ∈ 𝒯* is stored as **`v[DB.node_type_str`**], α_V_ contains canonical fields such as ID and Version for *Ensembl* nodes, and α_E_ contains either history metadata (**`old_release, new_release, weight`**) or cross-reference metadata (connection and derived available releases). This explicit typing is a methodological choice: it ensures that downstream algorithms can enforce invariants such as “history travel only within a node type” and “xref validity is release-scoped” by local checks rather than by global, fragile assumptions.

#### Node taxonomy (types, naming, and normalization)

Node types are enumerated in DB as form-specific *Ensembl* labels (**DB.nts_*Ensembl***), assembly-specific *Ensembl* labels (**`DB.nts_assembly`**), base gene labels without version suffixes (**DB.nts_base_*Ensembl***), and external labels (**`DB.nts_external; idtrack/idtrack/_db.py`**). Versioned *Ensembl* nodes are labeled <**`stable_id`**>.<version> using **`DB.id_ver_delimiter`** (“.”), and the graph stores both ID and Version attributes to make version stripping and base-ID reasoning explicit and reversible (**`idtrack/idtrack/_database_manager.py:node_n ame_maker`**). External nodes are labeled by the raw external identifier string id db and may include synonym-prefixed nodes such as **`synonym_id::<label>`** that originate from *Ensembl* **`external_synonym`** materialization (Section 6.3), which prevents conflating free-text labels with accession-like identifiers while still allowing synonyms to participate as explicit, allowlisted bridge nodes.

#### Edge taxonomy (history versus cross-reference edges)

History edges E_H_ encode the *Ensembl* time axis and are derived from **`stable_id_event`** joined to **`mapping_session`** via **`DatabaseManager.create_id_history`**, producing directed edges between *Ensembl* identifiers with attributes **`old_release, new_release`**, and **`weight`** (*Ensembl* score with zeros treated as missing), and optionally mapping-session metadata when built with **`narrow=False (idtrack/idtrack/_graph_maker.py: construct_graph_form`**). To make the time axis total for traversal, **`GraphMaker.construct_graph_form`** also adds explicit birth edges from a sentinel “Void” version and retirement edges to a sentinel “Retired” version, and it adds an ∞ self-loop for latest-release entries, which turns “still active” into an explicit edge case rather than a missing-edge convention (**`idtrack/idtrack/_db.py and idtrack/idtrack/_graph_maker.py`**). Cross-reference edges E_X_ encode non-history relations (external → *Ensembl*, transcript → gene, translation → transcript, base → versioned) and carry a structured payload under **`DB.connection_dict`** with canonical shape **connection_e_ : name db ↦ assembly ↦** 2^ℛ^, i.e. **`{db_name: {assembly: set(releases)}}`**, which records provenance and temporal validity as an explicit set-valued object **`(idtrack/idtrack/_graph_maker.py:construct_graph`**). Unlike history edges, cross-reference edges are intentionally collapsed to one multi-edge per ordered node pair (edge key 0), and repeated evidence across releases or databases is accumulated by set union inside **`connection`**, which avoids edge explosion while ensuring the graph remains a provenance-preserving ledger rather than a lossy lookup table.

#### Release validity on cross-reference edges (derived **available releases**)

After construction, every cross-reference edge is annotated with an **`available_releases`** set computed as ⋃_d,a_ connection_e_ [d][a], because synonym expansion must enforce that a hop is valid at a declared release rather than treating a cross-reference as timeless. This derived attribute is used directly by synonym-search routines (Section 6.7) as a constant-time membership test **`from_release in available_releases`**, which is a deliberate design choice that moves repeated temporal validity checks from adjacency scans into cached set membership.

**Algorithm. S2.**
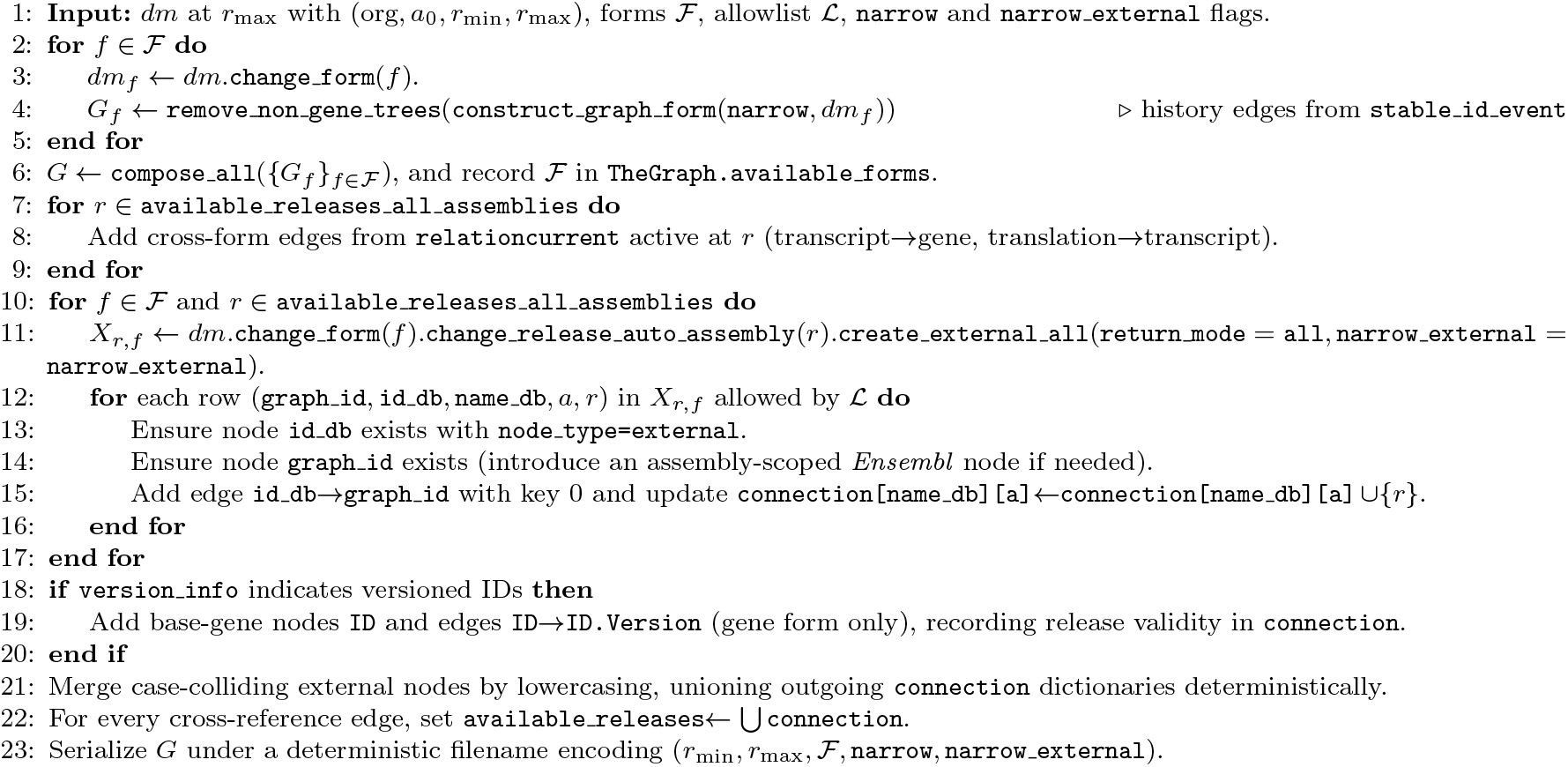
Snapshot graph construction as a deterministic pipeline from cached *Ensembl* tables and an explicit external allowlist, with edge payloads designed to preserve provenance and release validity as machine-checkable objects.

#### Graph construction algorithm (GraphMaker; full-width pseudo-spec)

##### Case-collision handling (external-node merging)

Real identifier lists frequently contain external accessions that differ only by capitalization, so **`GraphMaker`** merges external nodes that become identical under **`lower()`** and unions their outgoing edge metadata, selecting a canonical representative name by a deterministic ordering over degree, reverse-degree, and uppercase count (**`idtrack/idtrack/_graph_maker.py: _merge_nodes_with_the_same_in_lower_case`**). Because this merge is performed before **`TheGraph.lower_chars_graph`** is materialized, subsequent case-insensitive resolution becomes a bijection (lowercase_id ↦ canonical_id) rather than a heuristic, which eliminates nondeterministic casing behavior at query time and therefore strengthens reproducibility under heterogeneous input formatting.

#### Complexity (build-time)

Building history edges is O(|**`stable_id_event ⋈ mapping_session`**| +Σr_∈ ℛ_ |**`ids`**_r_|) for the chosen form(s) plus I/O proportional to serialized tables, while adding cross-reference edges is O(Σ_r,f_ |X_r,f_ |) with constant-factor overhead for dictionary unions inside connection. The final graph occupies O(N + M_H_ + M_X_) memory, and the case-collision merge runs in O(N_external_ + M_X,external_) time because it groups nodes by lowercase key and unions outgoing edge metadata, which is linear in the number of affected external nodes and their outgoing edges.

### 6.6 Query optimization and cached summaries (why queries are tractable)

IDTrack’s query-time performance is not achieved by approximating semantics but by constructing deterministic derived indexes as cached graph properties, so traversal routines can consult compact dictionaries instead of repeatedly iterating raw NetworkX adjacency structures and recomputing release/assembly validity. These caches are implemented as **@cached property** attributes on **`TheGraph (idtrack/idtrack/ _the_graph.py`**), meaning each is computed once on first access and then stored as a normal attribute for the life of the graph instance, with an optional eager warm-up via **`API.calculate_graph_caches`** to stabilize latency across batch runs (**`idtrack/idtrack/_api.py`**).

#### Cache contract (derived relations as pure functions of the snapshot)

Conceptually, each cached property is a deterministic function Ψ (G) that preserves semantics while changing representation, and the contract posture is that Ψ must depend only on the serialized snapshot graph and not on external services, clocks, or randomized orderings, so cached summaries remain reproducible evidence rather than hidden state. This is why caches are stored on the graph instance and can be re-materialized from the pickled snapshot, instead of being recomputed ad hoc per query where subtle differences in iteration order or endpoint availability could create hard-to-audit divergences.

#### Case-insensitive resolution cache (**lower_chars_graph**)

**`TheGraph.lower_chars_graph`** constructs a bijection **`lowercase_id_↦ canonical_id`** by iterating over all nodes once and raising an error on collisions, which turns case-insensitive resolution into a deterministic precondition of the graph rather than a query-time heuristic (**`idtrack/idtrack/_the_graph.py: lower_chars_graph`**). This engineering detail is scientifically relevant because it prevents ambiguous casing from becoming a hidden degree of freedom in downstream joins and conversions, and it makes failures explicit at build/load time where they can be audited as substrate problems rather than misattributed to individual query inputs.

Formalization (why these summaries are the right indexes). For each node v, the cross-reference payload induces a relation Г(v) = {(d, a, r) : ∃e = (v → u, 0) ∈ E_X_ ^ r ∈ connection_e_[d][a]}, and **`combined_edges`** is the dictionary representation of Г(v) projected into the nested map d ↦ a ↦ {r}, while **`available_releases`** is the marginal {r} used as a constant-time validity predicate. Similarly, **`node_trios`** is the per-node expansion of Г(v) into explicit triples (**`database, assembly, release`**), which is computationally heavier but turns provenance inference and reporting into a frequency-count problem rather than an implicit guess.

#### Reverse view (zero-copy temporal inversion)

**`TheGraph.rev`** is a non-copying reversed view (**`reverse(copy=False`**)) that shares underlying data structures while inverting edge direction, allowing backward-in-time traversal without duplicating 𝒢 in memory (**`TheGraph.rev; idtrack/idtrack/_the _graph.py`**). This is a small but important engineering choice: reverse-time conversions become a change in view rather than a change in representation, so both directions share the same cached edge metadata and remain consistent under serialization.

#### Combined edge dictionaries (fast release-scoped synonym lookups)

**`TheGraph.combined_edges`** aggregates cross-reference edge metadata into nested dictionaries implementing **`db_name -> assembly -> set(releases`**) per source node, and **`combined_e dges_genes`** and **`combined_edges_assembly_specific_genes`** provide analogous gene-scoped variants derived by traversing the reversed graph (**`idtrack/idtrack/_the_graph.py`**). These structures are not merely convenient: synonym search and source inference use them to test release validity by set membership (**`from_release in available_releases`**) rather than by scanning edge lists, which shifts repeated checks from O(deg(v)) per hop to O(1) average per hop at query time.

Activity ranges and origin trios (time and coordinate inference). **`TheGraph.get_active_ranges_of_id_caches`** inclusive active release intervals for every node, while **`TheGraph.node_trios`** caches, for each node, the set of origin triples (database, assembly, release) in which that node is active, both of which are derived deterministically from combined edge metadata and used by temporal orientation and source inference (**`idtrack/idtrack/ _the_graph.py`**). Computationally, materializing these caches is O(N + M_X_) in the sense that it requires iterating nodes and consulting their combined-edge metadata, but this cost is paid once per loaded snapshot and then amortized across all conversions and QC routines, which is a deliberate trade-off given that atlas-scale workflows often involve millions of identifier queries against a fixed snapshot.

#### Hyperconnective hubs (bounded branching)

**`TheGraph.hyperconnective_nodes`** caches the set of external nodes whose out-degree exceeds **`DB.hyperconnecting_threshold`** (default 20; **`idtrack/idtrack/_db.py`**) and is used as a hard pruning rule in synonym expansion, because these nodes explode the frontier and frequently represent coarse-grained accessions with weak semantics (Section 6.8). Computing this cache is O(N) by scanning node degrees once, and applying it turns potentially unbounded synonym branching into a controlled, auditable policy decision that can be reported as part of the mapping contract.

### 6.7 Pathfinder conversion semantics (formal pseudo-spec)

A conversion request in IDTrack is a constrained search problem parameterized by the mapping contract. Given a raw identifier string, an optional origin release (**`from_release`**), a target release (**`to_release;`** defaulting to **g.graph[‘*Ensembl* release’]**), an optional output namespace (**`final_database`**), a selection policy (**`strategy=‘best’|’all’`**), and an audit flag (**`explain`**), the software resolves the query into the snapshot graph and enumerates admissible time-respecting paths. It can optionally perform bounded external bridging, scores candidates deterministically, and returns explicit outcome semantics. This framing matters scientifically because it makes every boundary explicit: “a mapping” is not a timeless lookup but a reachable target under (r_min_, r_max_, assembly, allowlist, search bounds, scoring order), and the output explicitly reports when no mapping exists, when a unique mapping exists, and when multiple plausible mappings exist under the declared contract.

#### Formal optimization view (admissible paths and deterministic objective)

Let q be the raw query string and let v_0_ = c(q) be its canonical node after normalization (Step 0), and let r_to_ be the target release coordinate, so the conversion problem is to identify the set of candidate targets T (q) ⊆ V that are reachable under the declared contract and active at r_to_. We denote the set of admissible paths from v_0_ to a target t under contract parameters by П(v_0_, t; 𝒞), where admissibility enforces (i) time-respecting history traversal, (ii) release-valid xref steps, (iii) bounded external jumps and synonym depth, and (iv) explicit pruning of hyperconnective hubs, and we emphasize that П is empty when the contract boundary excludes a mapping even if some mapping exists under a different boundary. For each admissible path p ∈ П, IDTrack computes a lexicographically ordered score vector.

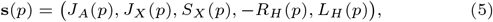

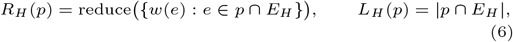

Here J_A_ is the assembly-jump penalty computed by **`Track.minimum_assembly_jumps`**, J_X_ and S_X_ are external jump/step tallies, w(e) are *Ensembl* history weights aggregated under the declared **`reduction`** and **`remove_na`** policy, and the sign convention matches code by minimizing **`edge_scores_reduced`** = − R_H_ (p) (**`idtrack/idtrack/_track.py:calculate_score_ and_select`**). Under strategy=‘all’, the method returns the per-target argmin set 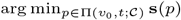 for all surviving targets, while under **`strategy=‘best’`** it applies an additional global selector over targets that incorporates final-conversion confidence and assembly priorities, making “best” a declared policy rather than an unreported coercion (**`idtrack/idtrack/_t rack.py:_path_score_sorter_all_targets`**).

#### Step 0: normalize the query into a canonical graph node

The raw string is resolved to a canonical node label via **`TheGraph.node_name_alternatives`**, which implements a deterministic cascade (exact match in g.nodes, case-insensitive match via **`lower_chars_graph`**, version-suffix trimming and delimiter normalization, dash/underscore substitutions, and a single retry under the **`DB.synonym_id_nodes_prefix`** prefix) so common formatting drift does not become silent mapping failure (**`idtrack/idtrack/_the_graph.py`**). Because **`lower_chars_graph`** is a bijection after case-colliding external nodes are merged during graph construction, this normalization step is not a heuristic but a deterministic resolution procedure whose failure (**`graph_ id=None`**) is itself a reportable outcome class.

**Algorithm. S3.**
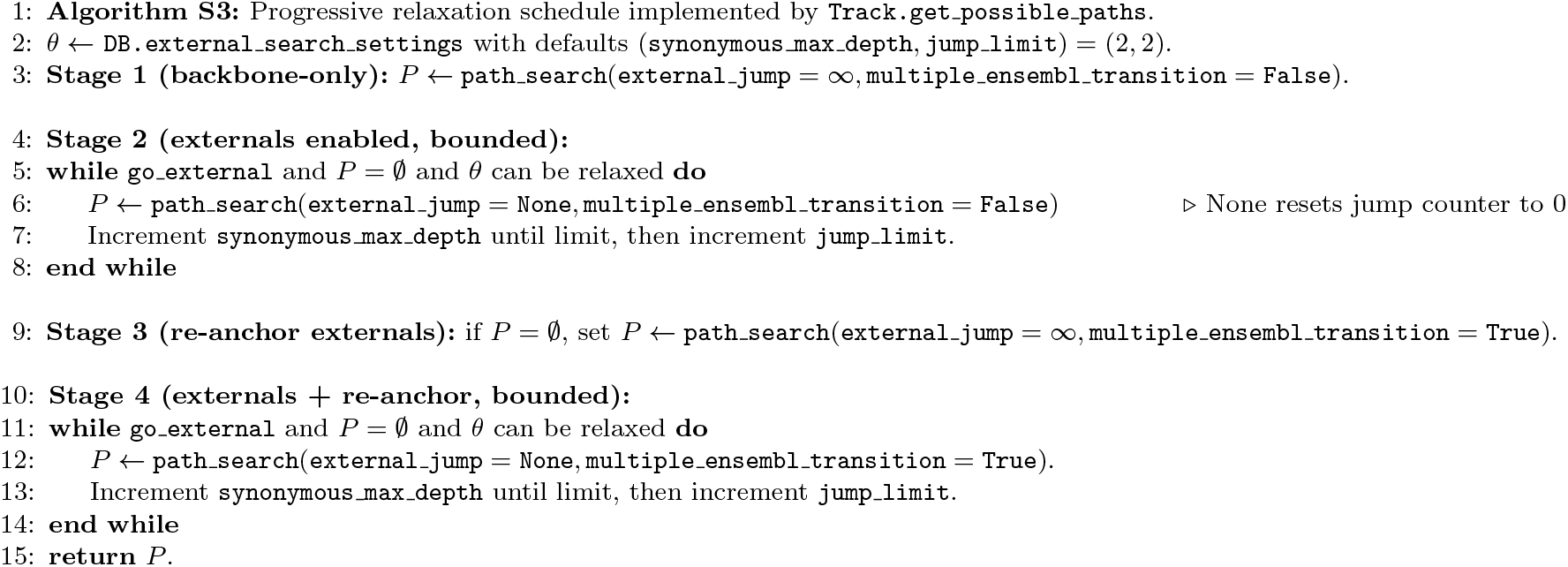
IDTrack searches for an admissible explanation under progressively relaxed bounds, beginning with backbone-only time travel and enabling bounded external bridging only when necessary, so returned paths remain interpretable under a declared contract.

#### Step 1: determine temporal orientation when the origin is unknown

If **`from_release`** is not provided, **`Track.should_graph_reversed`** infers whether a forward search (old → new), reverse search (new → old), or a split “both” search is required, based on the active release intervals in **`TheGraph.get_active_ranges_of_id`** and by selecting start releases closest to **`to_release (Track.get_from release_and_reverse_vars; idtrack/idtrack/_t rack.py`**). This explicit orientation step is a reproducibility safeguard: it forces the method to declare which time direction is being traversed rather than implicitly assuming “latest → target” or “target → latest”, which would collapse temporal semantics into a hidden default.

#### Step 2: enumerate admissible paths with progressive relaxation

Candidate paths are generated by **`Track.get_possible_paths`**, which runs **`Track.path_search`** under a staged relaxation schedule that is intentionally biased toward interpretable explanations. It first disables external jumps entirely (external jump=np.inf), then enables bounded external bridging under **`DB.external_search_settings`** and incrementally increases synonym depth and jump limits until a path exists or configured maxima are reached. Finally, it enables a “multiple-*Ensembl* transition” fallback for cases where an external starting node cannot be anchored to the inferred **`from_release (idtrack/id_track/_track.py:get_possible_paths`**). Default bounds are explicitly stored in **`DB.external_search_settings`** (**`jump_limit=2, synonymous_max_depth=2; idtrack/idtrack/_db.py`**), and any deviation (including relaxation bounds) changes the definition of “admissible path” and should therefore be reported as part of the mapping contract. Algorithm S3 (Figure 3) provides a minimal pseudo-spec of this relaxation schedule, so the admissibility boundary is not inferred from prose but is stated as a reproducible stage ordering with explicit knobs.

#### End-to-end conversion orchestration (Track.convert; full-width pseudo-spec)

Algorithm S4 (Figure 4) states the end-to-end orchestration contract as a code-faithful pseudo-spec, so normalization, temporal orientation, staged path enumeration, scoring, and final-namespace conversion are exposed as an explicit decision tree rather than as an implicit side effect of the API call.

#### Step 3: path search engine (history DFS with bounded synonym “beam-up”)

**`Track.path_search`** performs a depth-first traversal over history edges (edges between nodes of the same *Ensembl* node type), stepping only along chronologically admissible transitions returned by **`Track.get_next_edges`** and terminating when the target release is reached in the appropriate direction, which in the absence of synonym bridging is an O(M_H_ + N) traversal over the reachable backbone subgraph in the standard DFS sense (worst-case O(|V |+|E|) for explicit graphs), while recognizing that returning all admissible paths can be exponential in the number of branch points. When the traversal is stranded (e.g., starting from an external node or encountering a non-backbone segment), the algorithm can “beam up” through external identifiers via **`choose_relevant_synonym`**, subject to jump and depth bounds, and it suppresses recurrence on non-backbone segments by tracking a set of visited non-backbone nodes so that synonym search does not cycle through repeated external hubs (**`idtrack/idtrack/ track.py:path_search`**). Algorithm S5 (Figure 5) gives a code-faithful pseudo-spec of this DFS, including the precise placement of external “beam-up” segments as explicit xref edge records with release-context payloads.

#### Synonym discovery (synonymous nodes and depth semantics)

Synonym discovery is implemented by Track.synonymous nodes, which delegates to a recursive search recursive synonymous that explores cross-reference edges while forbidding history travel. It never traverses an edge whose source and target share the same node type, and it enforces a depth budget defined as the maximum count of any single node type along the path. This implementation-level choice bounds repeated excursions into the same node-type strata (notably external) even when edge counts are large (idtrack/idtrack/ track.py: recursive synonymous). Release validity is enforced during this search by testing whether from release belongs to available releases on the traversed edge (or to the combined-edge summary when terminating on an external node), and bounce patterns (e.g., A → B → A with the same underlying *Ensembl* ID but different Version) are suppressed to avoid spurious oscillations between closely related *Ensembl* forms that do not contribute new mapping evidence (**`idtrack/idtrack/_track.py`**). Algorithm S6 (Figure 6) specifies **`Track._recursive_synonymous`** as executed, including the depth semantics, directionality rules for shallow backbone search, hyperconnective pruning, and release-valid edge filtering.

**Algorithm. S4.**
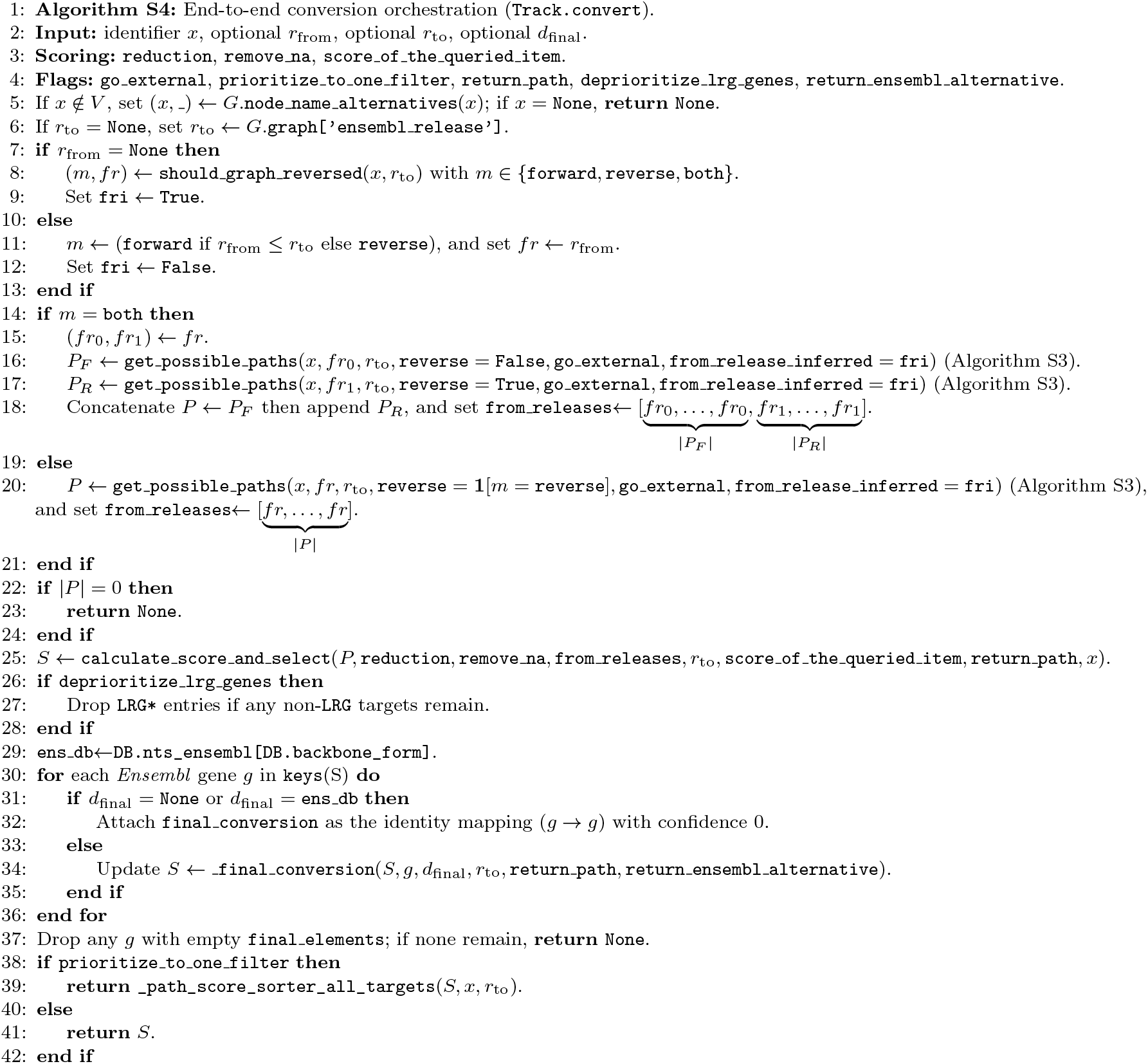
Track.convert is the orchestration layer that normalizes identifiers, selects a temporal orientation, enumerates admissible paths under progressive relaxation (Algorithm S3), scores deterministically, and only then performs final-namespace conversion and global tie-breaking under a declared policy.

**Algorithm. S5.**
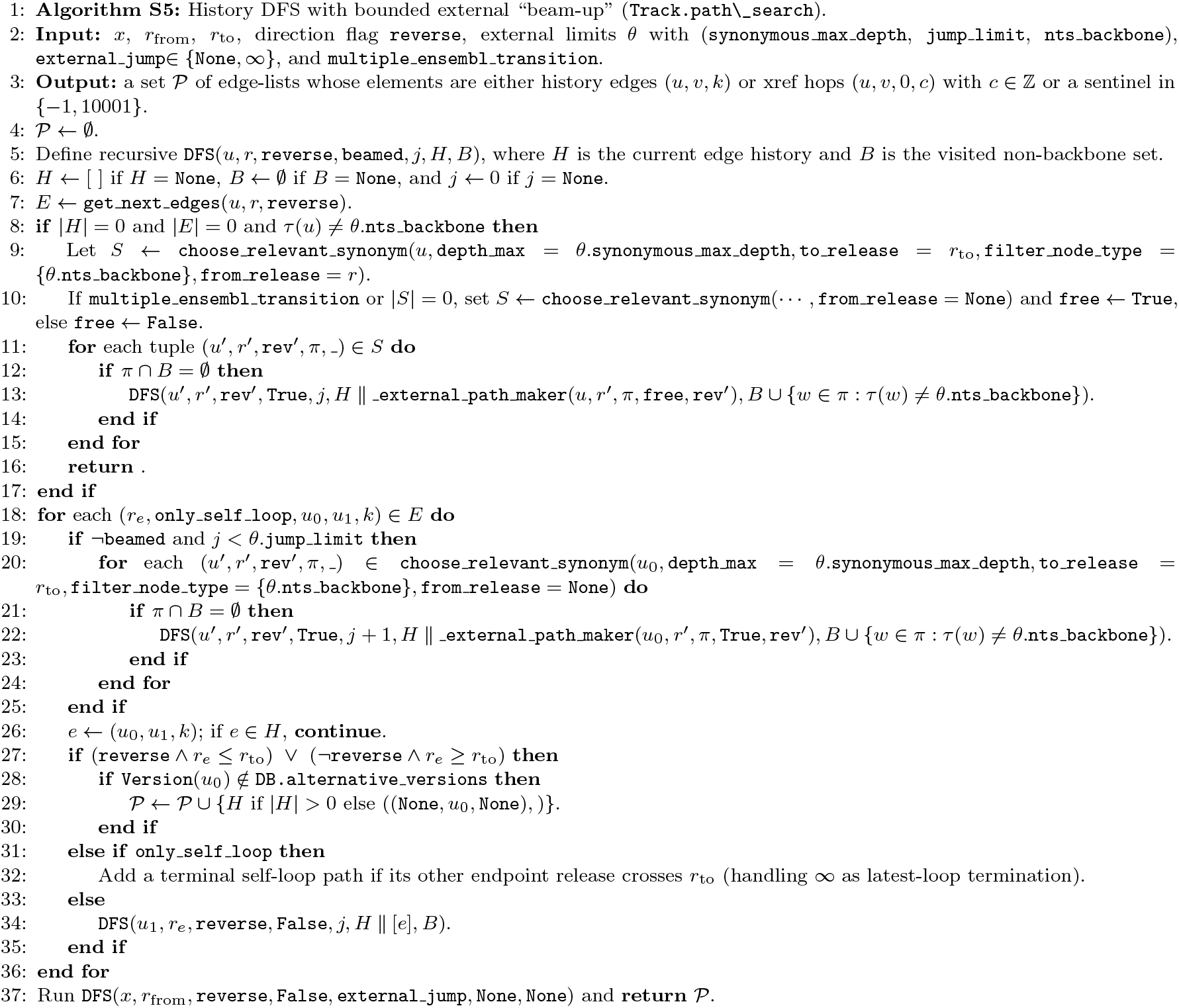
Track.path search enumerates admissible time-respecting history paths and inserts bounded external “beam-up” segments as explicit xref edge sequences whose release context is recorded as either an integer release or a sentinel placeholder for inferred entrances.

#### Scoring and deterministic selection (best per target, and optional best overall)

Once candidate paths are found, **`Track.calculate_score_and_select`** computes per-path metrics—**`external_step, external_jump, assembly_jump`**, **`edge_scores_reduced`** (negative reduced history evidence under the declared reduction and **`remove_na`** policy), and ***Ensembl*e_step** —and stores them as a destination-keyed score dictionary (**`idtrack/idtrack/_track.py:calculate_score_and_select`**). For each destination *Ensembl* gene, **`_path_score_sorter_s ingle_target`** selects the lexicographically smallest score vector in the fixed importance order **assembly_jump, external_jump, external_step, edge_scores_reduced, *Ensembl*e_step**, which makes tie-breaking explicit and keeps “best” contract-defined rather than implicit (**`idtrack/idtrack/_track.py:_path_score_sort er_single_target`**). If the API is called with **`strategy=‘best’`** (implemented by setting **`prioritize_to_one_filter=True`** in **`Track.convert`** via **`API.convert_identifier`**), the method then applies a global selector across all surviving target pairs. This selector incorporates **`final_conversion_confidence`** and assembly-priority summaries, and it breaks ties by preferring “same as input” targets and, when needed, by a two-hop synonym richness heuristic **`calculate_node_scores`**, thereby making “best” a declared policy with explicit ordering rather than an implicit coercion (**`idtrack/idtrack/_track.py:_path_score_sorter _all_targets`**). Algorithm S7 (Figure 7) provides a compact pseudo-spec of this scoring loop and its per-target lexicographic selector, making the ranking objective explicit and therefore reconstructible.

#### Final conversion into an external namespace (confidence and fallbacks)

If **`final_database`** is requested, **`Track._final_conversion`** performs a final synonym search from the resolved *Ensembl* gene to the requested external database. It first restricts to synonyms active at the target release (confidence flag 0) and then, if necessary, falls back to a release-agnostic search (confidence flag 1) that annotates each hop with the set of releases in which it is valid, because external availability is itself release- and assembly-scoped and should not be silently assumed (**`idtrack/idtrack/_track.py:_final_conversion`**). If no external synonym exists and **return_*Ensembl*_alternative=True** (default), the method returns the *Ensembl* gene as an explicit fallback with **`final_conversion_confidence=np.inf`**, which is surfaced at the API layer as **`no_target=True`** so downstream code can distinguish “no path under the contract” from “path exists but the requested namespace has no admissible label” (**`idtrack/idtrack/_api.py:convert_identifier`**).

#### Result schema (explicit outcome semantics)

**`API.convert_identifier returns`** a machine-readable dictionary whose fields encode outcome semantics rather than requiring the caller to infer them from missing values: **`no_corresponding`** signals that the query could not be normalized to any graph node, **`no_conversion`** signals that the query exists but no admissible path exists under the contract, and **`no_target`** signals that time travel succeeded but the requested external namespace lacked a synonym, while **`target_id, last_node`**, and **`final_database`** report what was returned and under which namespace label (**`idtrack/idtrack/ api.py:convert_identifier`**). When **`explain=True`**, the API attaches **`the_path`**, a structured mapping from (**target_id, *Ensembl*e_gene_id**) to an ordered tuple of edge records that concatenates the time-travel path and the final conversion hop. History edges are stored as (**`u,v,key`**) tuples and cross-reference hops as (**`u,v,0,context`**) tuples, where the final element records either an explicit release, an available-release set, or a placeholder sentinel for inferred entrance releases. This makes the audit object mechanically verifiable against the cached snapshot rather than narrative.

**Algorithm. S6.**
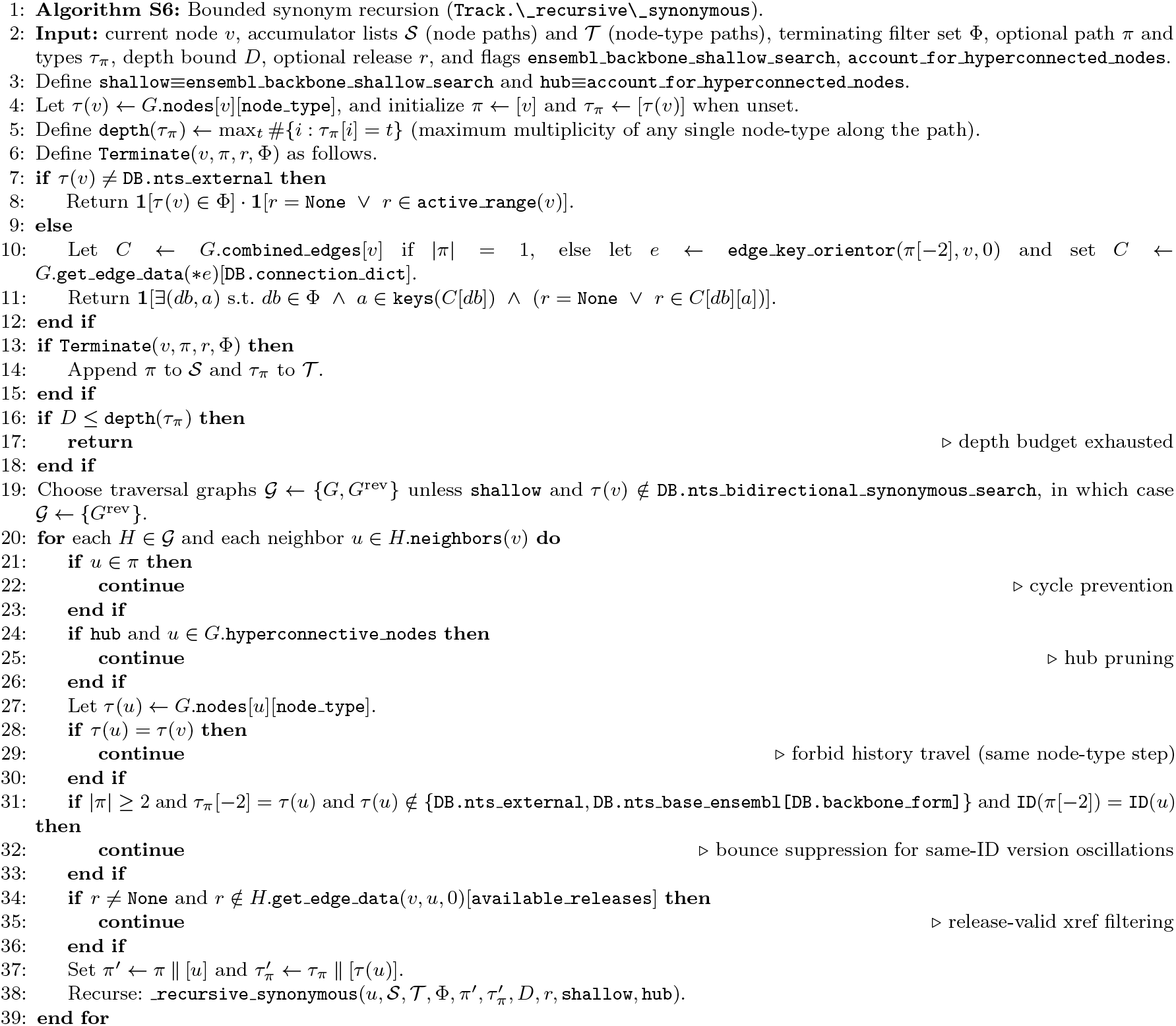
**`Track._recursive_synonymous`** enumerates release-valid synonym paths under a depth metric defined on node-type multiplicities, while forbidding same-type steps (no history travel), pruning hyperconnective hubs, suppressing same-ID bounce patterns, and supporting a shallow mode that restricts directionality except for explicitly bidirectional node-type classes.

#### Complexity (query-time, with explicit bounds)

For a fixed snapshot graph, node normalization is O(1) average in the common case because it performs a bounded number of dictionary lookups over **`g.nodes`** and **`lower_chars_graph`**. The cost of the path search is dominated by the number of admissible history branches and bounded synonym expansions, so worst-case path enumeration is exponential in the number of branch points even when each local step is O(1) average. In practice this explosion is constrained by the finite release window size R, by explicit bounds **`jump_limit`** and **`synonymous_max_depth`**, by release validity checks via **`available_releases`**, and by hyperconnective-hub filtering. Scoring is linear in the total length of enumerated paths because it reduces per-edge weights and computes assembly penalties by per-step dictionary and set operations, so once admissible paths are bounded the remainder of the conversion pipeline is deterministic and computationally transparent.

### 6.8. Hyperconnective nodes (tractability and semantic safeguards)

External identifiers can be hyperconnective (high out-degree) due to low-specificity naming or aggregation conventions, and such nodes are dangerous in two ways: computationally they explode synonym-search branching, and semantically they can masquerade as evidence by connecting unrelated entities through promiscuous hubs. IDTrack therefore detects hyperconnective nodes as a cached graph property by scanning all nodes once, selecting those with **`node_type==external`** and **`out_degree > DB.hyperconnecting_threshold`** (default 20), and caching the resulting dictionary **`{node: out_degree}`** for subsequent constant-time membership tests during traversal (**`TheGraph.hyperconnective_nodes; idtrack/idtrack/ _the_graph.py`** and **`idtrack/idtrack/ _db.py`**).

**Algorithm. S7.**
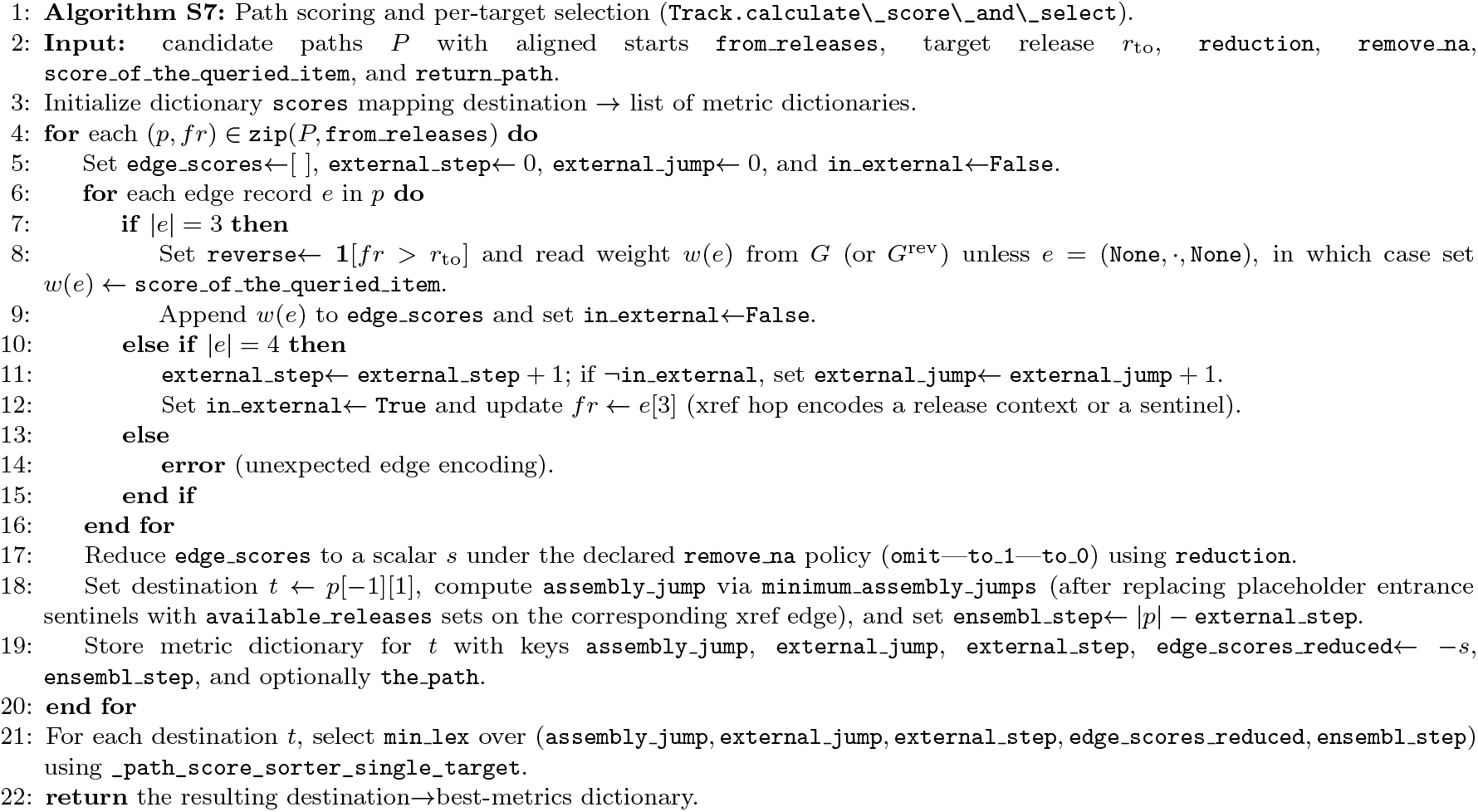
Scoring is a deterministic collapse of enumerated paths into one best explanation per target under a lexicographic objective, with explicit accounting for external steps/jumps, assembly transitions, and reduced history evidence.

#### Formal definition (degree-threshold pruning as an explicit bound)

Let Δ^+^(v) denote the out-degree of node *v* in *G*, and define the hyperconnective set *H* = {v ∈ V : τ (v) = external ∧ Δ^+^(v) > τ_hub_} with τ_hub_ = **`DB.hyperconnecting_threshold`**, so hub filtering is a declared predicate applied during synonym expansion rather than an implicit performance hack. Because synonym discovery is the main source of combinatorial growth in the search frontier, explicitly bounding H converts an otherwise unbounded local branching factor into a contract parameter that can be reported and audited, and it makes failure modes interpretable as “pruned by declared policy” rather than as opaque timeouts or silent truncation.

#### Complexity and interpretability impact

Computing the cache is *O(N)* (degree lookups over nodes), and applying it prunes neighbor expansions in synonym search without altering backbone history structure, which is precisely the trade-off we want under a contract: constrain search to paths that are more likely to be semantically defensible while keeping the pruning rule explicit, deterministic, and reportable. In practice, the precision penalty is typically negligible because hyperconnective nodes tend to correspond to coarse-grained identifiers with weak one-to-one semantics, while meaningful synonym relations are usually reachable through alternative, more specific external accessions, so filtering improves both tractability and the interpretability of returned explanations.

### 6.9 Explainability payload (path objects and auditability)

Explainability is exposed as an opt-in payload that returns the concrete traversal path taken through the graph for each surviving candidate target, enabling users to audit conversions, debug unexpected ambiguity, and generate provenance records that are more informative than a single target string. At the API layer, **`API.convert_identifier(…, explain=True`**) constructs **`the_path`** as a mapping from (**target_id, *Ensembl*e_gene_id**) to a concatenated edge list that includes both the time-travel segment (**`Track.convert(…, return_path=True`**)) and the final conversion hop (**`final_conversion[‘final_elements’][target][‘the_path’]; idtrack/idtrack/ _api.py`** and **`idtrack/idtrack/ _track.py`**).

#### Formal edge encoding (why this is machine-checkable)

History edges are stored as (**`u,v,key`**) triples that correspond to concrete MultiDiGraph edges carrying **`old_release`**, **`new_release`**, and **`weight`**. Cross-reference edges are stored as (**`u,v,0,context`**) tuples whose final element records the release context used to justify that hop (either an integer release, a set of releases in which the edge is valid, or a sentinel placeholder for inferred entrance releases), making the path a structured object rather than a free-text narrative. Concretely, inferred entrance releases are encoded as sentinel integers **`-1`** (forward search) and 10001 (reverse search) via **`Track. _external_entrance_placeholder`**, and because these lie outside any plausible *Ensembl* release window they make the audit payload unambiguous about which hops used inferred temporal anchoring (**`idtrack/idtrack/ _track.py`**). Because these edge records are direct projections of graph attributes (and, for cross-reference edges, of **`available releases`**), a reviewer or downstream QC script can validate a returned explanation mechanically against the cached snapshot, which is the intended outcome when “explainability” is treated as reproducible evidence rather than as interpretive commentary.

#### Mechanical verification recipe (turning explanations into checkable evidence)

Given **`the_path`**, validation reduces to deterministic lookups: for each history edge (**`u,v,k`**) one verifies that the graph contains that multiedge and that its **`old_release/new_release`** metadata are chronologically compatible with the declared traversal direction. For each xref edge (**`u,v,0,context`**) one verifies that **`available_releases`** contains the integer **`context`** or intersects the set-valued **`context`**, which is precisely why IDTrack stores release validity as explicit sets rather than as implicit assumptions. When a path includes inferred entrance steps, the placeholder is replaced during scoring by the edge’s **`available_releases`** set for assembly-jump computation, so the audit object retains an explicit, checkable representation of uncertainty rather than hiding it in narrative (see **`er_maker_for_initial_conversion`** in **`idtrack/idtrack/_track.py:calculate_score_and_select`**).

#### Operational trade-offs

Explainability can increase output size, especially under ambiguity when **`strategy=‘all’`** returns many candidate targets, so it is exposed as an opt-in payload that allows high-throughput pipelines to keep compact outcome summaries by default while enabling targeted audits when a conversion becomes scientifically consequential. Under the contract framing, this is the correct trade-off: explainability is the mechanism that turns a conversion into an auditable argument, but it should be paid for only when the analysis stage demands that level of evidence density.

### 6.10. Validation and quality control (graph invariants and test harness)

Reproducibility depends not only on documenting intended behavior but also on detecting when upstream irregularities (annotation mistakes, duplicated history events, schema anomalies) would cause the constructed snapshot to violate the invariants assumed by the pathfinder, so IDTrack includes both internal build-time consistency checks and an explicit developer-facing test harness. Build-time checks are enforced by raising errors when invariants are violated (e.g., unexpected delimiter usage inside stable IDs, duplicated **`mapping_session`** rows per **`new_release`**, or inconsistent edge multiplicity for cross-reference edges that must be unique per ordered pair), which is an explicit design decision to fail loudly rather than to silently absorb corrupted history into a misleading mapping substrate (**`idtrack/idtrack/ _database_manager.py`** and **`idtrack/idtrack/ _graph_maker.py`**).

#### TrackTests (white-box invariant checks)

**`TrackTests (idtrack/idtrack/ _track_tests.py`**) is a subclass of **`Track`** that adds read-only invariant checks intended for developers and advanced users, and it can be instantiated via **`API.build_graph(…, return_test=True`**) when a study wants to validate a built snapshot before using it in large-scale conversions. Examples of enforced invariants include consistency between SQL-derived ID lists and graph-derived ID lists across releases, agreement between cached activity ranges and raw combined-edge metadata, and non-overlap of sibling active ranges for base *Ensembl* IDs, because these conditions are what make the time axis coherent and ensure that pathfinding semantics (e.g., “closest active boundary”) are well-defined.

#### Representative invariants (expressed as checkable equations)

Let 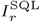 be the set of *Ensembl* IDs returned by **`DatabaseManager.get_db(‘ids’)`** at release r for the primary assembly, and let 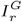 be the set returned by the graph’s release-scoped ID listing utilities. The core integrity requirement is 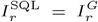 for all r ∈ **`g.graph[‘confident_for_release’]`**, because any discrepancy means the snapshot is not a faithful materialization of the declared coordinate system (**`idtrack/ idtrack/_track_tests.py:is_id_functions_consistent_ens embl`**). Let *b* be a base-gene node (stable ID without version suffix) and let *{v_i_}* be its versioned children. The time-axis coherence requirement is that the active sets are disjoint, i.e. 𝒜 (v_i_) ∩ 𝒜 (v_j_) = ∅ for *i ≠ j*, because overlapping sibling activity would make “which version was active at release r” ill-defined and would break deterministic orientation and scoring (**`idtrack/idtrack/_track_tests.py:is_range_functions_robust`**). Similarly, cache consistency is asserted by requiring that active ranges computed from raw cross-edge release sets equal the cached **`get_active_ranges_of_id`** output for all relevant nodes, which is what makes orientation inference and provenance inference logically grounded rather than heuristic (**idtrack/idtrack/_track_tests.py:is_id_functions_consistent_*Ensembl*_2**).

#### QC as a reportable workflow step

In practice, we recommend treating QC as a build-stage step that is itself part of the mapping contract, because a snapshot graph that violates invariants is not merely a performance issue but a semantic failure that can invalidate downstream mapping claims. This posture also strengthens credibility in the engineering sense: exposing invariants and tests signals that the mapping layer is engineered as a scientific instrument whose internal consistency can be checked, rather than as a black-box conversion utility that implicitly assumes upstream correctness.

### 6.11. Batch conversion and outcome classification

Batch conversion is implemented as a thin wrapper over single-identifier conversion that adds progress reporting and yields per-identifier result dictionaries without changing semantics: **`API.convert_identifier_multiple`** simply iterates **`API.convert_identifier`** over an input list and preserves the same contract-facing fields (**`target_id, final_database, no_corresponding, no_conversion, no_target`**, and optional **`the_path; idtrack/idtrack/ _api.py`**). This design is essential for pipeline integration because it makes conversions first-class records rather than strings, enabling downstream code to aggregate outcomes into reproducible summaries without re-implementing conversion semantics.

#### Outcome classification (overlapping diagnostic bins)

**`API.classify_multiple_conversion`** bins batch results into overlapping semantic categories that separate mapping failure modes and ambiguity classes, notably **`matching_1_to_0`** (either **`no_corresponding`** or **`no_conversion`**), **`matching_1_to_1`**, **`matching_1_to_n`**, **`changed_only_1_to_1`**, **`changed_only_1_to_n`**, and the **`alternative_target_*`** categories that flag *Ensembl* fallbacks when **`no_target=True`** (**`idtrack/idtrack/ _api.py`**). **`API.print_binned_conversion`** then reports counts and percentages for these bins, which is the operational bridge between low-level conversion records and manuscript-facing summaries such as Figure 7, because it converts per-identifier outputs into interpretable outcome profiles without collapsing ambiguity into a single number.

#### Complexity

Batch conversion is *O(K)* calls to the single-item converter for *K* inputs plus post-processing time linear in *K* for classification, so the method scales transparently with input size and does not introduce hidden behavior beyond the per-identifier pathfinder cost described in Section 6.7. This transparency is deliberate: batch wrappers should not change mapping semantics, because doing so would make large-scale pipeline behavior less auditable than interactive behavior.

**Algorithm. S8.**
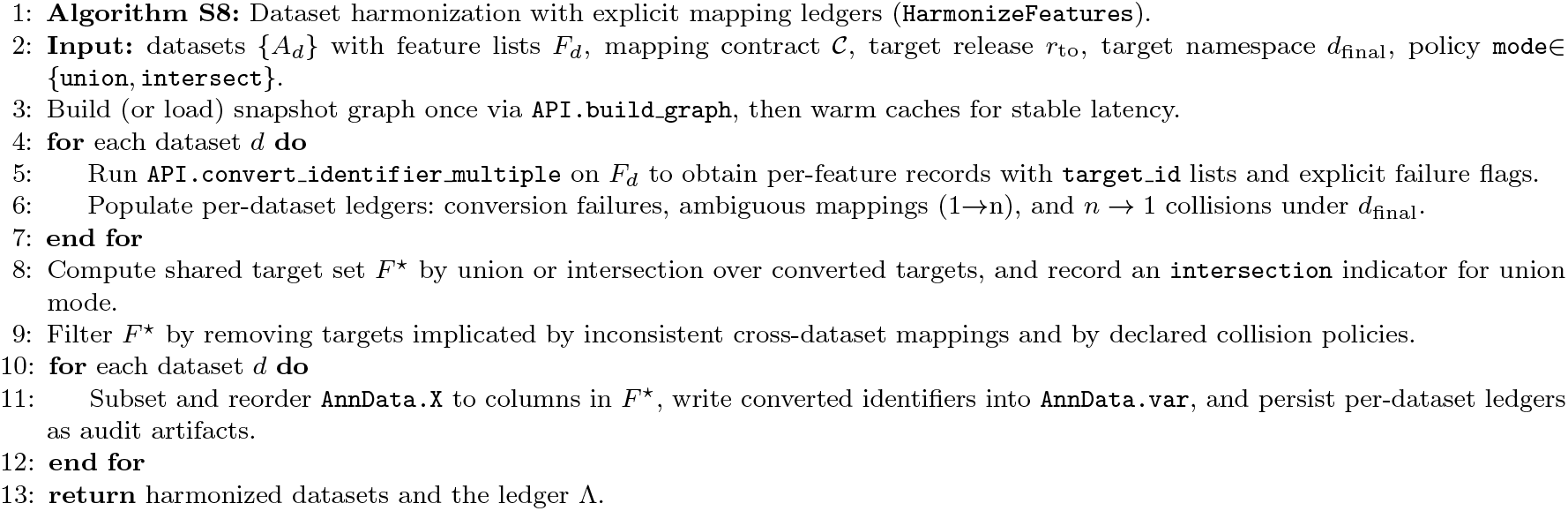
Algorithm S8: Harmonization is treated as a reproducible join over explicit IDTrack outcomes, with ledgers that preserve loss and ambiguity semantics instead of collapsing them into silent preprocessing.

### 6.12. HarmonizeFeatures internals (auditable multi-dataset harmonization)

**`HarmonizeFeatures (idtrack/idtrack/_harmonize_features.py`**) operationalizes feature-space harmonization by applying an IDTrack mapping contract across multiple single-cell datasets represented as **`AnnData`** objects, converting per-dataset feature identifiers into a shared target release and target namespace, and emitting harmonized outputs together with diagnostics that make the transformation auditable rather than implicit. The workflow is intentionally oriented toward integration hygiene rather than toward biological claims, because the central scientific object in atlas-scale analysis is the feature-definition boundary, and that boundary is only reusable when the mapping layer preserves explicit loss and ambiguity semantics.

#### Core data structures and outputs

Harmonization operates on **`AnnData.var`** identifiers and produces (i) filtered and reordered matrices whose feature axis corresponds to the harmonized identifier set, (ii) per-dataset ledgers that record which identifiers failed conversion, produced multiple plausible targets, or collided onto the same target, and (iii) auxiliary dictionaries such as **multiple_*Ensembl*_dict** that preserve many-to-one collapses as explicit evidence rather than discarding it. Because these objects are structured (sets, dicts, and per-dataset tables) rather than log text, projects can archive them alongside datasets and can compare ledgers across runs to detect drift in the mapping layer, which is exactly the reproducibility posture the paper argues for.

#### Formalization (harmonization as a ledgered join over mapping outcomes)

Let datasets be indexed by *d ∈ {1,…, D}* with raw feature sets F_d_, and let m(·; 𝒞) be the IDTrack mapping operator that returns, for each *x ∈ F_d_*, an outcome triple (status(x),T (x), evidence(x)) consisting of an explicit status (no-corresponding, no-conversion, no-target, or mapped), a set of target identifiers *T (x) ⊆ 𝒰* in the requested **`final_database`**, and an optional audit payload. The harmonization objective is then to construct a shared target feature set *F ^⋆^ ⊆ 𝒰* under a declared policy (**`union`** or **`intersect`**) while emitting an explicit ledger Λ that records which x were removed due to mapping failure, ambiguity, cross-dataset inconsistency, or *n* → 1 collisions, so the transformation can be audited and reproduced as a structured artifact rather than inferred from downstream matrices.

#### Union versus intersection as an explicit policy

At the multi-dataset unification layer, **`unify_multiple_anndatas`** merges datasets either by preserving the union of all identifiers (outer join) while recording a derived **`intersection`** flag per feature (**`mode=‘union’`**), or by restricting to the strict intersection (features present in every dataset; **`mode=‘intersect’`**), which makes the choice between “maximum coverage” and “maximum comparability” a reportable analysis parameter rather than an implicit side effect of preprocessing scripts. This design is intentionally contract-aligned: when a harmonization result is used to train a model or to integrate cohorts, the resulting feature set is itself a scientific artifact, so the method forces the analyst to choose and report the reconciliation policy rather than letting it be determined implicitly by tool defaults.

#### Loss, ambiguity, and collisions as first-class diagnostics

No-mapping outcomes are tracked explicitly as removal sets (e.g., **`removed_conversion_failed_identifiers`**) or, when appropriate, as retained sets when failure is consistent across datasets (**`kept_conversion_failed_identifiers`**), multiple plausible mappings contribute to ambiguity diagnostics rather than being silently collapsed, and cross-dataset inconsistencies and n→1 collisions are surfaced as reportable issues (**`removed_inconsistent_identifier_matching`**, **`removed_final_database_duplicates`**). In contract terms, this is the point where “identifier mapping” becomes operationally reproducible at atlas scale: the same outcome semantics used by **`API.convert_identifier`** are preserved while constructing a shared feature space, so the mapping layer remains inspectable even when applied across many datasets.

#### Complexity

The dominant computational cost is converting F_d_ features per dataset (linear in feature count times per-identifier conversion cost) plus sparse matrix subsetting and alignment operations whose cost is proportional to the number of nonzero entries retained, so the method scales transparently with dataset size while keeping mapping semantics unchanged. Because harmonization is built as a deterministic ledger over a fixed snapshot graph, its computational scaling is separable from endpoint drift, which is why it serves as a credible integration primitive rather than as an ad hoc preprocessing step.

### 6.13. Source inference (when metadata are missing)

Source inference exists because real-world datasets frequently arrive with incomplete provenance, and conversion quality depends on knowing the source coordinate (release/assembly/database), so IDTrack provides **`API.infer_identifier_source`** as a helper that resolves each input to a canonical node where possible, consults **`TheGraph_node_trios`**, and tallies origin candidates via **`Track.identify_source`** (**`idtrack/idtrack/ _api.py`** and **`idtrack/idtrack/ _track.py`**). The API supports multiple reporting modes (**complete, assembly, *Ensembl*e_release, assembly_*Ensembl* _release**) and can return either the single most frequent origin or the full ranked list (**`report_only_winner=False`**), which is important for mixed identifier lists where a single “winner” may be scientifically misleading.

#### Algorithm (frequency vote over explicit trios)

Mechanistically, each resolved node contributes a set of possible origin trios (**`database, assembly, release`**) describing where it is active (**`TheGraph.node_trios`**, derived from combined-edge metadata and active ranges), and **`Track.identify_source`** performs a frequency count (via **`Counter.most_common`**) over the requested projection of these trios, returning the ranked distribution rather than an opaque score. This is intentionally not a learned model: the goal is to expose plausible provenance hypotheses as explicit counts that can be validated or reported with uncertainty, which is the scientifically honest posture when metadata are missing. In formal terms, for a projection operator *π (·)* (e.g. dropping database or release), the inferred source is the maximizer of the vote count 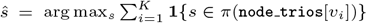, and returning the full ranked list corresponds to reporting the empirical distribution over s rather than collapsing uncertainty into a single point estimate.

#### Complexity and scientific use

Given a list of K identifiers, the tally is 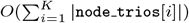 after normalization, and because `node_trios` is cached this reduces to set unions and integer counting, making the method fast enough to be used interactively to select snapshot parameters before running large conversions. Scientifically, inferred origins should be treated as hypotheses rather than as ground truth, and when used they should be reported explicitly (including mixed distributions) so the mapping contract remains an auditable object rather than a hidden assumption.

### 6.14. ConnectionBridge (HPC operation without direct internet access)

ConnectionBridge provides an operational pattern for environments where compute nodes cannot reach external endpoints directly, enabling IDTrack to run on restricted servers by routing the current *Python* process through a SOCKS5 proxy exposed via an SSH reverse tunnel (e.g., **ssh -R 1080 user@server**) so that *Ensembl* REST /HTTPS/FTP requests issued by that process can succeed during snapshot construction (**`idtrack/idtrack/ _connection_bridge.py`**). The bridge is intentionally process-scoped rather than system-scoped: enabling it monkeypatches **`socket.socket`** to a SOCKS-aware socket and can set **`ALL PROXY/all_proxy`** so subprocesses inherit proxy configuration, while stop() restores prior state, making it an operational enabler that does not alter mapping semantics once the snapshot artifacts exist.

#### Contract implication

ConnectionBridge is not part of the scientific method per se, but it reduces friction in the infrastructure settings where atlas-scale integration is executed, and it reinforces the contract framing by making it easier to build the same snapshot artifacts under constrained environments rather than motivating ad hoc, endpoint-dependent workarounds. Under the contract model, once the graph snapshot and allowlist configuration are built and cached locally, conversions can be rerun without live network access, and the provenance that must be published remains the same independent of whether ConnectionBridge was used to enable acquisition.

### 6.15. Cross-species humanization utilities (advanced, non-core)

IDTrack includes advanced utilities for cross-species workflows that combine within-species conversion on the *Ensembl* backbone with orthology resources and optional external mapper backends, enabling “humanization” style mapping from model-organism identifiers into human-target spaces (tutorials Part 6 and Part 7; **`idtrack/idtrack/ _external_mappers/`**). These utilities are explicitly non-core for the main manuscript Results and are positioned as workflow extensions for comparative analysis preparation rather than as central evidence for the drift/contract claims, because cross-species mapping introduces additional ambiguity semantics that are orthogonal to the within-species time-axis contract.

#### Expanded contract surface (what must be reported)

The contract principle still applies, but the surface becomes larger: in addition to snapshot boundary, assembly, allowlist identity, and ambiguity policy, cross-species workflows must report which orthology resource and version were used and how one-to-many orthology relationships were handled, because orthology definitions evolve and can introduce their own ambiguity semantics that must not be conflated with identifier drift. IDTrack’s role is to make the identifier layer explicit before and after orthology steps, so that a comparative pipeline can report a reconstructible sequence of coordinate transformations rather than a single opaque cross-species mapping table.

#### Implementation notes

Ortholog retrieval can be performed via *gget*-backed *Bgee* queries and, when enabled, sequence-alignment features can be computed via *BioPython* (**`idtrack/idtrack/ _external_mappers/_ortholog.py`**), meaning that cross-species pipelines must report not only which mapping contract was used on each within-species leg, but also which orthology endpoint and alignment feature set was used, because those components have their own versioning and ambiguity semantics (30; 46; 47).

## 7. Provenance Ledger and Reporting Template

### 7.1. Minimal provenance tuple (what must be reported)

To make an IDTrack-based mapping claim reconstructible, we recommend reporting a minimal provenance tuple that separates what is fixed by the snapshot graph from what is chosen at query time, because drift turns both layers into methodological coordinates.

#### Graph snapshot identity

At minimum, report IDTrack version (and commit hash when available), organism, snapshot boundary (**`snapshot_release`**) and historical window start (**`ignore_before`** when changed from defaults), and the primary genome assembly, because these fields define the time-axis interval and assembly context of the graph. If non-default graph-build settings were used (e.g., included forms, **`narrow`** mode, **`narrow_external=False`**, or modified external search limits), report them explicitly, because they change which nodes and edges exist in the snapshot and therefore what the pathfinder can consider reachable. Because snapshot filenames encode organism and release window but not the primary assembly, reporting the assembly explicitly (and, when possible, preserving the graph header fields such as ***Ensembl*_release**, **`genome_assembly`**, and **`version_info`**) makes the snapshot artifact self-describing even when copied out of its original cache directory.

#### External inclusion policy

Report the external configuration identity (filename) and, ideally, a content hash, because enabling or disabling particular namespaces changes both temporal reachability (via bridges) and cross-namespace ambiguity, meaning the policy is part of the scientific contract rather than a cosmetic preference.

#### Query-time choices and uncertainty semantics

Report the conversion target release (**`to_release`**) and target database (or the *Ensembl* backbone), the ambiguity strategy (**`strategy=‘best’`** vs **`strategy=‘all’`**), and whether bounded external bridging and explainability were enabled, because these choices determine whether multiple plausible targets are retained, whether external re-entry points are admissible, and whether audit trails are produced. When harmonizing datasets, also report the feature-space policy (union vs intersection) and how ambiguous and colliding targets were handled, because these decisions change the final feature definition that downstream models inherit. If non-default pathfinder bounds were used (e.g., synonym depth limits, external jump budgets, or permissive multi-transition stages), report them explicitly, because they change reachability and therefore change which failures are interpreted as “no mapping” versus which are interpreted as ambiguity under a fixed snapshot.

#### Verifiable artifacts

When possible, report concrete artifact identifiers that make the contract verifiable rather than merely described, including the graph snapshot filename produced by **`GraphMaker.create_file_name`** (which encodes the release window and build settings) and a hash of the allowlist configuration file content. If a full cache directory cannot be archived, publishing hashes of the contract-defining files still provides a reproducibility handle that allows reviewers and readers to detect silent drift in the mapping layer.

#### Unknowns and moving coordinates

If any element is unknown (e.g., the exact commit hash in a deployed environment), the scientifically safer practice is to report the uncertainty explicitly rather than to substitute informal labels such as “latest”, because under drift those labels are moving coordinates and therefore undermine the contract framing that this manuscript advocates.

### 7.2. Copy/paste reporting snippet

The following reporting template can be used verbatim in Methods sections (fill values, keep structure stable). Publishing the allowlist configuration file (or at least its hash and retrieval path) should be treated as part of the contract rather than a convenience.

~~~
We used IDTrack (version: <idtrack_version>, commit:
→ <git_commit_if_known>) to build a snapshot-bounded
→ identifier graph for <organism> with
→ snapshot_release=<snapshot_release>
→ (ignore_before=<ignore_before>), primary genome
→ assembly=<assembly>, and
→ allowlist_config=<allowlist_path_or_name> (hash:
→ <allowlist_hash_if_known>), yielding
→ graph_snapshot=<graph_snapshot_filename> (hash:
→ <graph_snapshot_hash_if_known>).
We converted identifiers to target_release=<to_release>
→ with final_database=<final_database_or_none> under
→ ambiguity policy strategy=<best|all>, with
→ external_bridging=<on|off>, explainability=<on|off>,
→ and optional pathfinder_bounds=<default|custom>
→ (synonym_depth=<…>, external_jump_budget=<…>).
We reported, per input, whether the outcome was
→ 1$\rightarrow$0 (no mapping), 1$\rightarrow$1 (unique
→ mapping), or 1$\rightarrow$n (multiple plausible
→ mappings), and we archived the built artifacts
→ (\Tool{HDF5} cache, allowlist, and graph snapshot) as
→ the contract-defining substrate.
~~~

In manuscripts with tight Methods space, this snippet can be shortened by omitting optional fields (e.g., **`ignore_before`** when default) while keeping the boundary-defining fields (snapshot boundary, assembly, allowlist identity, and ambiguity strategy), because those are the minimal coordinates needed to interpret mapping outcomes under drift.

---

**Fig. S1.**
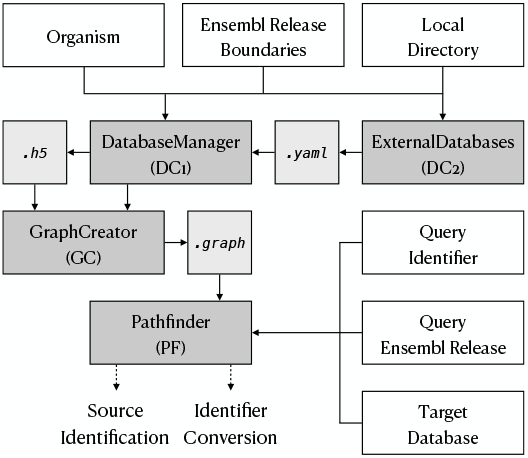
Workflow and artifact boundary in IDTrack (build once, query many). Build-time contract coordinates (organism, snapshot window, assembly context, and an external allowlist) drive database acquisition and caching and materialize a frozen identifier graph as a portable artifact whose identity can be archived and cited. Query-time tasks—source inference, identifier conversion, and dataset harmonization—then execute as contract-constrained traversals on this frozen snapshot through the pathfinder, returning typed outcomes and optional audit paths under the same declared policy.

